# Convolvulaceae of Guinea: taxonomy, conservation and useful plants

**DOI:** 10.1101/2024.07.15.602708

**Authors:** Becca Davis, Charlotte Couch, Ehoarn Bidault, Faya Julien Simbiano, Denise Moumou, Rafael Felipe Almeida, Ana Rita G. Simões

**Affiliations:** Royal Botanic Gardens, Kew, Richmond, Surrey, UK; Queen Mary University of London, London, UK; Missouri Botanical Garden, Saint Louis, USA; Muséum National d’Histoire Naturelle, Paris, France; Herbier National de Guinée, Université Gamel Abdel Nasser, Conakry, Guinea; Systematic and Evolutionary Botany Lab, Ghent University, Ghent, Belgium; East African Herbarium, National Museum of Kenya, Nairobi, Kenya

**Keywords:** climbers, morning glories, Red list, West Africa

## Abstract

Convolvulaceae is a diverse and economically important plant family in Tropical Africa, including the crop sweet potato (*Ipomoea batatas* (L.) Lam.) and its wild relatives, morning glories (*Ipomoea* L.), bindweeds (*Convolvulus* L.), invasive weeds (e.g. *Cuscuta* L.) and several other species with food, medicinal, or traditional uses. A taxonomic treatment of Convolvulaceae for Guinea is here presented, including comprehensive morphological descriptions, identification keys to genera and species, distribution and ecological information, specimen data, and documentation of known uses. A total of 51 species belonging to 16 genera are documented, 38 of which are native, and 13 presumed introduced; 33 of these are used as medicine, 18 as ornamental, 12 as food and 15 with a range of other uses. Preliminary IUCN Red List assessments were carried out for 45 species for which there were no previous assessments All 45 preliminary conservation assessments were categorised as Least Concern for the global Red List, with 61% of native species having at least one occurrence point falling within protected areas.

## INTRODUCTION

Guinea, West Africa, covers 245,857 km^2^, with a population who rely on agriculture and open-cast mining for income (Couch *et al*. 2020). The country is covered by five ecoregions, the largest of which are West Sudanian savannah and Guinean forest-savannah (Atsri *et al*. 2018; Brugière 2012). Guinean mangrove forests reach along the coast, and the southern regions contain the most threatened ecoregion, fragments of Western Guinean lowland forests (Atsri *et al*. 2018). The last ecoregion, Guinean montane forest, occurs in the Guinea Highlands of Fouta Djallon or Loma-Man Highlands (Couch *et al*. 2017). These flat topped plateaus reach 1,700 m (White 1983) and contain niche habitats including sandstone cliffs and high elevation (montane and submontane) *bowal* grassland. The unique habitats, particularly the high altitude and infrequent disturbance, give rise to the rich diversity of flora and fauna in Guinea, and some of the highest levels of plant endemism in West Africa, estimated at 4.7% (Sosef *et al*. 2017; Marshall *et al*. 2022; Gosline *et al*. 2023).

However, this rich flora has been reduced and fragmented due to habitat loss from agricultural and mining practices, with up to 96% of Guinea’s original forest having been lost (Sayer *et al*. 1992). In the past, wildlife conservation was not a high priority for Guinea, which resulted in few field surveys and insufficient data for all taxa, but particularly for plants (Brugière & Kormos 2009). Historically, plant exploration of Guinea has occurred in waves of floristic surveys, with a concentration of collections and studies during the colonial period, in the 1960s and 1970s, which paused and picked up more recently, in the 2000s, with a contribution of environmental surveys mandated by international mining companies. Previous conservation efforts have focused on the creation of national parks for preserving large game mammals, classified forests for timber exploitation and Ramsar wetland sites. However, these areas lack funding, management and up to date legislation, so the habitats within continue to degrade (Brugière & Kormos 2009). Crucially, the majority of Guinea’s threatened habitats and species fall outside of these sites, meaning the rich biodiversity lacks protection, and the need for increased resources and knowledge to prioritise areas for conservation remains high (Brugière & Kormos 2009; Couch *et al*. 2019).

Royal Botanic Gardens Kew (RBG Kew) have worked in Guinea since 2005, facilitating advances in botanical knowledge and conservation. A national herbarium, and later, a seed bank, were set up in Conakry to allow identification and storage of rare species (Cheek *et al*. 2018). In 2015, RBG Kew established the Tropical Important Plant Areas (TIPAs) programme, building upon the existing Important Plant Areas framework whilst taking into account the differences associated with the tropics: higher plant richness, higher socio-economic dependence on native plants, and less availability of data (RBG Kew TIPAs). The 3 criteria include: a) presence of threatened plant species, b) presence of high botanical richness and c) presence of threatened habitats (Darbyshire *et al*. 2017). These criteria enable the designation of TIPA’s which are used to identify areas to prioritise for conservation. RBG Kew works with many international partners in these countries so that the data created from TIPAs can feed directly into the management and protection of plants (RBG Kew TIPAs). The TIPAs project was set up in Guinea in 2015, the first in Africa, to use threatened species data to identify site-based areas for conservation priority (Couch *et al*. 2019). The resulting field surveys made large advances in Guinea’s commitments to document and protect its biodiversity (RBG Kew 2019). The outputs included 9 threatened habitats, 273 threatened plant species, and the designation of 22 TIPA sites (Couch *et al*. 2019). Of the 273 identified threatened plant species, a pilot project developed conservation action plans for 20 species: however, the creation of these 20 plans took 1,200 hours over 9 months, meaning it would take years to create plans for the remaining 253 threatened species (Couch *et al*. 2022). Therefore, local area-based conservation may be beneficial to effectively preserve the populations of threatened species through a more efficient use of time and resources (Couch *et al*. 2022). In response to a lack of plant data in the new management plans for Mt Bero and Diécké Classified Forests, two site-based conservation action plans were developed in 2022 (Diaby *et al*. 2022; Couch *et al*. 2022 unpublished). Additionally, a multispecies national action plan was developed for threatened trees of Guinea to address the policy requirements needed for their conservation and was submitted to the government in 2023 (Couch, Magassouba, & Kante 2023; Couch *et al*. 2023).

Alongside this, work is being done to increase engagement from local citizens in the conservation actions, so they can learn about their unique biodiversity and play a key role in its protection. Recently, Kew, in partnership with HNG (Herbier Nationale de Guinée - Université Gamal Abdel Nasser de Conakry) and Guinean NGOs, have created village tree nurseries to propagate and plant out threatened trees in the buffer zones of five TIPAs, created a teacher’s booklet to introduce plant conservation into school curricula, trained university students and forestry officers on how to ellaborate plant surveys and collect specimens, and ran a Campaign for a National Flower (Larridon & Couch 2016; Cheek *et al*. 2018). Despite these efforts, there is still much to be done to conserve Guinea’s plant diversity.

Floristically, the country is fairly well documented, but with great need for further work. Whilst the Flore de la Guinée was released in 2009, a 30 year delay in publication meant it was already out of date by the time it was published (Lisowski 2009). Meanwhile, the increase in field surveys resulting from internationally funded projects, and digitisation and georeferencing of herbarium specimens led to a large increase in botanical data and the preparation of the Checklist of Vascular Plants of the Republic of Guinea (CVPRG) (Couch *et al*. 2020; Gosline *et al*. 2023), which. reports 3,505 vascular plant species occurring naturally in Guinea, and 396 non-native plants (Gosline *et al*. 2023).

One of the 20 most taxonomically diverse plant families identified in the CVPRG is Convolvulaceae, commonly known as the ‘bindweeds’ or ‘morning glories’. Containing over 2,000 species across 61 genera, it has a worldwide distribution but is primarily found in the tropics in areas of secondary vegetation (Mitchell *et al*. 2016; Simões *et al*. 2022). They occupy a wide variety of habitats, from roadsides and hedges to forests or beaches, and are particularly common in forest transition zones. Globally, the family holds importance for humans due to the inclusion of sweet potato *Ipomoea batatas* (L.) Lam., one of the seven major food crops grown in all subtropical and tropical regions worldwide which yields 131 million tons per year (Sapakhova *et al*. 2023). Convolvulaceae are also grown ornamentally worldwide due to their attractive large sized and bright-coloured flowers and also have a wide range of medicinal and traditional uses, such as the pharmacological properties of *Ipomoea asarifolia* (Desr.) Roem. & Schult. leaves, which provide diuretic and purgative effects in West Africa (Khaled *et al*. 2017), and the seeds of many *Ipomoea* L. species which have hallucinogenic effects, due to high content of ergot alkaloids, often used for spiritual rituals in Africa (Chen *et al*. 2018). Convolvulaceae are usually herbaceous climbers with twining or prostrate stems, but can be trees, shrubs, or erect or prostrate herbs (Simões *et al*. 2022). They usually have funnel shaped flowers with five fused petals and conspicuous midpetaline bands, as well as alternate leaves which vary widely in size and shape; the genus *Cuscuta* L. does not follow these typical characteristics, but instead is a parasitic climber with a filiform stem, leaves reduced to scales and small flowers (Simões *et al*. 2022).

Within Guinea, 58 taxa of Convolvulaceae have been listed in the CVPRG (Gosline *et al*. 2023); however, this work provides minimal information about species. The present work aims to provide a taxonomic treatment for the Convolvulaceae of Guinea, building on the preliminary list presented in CVPRG, by doing further taxonomic and nomenclatural verifications, expanding the analyses of herbarium specimens, and presenting complete morphological descriptions and an identification key to help with the identification of future collections. Three species of *Cuscuta* have been documented (Gosline *et al*. 2023); however, issues were encountered which required further investigation, and for this reason, this genus is here excluded and will be treated in a separate work. The conservation status of each species is here assessed to identify any threatened species and occurrences within Guinea’s protected area network. This work will also assist in meeting Guinea’s Convention on Biological Diversity (CBD) targets by providing identification and knowledge of Guinea’s biodiversity, and the taxonomic and conservation data generated will contribute to the ongoing TIPAs programme in Guinea.

## MATERIAL AND METHODS

### Literature search

This study used the Checklist of Vascular Plants of Guinea (Gosline *et al*. 2023) as a starting point for listing the native and introduced Convolvulaceae species occurring in Guinea. The checklist has at least one expert-validated collection based record for each species, and includes all species recorded in the Flore de Guinée (Lisowski 2009). It records 44 native Convolvulaceae and 14 introduced or cultivated species, in a total of 58 species. A further comprehensive literature review was carried out to collect information on the morphological characteristics of each species. Floras from Africa were used as much as possible to account for potential variation seen across the large ranges of widespread species. The Flore d’Afrique Centrale (Mwanga Mwanga *et al*. 2022) was used preferentially as it contains up to date descriptions from nearby countries, but. other floras (Breteler 2015; Meeuse & Welman 2000; Verdcourt *et al*. 1963; Deroin 2001; Lisowski 2009; Oliver 1905; Pope & Launert 1987) were also used to create the first drafts of the species’ morphological descriptions, as well as a range of scientific articles (Acevedo-Rodríguez 2005; Austin 2008a; Breteler 2010; Breteler 2013; Lewis 1986; Lakshminarayana & Raju 2017; Lejoly & Lisowski 1992; Myint & Ward 1968; Priyashree *et al*. 2010; Wood *et al*. 2020). The characters’ information was collated into a morphological data matrix in Microsoft Excel, containing details of the habit, stems, leaves, inflorescence, flowers, fruits and seeds for each species.

### Morphological analyses

Herbarium specimens of Convolvulaceae from Guinea, were analysed, physically and digitally, from nine herbaria (BR, BRLU, HNG, K, MO, P, POZG, SERG, WAG) representing 51 species. For each specimen, 2D shapes e.g. leaf lamina, base and apex, were examined, alongside indumentum, margins, venation and all relevant measurements e.g. stem diameter and corolla length were taken. A preliminary morphological data matrix based on literature review was updated to reflect the variation among the Guinean specimens, and extra data was added as necessary. When available, flowers were rehydrated using a 1% solution of Libsorb (wetting agent), and dissected under the stereomicroscope to allow examination of the floral structures. Due to the limited number of available specimens, and lack of floral material on those available, flowers from only 5 species were rehydrated and dissected, with literature being relied up on for the remaining species. Information regarding the habit, flower colour and fruit was noted alongside altitude, collection location, uses and additional ecological data, retrieved from herbarium labels.

### Specimen databasing

Specimen data for all species in Guinea were downloaded from the Global Biodiversity Information Facility (GBIF 2023) and online herbaria databases (P, BR, WAG, MO), taxonomically verified by analyses of online images of the specimens, and complemented with data extracted from specimens analysed in person at K and HNG herbaria.These were compiled into a database, collecting information on collector, collector number, collection date, herbaria barcode, locality information and coordinates if available.

### Georeferencing and mapping

Specimens lacking coordinates were georeferenced from location descriptions where possible using Google Earth Pro Version 7.3 (2022). The data was recorded as decimal latitude and longitude, alongside the level of accuracy. The preserved specimen and human observation coordinates were cleaned using RStudio Version 06.0+421 (Rstudio 2020) CoordinateCleaner package (Zizka *et al*. 2019) to remove duplicate occurrences and those falling into botanical gardens, into the sea, or outside of Guinea.

### Conservation assessments

Only 9 of the Convolvulaceae species occurring in Guinea had previously been assessed for the IUCN Red List of Threatened Species (IUCN, 2022). Therefore, 45 preliminary Red List assessments were carried out for the remaining species. The majority of the species are distributed across multiple countries and continents, therefore occurrence data with coordinates from GBIF (2023) were used directly for the assessments. For the few species with narrower distributions and fewer than 100 occurrence points, all GBIF occurrences without coordinates were georeferenced, and further resources were used for additional occurrence data. GeoCAT (Bachman *et al*. 2011) was used to calculate the Area of Occupancy (AOO) and Extent of Occurrence (EOO) for each species, and to overlay their global distribution with protected areas. The species were assessed using the IUCN Red List criteria B regarding the geographic range based on the EOO and AOO, due to the lack of data to support the other criteria.

### Taxonomic treatment

The taxonomic treatment was compiled using the *mailings* feature on Microsoft Word. The morphological descriptions were computed from the morphological data matrix. Information regarding species names, authors, basonyms and synonyms were collated from Plants Of the World Online (POWO 2024) and International Plants Names Index (IPNI 2024). Details regarding type specimens and worldwide distributions were sourced from floras, taxonomic and nomenclatural databases (TROPICOS, JSTOR; POWO, IPNI, the African Plant Database) and other specialised online literature databases (e.g. BHL). Habitat and altitude data for the species within Guinea were retrieved from specimen label information, alongside an overview of the species locations in Guinea down to province level. All specimens seen, both in person and online, are listed. An identification key to the genera was created, using the key to the Convolvulaceae genera from the Flore d’Afrique Centrale (Mwanga Mwanga *et al*. 2022) as a template; species level identification keys were generated specifically for this study.

## TAXONOMY

### Key to the genera of Convolvulaceae in Guinea

**1a.** Prostrate herb, sometimes stems ascending; flowers blue or yellow, never purple; corolla small, infundibuliform (never campanulate) less than 15 mm long 2

**1b.** Climber, or suberect sub-erect herb or shrub; flowers white, yellow, red, blue or purple; corolla of variable size and shape, up to 12 cm long 4

**2a.** Flowers blue; styles 2 1. **Evolvulus**

**2b.** Flowers yellow to white; style 1 3

**3a.** Leaf oblong-linear to narrowly elliptical, sometimes pinnatipartite 2. **Xenostegia**

**3b.** Leaf ovate to ovate-reniform, entire to ± 3-lobed or with 2 to 3 angles, base deeply cordate. 3. **Merremia hederacea**

**4a.** Suberect shrubby herb; leaves sessile, leaf lamina ≤ 7mm long; corolla small, ≤ 8 mm long 4. **Cressa**

**4b.** Climber; leaves petiolate; leaf lamina longer than 7 mm; corolla larger than 8 mm long5.

**5a.** Flowers white; styles 2, entire or partially free 6

**5b.** Flowers white, yellow, red, blue or purple; style 1 8

**6a.** Bracteoles or sepals not accrescent in fruit 5. **Bonamia**

**6b.** Bracteoles or sepals accrescent in fruit 7.

**7a.** Bracteoles accrescent in fruit; many-flowered; corolla 3-10 mm long 6. **Neuropeltis**

**7b.** Sepals accrescent in fruit; 1-3-flowered; corolla 15-30 mm long 7. **Calycobolus**

**8a.** Sepals leaf like; stigmas oblong, elliptical, rarely globular or linear 9

**8b.** Sepals not leaf like; stigmas always globular 10

**9a.** Corolla pure white, without purple centre; leaf narrowly oblong-elliptical, rounded at the base; fruit ovoid 8. **Aniseia**

**9b.** Corolla yellow, or rarely white with a purple centre; leaf cordate or hastate, rarely truncated or cuneate at the base; fruit globular or subglobular 9. **Hewittia**

**10a.** Stems winged or strongly angular 3. **Merremia pterygocaulos**

**10b.** Stems cylindrical or slightly angular 11

**11a.** Pair of inconspicuous spiniform projections at the insertion of the petioles; flowers in congested umbelliform cymes, yellow, with a tuft of hairs at the apex of the midpetaline bands; 10. **Camonea**

**11b.** Absence of inconspicuous spiniform projections at the insertion of the petioles; flowers solitary or in few-many axillary cymes or capitate heads, glabrous or pubescent, variable in colour 12

**12a.** Leaf 5-7-lobed, palmatisect to palmatipartite; 13.

**12b.** Leaf entire 14.

**13a.** Flowers pure white or bright yellow; sepals lanceolate, apex acuminate 11. **Distimake**

**13b.** Flowers mauve or purple; sepals ovate, apex obtuse…………………………………………………. **Ipomoea** p.p. (see *I. mauritiana*, *I. cairica*)

**14a.** Calyx completely surrounding the fruit; corolla very bright crimson with base of tube orange-yellow 12. **Stictocardia**

**14b.** Calyx not or slightly accrescent, but never surrounding the fruit; corolla any other colour. 15

**15a.** Filaments with a scale at the base; corolla urceolate 13. **Lepistemon**

**15b.** Filaments often enlarged at the base, but with no scale; corolla infundibuliform or campanulate 16

**16a.** Woody plant, never herbaceous; leaves broadly cordate, lower surface densely silky with silvery white to tawny grey and shiny tomentose; fruit indehiscent, with a fleshy, ossified or ± woody pericarp 14. **Argyreia**

**16b.** Herbaceous or woody plant; leaves of variable shape, if lower surface densely silky with silvery white hairs, leaves not cordate in shape; fruit with a thin pericarp, opening with valves or by irregular dehiscence 15. **Ipomoea**

## 1. Evolvulus L

### 1.1. Evolvulus alsinoides (L.) L

*Convolvulus alsinoides* L. in Sp. Pl.: 157 (1753).

Type: Sri Lanka, Herb. Hermann 3: 55, No. 76 (lectotype: BM000628009).

Annual or perennial *herb*. *Stems* trailing or prostrate, not climbing, spreading or ascending; flowering shoots ascending, terete, up to 1 mm in diameter, up to 50 cm long, not rooting at the nodes, densely covered with long patent silky hairs, silvery or reddish brown. *Leaves* subsessile or shortly petioled; *petiole* 0.1-3 mm long; lamina entire, elliptic to linear-oblong, 8-26 mm long, 2-10 mm wide, attenuate to rounded at the base, obtuse to acute and distinctly mucronate at the apex, densely silky pilose on both surfaces, with long flat silvery white hairs; midrib protruding below, 2-3 pairs of secondary veins. *Inflorescence* axillary cymes, 1-5-flowered; peduncle 4-40 mm long, filiform, pilose; bracteoles narrowly elliptic-ovate, unequal length,4-8(−15) mm long, with ciliate margin, opposite. *Flower*: pedicel 2-10 mm long, filiform, spreading, pilose like peduncle; sepals subequal, ovate-lanceolate, elliptical-oblong to elliptical, apex acute to acuminate, 2-4(−5) mm long, 2 mm wide, margin ciliate, densely silky or villous, not accrescent in fruit, outer sepals longer than the inner; corolla broadly infundibuliform, 4-6 mm long and wide, blue, rarely white, the folds paler beneath, glabrous, weakly lobed or truncate at the apex, midpetaline bands pilose; stamens 5, included, glabrous, white, filaments ± 2 mm long, very slender, linear, glabrous, epipetalous, anthers ovoid, base deeply sagittate, dithecous, 0.8-1 mm long, white; pollen smooth, pantoporate;ovary disc annular, united with the lower half of the ovary wall; ovary globose, glabrous, green, 2-celled, 2-ovuled per cell; styles 2, 3 mm long, clavate, united only at the base, glabrous, stigmas 4, ± 2 mm long, long, terete or subclavate, white. *Fruit*: capsule, globose, 3-4 mm long, glabrous, septum reduced to a membranous wing on the placenta, 4-valved. *Seed*s 4, ovoid, 1.7 mm long, pale brown-black, reddish, glabrous, smooth.

DISTRIBUTION – Widespread in the tropics and subtropics. In Guinea: Haute Guinée.

HABITAT – In Guinea: along the roadside, between 375-435 m (elsewhere up to 2400 m).

SPECIMENS EXAMINED – **Guinea**: Haute Guinée: Kankan Region: Kankan Préfecture, Ville Kankan, 17 Nov. 1966, *Lisowski 10634* (BR0000015986443!, POZG-V-0057205).

ADDITIONAL SPECIMENS – **Guinea**: Haute Guinée: Kankan Region: Kankan Préfecture, Kankan, 12 Jun. 1967, *Lisowski 10635* (POZG-V-0057204); Haute Guinée: Kankan Region: Kankan Préfecture, Kankan, 2 Jul. 1963, *Lisowski 90332* (POZG-V-0057215); Kouroussa, *Brossart 15750* (P), *loc. cit*., *Pobéguin 255* (P), *loc. cit*., *Pobéguin 291* (P).

CONSERVATION (PRELIMINARY ASSESSMENT)– Least Concern (LC), following the global IUCN (2012) guidelines; known from 14,761 occurrence points worldwide.

USES - *E. alsinoides* is a species with a wide range of uses: food, medicine, ornamental, materials, fodder, and other environmental, social and religious applications (POWO 2024). It is assumed to be originally from the Americas, and presumably introduced in other regions by Europeans due to its wide range of medicinal properties (Austin 2008b). In several countries of Africa and Asia, the very bitter leaves are used in the preparation of tonics and febrifuges. In Niger, a leaf decoction is taken as a laxative or purgative; in Togo, an infusion of the plant is taken to cure problems of menstruation; in Bénin, powdered leaves are applied to cure stiffness of the limbs; in Ethiopia, the leaves are ground and applied on burns in Kenya, the powder is applied to bleeding wounds; and, in Tanzania, the dried leaves are burned in a pipe against leprosy. Throughout West Africa, the plant is used as an amulet (*grigri*) to attract love or obtain favours. It is one of the plants known in traditional Indian Ayurvedic medicine and renowned for its memory strengthening properties, as well as antidepressant, anti-epileptic, aphrodisiac and immuno-modulatory actions. It is also an ingredient in preparations for fevers with indigestion or diarrhoea, as an astringent, or for treating internal bleeding; it is also febrifuge and tonic for its roots. In southern India, an infusion of the powdered entire plant is drunk against syphilis. A preparation of the plant in oil is used to stimulate hair growth. The leaves can be rolled into cigarettes and smoked in the case of chronic bronchitis and asthma (Mwanga Mwanga *et al*. 2022).

## 2. Xenostegia D.F. Austin & Staples

1a. Leaf lamina entire, base hastate or auriculate, lobes or auricles toothed **1. X. tridentata**

1b. Leaf lamina deeply pinnatifid, base not hastate or auriculate **2. X. pinnata**

2.1. *Xenostegia tridentata* (L.) D.F. Austin & Staples

*Convolvulus tridentatus* L. in Sp. Pl.: 157 (1753).

Type: Rheede tot Drakestein, Hortus malab., vol. 11: t. 65 (1692).

Perennial *herb*, glabrous or pubescent. *Stem* prostrate or twining, angular, not winged, slender in diameter, with a small woody root giving off several stems. *Leaf*: *petiole* 0-3 mm long; lamina entire, oblong-linear, 10-75 mm long, 2-12 mm wide, hastate or auriculate, auricles usually with 1-5 strongly acute teeth at the base, acute to obtuse, mucronate at the apex, upper surface slightly pubescent, particularly on the veins, to glabrous; midrib depressed above, projecting below, secondary veins numerous. *Inflorescence* axillary cymes, many-flowered; *peduncle* 2-6 cm long, cylindrical; *bracteoles* lanceolate to linear, apex acute, 1.5-2.5 mm long; *pedicel* 4-15 mm long, cylindrical; sepals unequal, ovate-oblong to ovate-elliptical, apex acuminate and mucronate, 4-5-(−10) mm long, 1-3 mm wide, pubescent, outer sepals slightly shorter, 4.5 mm long, 2 mm wide, inner sepals ± 4 mm long, 1.5 mm wide; corolla infundibuliform, 10-15 mm long, greenish, cream or sulphur yellow, dark throat, with 5 distinct lobes, midpetaline bands pubescent; stamens included, filaments equal, 3.5 mm long, grooved, pubescent with long hairs, or glabrous, inserted 2 mm from the bottom of the triangular base, anthers elongated, ± 1 mm long, straight at dehiscence; pollen smooth, pantoporate; ovary disc annular, distinctly 5 lobed, 0.4 mm high; ovary ovoid, 2.5 mm long, glabrous, 2-celled, 2-ovuled per cell; 1 style 4 mm long, stigma bilobed. *Fruit*: capsule ellipsoidal, ± 13 mm long, pericarp finely grained, reduced partition with 2 membranous wings, without frame or columella, surrounded by accrescent calyx at least 6.5 mm long, dehiscing in 4-valves, deciduous. *Seeds* 4, ovoid, 3-6 mm long, 2-3.5 mm wide, 2-3.5 mm high, black, glabrous, hilum encircled by a heart shaped rim.

DISTRIBUTION – Widespread in tropical and subtropical Africa, Asia and Australia. In Guinea: Guinée Maritime.

HABITAT – In Guinea: fallow land, forest-savannah, forest edge, roadside; found at 10-260 m (elsewhere up to 1,650 m).

SPECIMENS EXAMINED – **Guinea**: Guinée Maritime: Boké Region: Boké Préfecture, Sangaredi, Para Gogo, Feloparawol Aliou, 24 Nov. 2013, *Lopez Poveda 287* (HNG, K000749902!); Kabata, on route to Kabata, 24 Nov. 2017, *Camara 67* (HNG, K000683807!).

ADDITIONAL SPECIMENS – **Guinea:** Guinée Maritime: Conakry, 2 Jan. 1979, *Lisowski 51309* (POZG-V-0059222).

CONSERVATION STATUS (PRELIMINARY ASSESSMENT) – Least Concern (LC), following the global IUCN (2012) guidelines; known from 3,690 occurrence points worldwide.

USES-In Asia, all parts of the plant are considered purgative; roasted seeds are anthelmintic and diuretic; in Malaysia a poultice of the leaves is applied to the head against fever and against snakebites; in the Philippines and India a decoction of the roots is used as a mouthwash for toothache. In northern Nigeria, a decoction of the whole plant, together with *natron* (hydrated sodium carbonate), is taken to treat gonorrhoea (Mwanga Mwanga *et al*. 2022).

### 2.2. Xenostegia pinnata (Hochst. ex Choisy) A.R.Simões & Staples

*Ipomoea pinnata* Hochst. ex Choisy in A.P.de Candolle, Prodr. 9: 353 (1845).

Type: North Sudan, Darfur, Kordofan, Tejara, 19 Nov. 1839, *Kotschy 262* [lectotype G-DC (G00135552!); isolectotypes: BR (BR0000008885067!, BR0000008885395!), G (G00392708!, G00392706!), GH (GH00066982!), HBG (HBG505549!, HBG505550!), LD (LD1800417!), GOET (GOET005704!), K (K000097228!, K000097229!), M (M0109914!, M0109915!), MPU (MPU007053!), TCD, TUB (TUB005439!, TUB005440!), S (S11-38639!, S11-38640!), STU (STU000321!), WAG (WAG0000760!)].

Annual *herb*; main stem ± erect, short, branched at the base, twining or prostrate on the ground, emitting several ascending branches, villous with long hairs ± erect. *Leaf*: sessile; lamina 8-15-lobed, deeply pinnatisect, elliptic to elliptic-oblong, 1.5-5(−7.2) cm long, 1-2.5 cm wide, lobes linear segments, 8-15 pairs, 3-12 mm long, 0.5-1(−1.5) mm wide, the lower two segments dissected, stipuliform, covered with long slightly tawny hairs, sparse and spread out on both surfaces. *Inflorescence* axillary cymes, 1-3-flowered; peduncle 2.5-4 cm long, villous like the stems. *Flower*: pedicel 4-5 mm long; sepals unequal, elliptical, apex suddenly and largely acuminate, 4-8 mm long, 1.5-2 mm wide, villous on the outside, persistent in the fruit, outer sepals longer; corolla infundibuliform, 7-13 mm long, yellow to whitish, midpetaline bands pubescent; anthers straight at the dehiscence; pollen smooth, pantoporate; ovary globose, 1-1.5 mm in diameter, villous at the apex; style 1, 5-8 mm long, glabrous, stigma bi-globose. *Fruit*: capsule globose, 5-6 mm in diameter, villous above, 4-valved. *Seeds* 4, 2–2.8(–3) mm long, 2–3 wide, blackish, glabrous.

DISTRIBUTION – Widespread in tropical and southern Africa. In Guinea: Guinée Maritime, near the coast.

HABITAT – In Guinea: savannah, river edge; found at altitudes of 5-20 m (elsewhere up to 1500 m).

SPECIMENS EXAMINED – **Guinea**: Guinée Maritime: Boké Region: Boké Préfecture, Bel-Air, on the track to Drameya, 21 Oct. 2016, *Bidault 2485* (BRLU, HNG, K001753512!, MO-3062147, P01192527!, WAG).

ADDITIONAL SPECIMENS – **Guinea**: Guinée Maritime: Kindia Region: Forecariah Préfecture, Sengulen, *Molmou 690*, 3 Jul. 2012 (HNG0000654!); Ile Tristão, Nov. 1956, *Jacques-Félix 7329* (P03866968!); Conakry, *MacLaud s.n.* (P03866970!).

CONSERVATION STATUS (PRELIMINARY ASSESSMENT) – Least Concern (LC), following the global IUCN (2012) guidelines, known from 1,514 occurrence points worldwide.

## 3. Merremia Dennst. ex Hallier f

**1a.** Stem terete, not winged, leaf lamina margin entire to crenate or rarely 3-lobed, ovate in outline; corolla small, 0.6-1.2 cm long, campanulate, yellow, glabrous on the outside.………**. 1. M. hederacea**

**1b.** Stems angular, distinctly winged, leaf lamina 3-5-palmatilobed; corolla large, 3-5 cm long, infundibuliform, white or cream, throat red or purple inside, mid-petaline bands strigose, at least on the upper third **2. M. pterygocaulos**

### 3.1. *Merremia hederacea* (Burm.f.) Hallier f

*Evolvulus hederaceus* Burm.f. in Fl. Indica: 77 (1768).

Type: Indonésie, Java, *Pryon s.n.* [lectotype: G-DC (G00360097!)].

Perennial *herb*; stem twining or prostrate, angular, up to 1 mm in diameter, reddish-purple, ± 2 m, sometimes rooting at the nodes, often minutely tuberculate, glabrous or rarely weakly pubescent. *Leaf*: *petiole* 0.5-6 cm, slender, sparsely tuberculate, especially in the basal half, glabrous to very sparsely pubescent; lamina entire, crenate or shallowly to deeply 3-lobed, ovate, 1.5-5 cm long, 1.25-5 cm wide, cordate at the base, obtuse and mucronulate at the apex, membranous, glabrous or very sparsely pubescent with whitish hairs on both surfaces; 7 basal veins. *Inflorescence* axillary, lax cyme, few-flowered or flowers solitary; *peduncle* 0.5-10 cm long, thicker than the petioles, glabrous to glabrescent with whitish hairs; *bracteoles* oblong, apex mucronulate, 3 mm long, caducous; *pedicel* 2-4 mm long, sparsely pubescent with whitish hairs; *sepals* subequal, glabrous or sparsely pilose along the margins, 3 outer sepals broadly obovate, apex apiculate, 4-5 mm long, 2-3.3 mm wide, coriaceous, 2 inner sepals subtriangular, 3 lobes at the apex with 2 rounded ciliated lateral lobes, 5-6 mm long, 4-4.2 mm wide; corolla campanulate, 6-12 mm long, white or yellow, glabrous outside, pilose inside at the base, entire or slightly lobed; *stamens* included, filaments 4-6 mm long, flattened, not broadened at the base, anthers longitudinally dehiscent, not twisted after; *pollen* 3-colpate; *disc* annular; *ovary* ovoid, glabrous, 2-4-celled, 4-ovuled; style simple, filiform, glabrous, included; stigma bi-globose, warty. *Fruit*: capsule, globose or conic, somewhat 4-angled, 5-6 mm long, crowned at the base by persistent reflexed sepals, 1-4-celled, 4-valved or dehiscing regularly, valves reticulately wrinkled. *Seed*s 4, 2.5mm long, trigonal, pubescent or glabrous.

DISTRIBUTION – Widespread in tropical and subtropical Africa, Asia, Australia and Pacific Islands. In Guinea: Haute Guinée.

HABITAT – In Guinea: riverine bush, shrubland, gallery forest, river edge; found at 380-500m (elsewhere up to 500 m).

SPECIMENS EXAMINED – **Guinea**: Haute Guinée: Kankan Region: Kérouané Préfecture, Kérouané city, Bank of river Milo, 16 Nov. 2005, *Tchiengue 2408* (HNG, K000460110!, P04525709!); Siguiri Préfecture, Siguiri, bord de la rivière Tinkisso, 4 Dec. 1937, *Portères 2623* (P01183699!); 4km N. of Kankan, edge of River Milo, 2 Jan. 1967, *Lisowski 60927,* (BR0000017253567!, P, POZG-V-0059154).

ADDITIONAL SPECIMENS – **Guinea**: Haute Guinée: Kankan Region: Kankan Préfecture, 8km to the Northeast of Kankan, 29 Dec. 1966, *Lisowski 91034* (POZG-V-0059153); Kankan, between Kankan and Karifamoriya, 22 Dec. 1966, *Lisowski 91016* (POZG-V-0059155).

CONSERVATION STATUS (PRELIMINARY ASSESSMENT) – Least Concern (LC) following the global IUCN (2012) guidelines; known from 1121 occurrence points worldwide.

USES - In Asia, the seeds of the plant are used to treat fever, colds, sore throats, haematuria, conjunctivitis and boils. In Sudan, the long slender stems are used to tie loads to animal saddles. In several countries, the plant is also cultivated in gardens for its beautiful flowers (Mwanga Mwanga *et al*. 2022). In India, leaves of *M. hederacea* have been reported to be used in treating chapped hands and feet (Olatunji *et al*. 2021).

### 3.2. *Merremia pterygocaulos* (Choisy) Hallier f

*Ipomoea pterygocaulos* Choisy in A.P.de Candolle, Prodr. 9: 381 (1845).

Type: Erythrea, Tigray, Schire, Mai-Dogale, prope Mai Dogale, 13 Nov. 1839, *Schimper 630* [holotype G-DC (G00135790!); isotypes BM, BR (BR00000825066!, BR000000088547!, BR0000008885470!), G-DC (G00392716!, G00392717!), K (K000097233!, K000097234!), L (L0004242!, L0004243!), LG (LG0000090028229!), M (M0109912!), MO (MO-345997), MPU (MPU007054!), S (S11-38661!), STU (STU000453!), TUB (TUB005442!, TUB005443!, TUB005444!)].

Perennial *herb* or *shrub*, entirely glabrous with the exception of the base of the stem being very densely hirsute; stem twining or prostrate, conspicuously 4-winged, with wings up to 2 mm wide, ultimate branches slender, 3mm in diameter, often reddish, can reach several meters. *Leaf*: *petiole* 1-2(−7.5) cm long, narrowly or widely winged; lamina 3-7-lobed, palmatifid to palmatipartite, ovate-orbicular in outline, 5-8.5cm long, 6-15cm wide, cordate at the base, mucronate at the apex, lobes elliptic, ovate or oblong, ± mucronate, 3.2-9 cm long, 2.5-4 cm wide; venation depressed above, protruding below, 5 palmate basal veins, numerous secondary veins in each segment. *Inflorescence* axillary cymes, few-many-flowered, rarely 1-flowered; peduncle 1-10 cm long, winged; *bracteoles* narrowly elliptical-oval to narrowly oblong or linear, apex subulate, 2-8 mm long. *Flower: pedicel* 5-30 mm long, thickened and remaining erect in fruit, often conspicuously scarred; *sepals* subequal, oval, apex usually obtuse, 8-10 mm long, 4-8 mm wide, thinnish, chartaceous, accrescent, becoming broadly ovate to circular and ultimately spreading in fruit, imbricate, outer sepals broadly ovate, apex obtuse to apiculate, margin scarious, inner sepals oblong to orbicular; corolla infundibuliform or campanulate, 3.6-5 cm long, 4-5 cm wide, white or cream with red or purple throat, externally glabrous, entire or 5 lobed at apex, mid-petaline bands strigosely pilose in the upper third; *stamens* included, filaments equal, 11-18 mm long, triangular, grooved and pubescent at the base, inserted ± 4 mm from the bottom of the tube, anthers 3-4 mm long, spirally dehiscing; pollen 3-colpate; disc 0.8-1 mm high; ovary subglobose to pyriform, ± 2 mm in diameter, green, 2-4-celled, 4-ovuled, with voluminous cells; *style* simple, filiform, 15-20 mm long, included; *stigma* bi-globose. *Fruit*: capsule, ovoid-conical ± truncate, or flattened-depressed at the apex, 1.2-2 cm in diameter, brown, upper half of the pericarp fleshy, circumsessile, septum reduced to 2 membranous and triangular wings on the placenta, crowned with the persistent style-base, 1-4-celled, opening with 4 valves, dehiscing regularly. *Seed*s 4, subglobose, 8mm long, 5mm wide, black, glabrescent, smooth.

DISTRIBUTION – Widespread in tropical and southern Africa. In Guinea: across all four provinces.

HABITAT – In Guinea: secondary vegetation, submontane forest, roadside, river edge; found at 120-1279m (elsewhere up to 2200m).

SPECIMENS EXAMINED – **Guinea**: Guinée Forestière: Nzérékoré Region: Macenta + Beyla Préfecture, Simandou Range, Canga East, 10 May 2009, *Haba 578* (HNG, K000683677!); Simandou Range, 500m S. of Oueleba, 6 Jul. 2006, *Cheek 73* (K000437763!).

ADDITIONAL SPECIMENS – **Guinea**: Guinée Maritime: Kindia Region: Forécariah Prefecture: Singuelen Village, 6 May 2012, *Sow 864* (HNG0000827!); Haute Guinée: Kankan Region: Kankan Préfecture, near the village Sadia, S. of Kankan, 23 Jan. 1967, *Lisowski 10625* (BR0000017256674!, POZG-V-0059199, POZG-V-0059200); Moyenne Guinée: Mamou Region, Mamou Préfecture, Kaba, *Chevalier 20390,* 10 Jan. 1909 (P00434226!, P00434227!, P00434228!); Entre Kaba et Toufili, 11 Jan. 1909, *Chevalier 20397* (P00434229!, P00434230!, P00434231!, P00434241!, P00434242!, P00434243!).

CONSERVATION STATUS (PRELIMINARY ASSESSMENT) – Least Concern (LC) following the global IUCN (2012) guidelines; known from 273 occurrence points worldwide.

## 4. Cressa L

### 4.1. Cressa cretica *L*

Type: Créte, Herb. Linné 317.1 [lectotype: LINN (LINN-HL317-1!)].

Perennial shrubby *herb*; *stems* woody at base, spreading or ascending, terete, ±1 mm in diameter, caniculate, older stems glabrous, younger stems pilose with long simple whitish hairs, with densely-leaved branchlets. *Leaf*: sessile; lamina entire, ovate-lanceolate to ovate, 2-7mm long, 1-4mm wide, cuneate, rounded or even subcordate at the base, acute at the apex, grey-green and adpressed pubescent on both surfaces with long simple whitish hairs. *Inflorescence* axillary, 1-flowered, in spikes to head-like clusters, aggregate at the tips of the branchlets; sessile or shortly pedunculate, < 1mm long; *bracteoles* mostly linear, unequal in length, 2-3mm long, 1mm wide. *Flower*: sessile; *sepals* subequal, ovate-obovate, herbaceous, coriaceous, silky pubescent, imbricate, outer sepals ovate, apex acute to obtuse, 3-4mm long, 1.5-2mm wide, inner sepals ovate, apex acute, 2.5-3mm long, 1-1.5mm wide; corolla salver shaped, tube enveloped by calyx, 5-8 mm long, tubes and lobes about equal, white with pink flush, lobes 5, ovate, imbricate, spreading, pilose on the outside *stamens* exserted from the tube for 2.5-3.5 mm, *filaments* equal, filiform, glandular-pubescent at base, *anthers* oblong; ovary ovoid, hirsute at the apex, 2-celled, 4-ovuled; styles 2, unequal, distinct to the base, exserted, 2 stigma, large, capitate. *Fruit*; capsule ovoid, pilose at the apex, 1-celled, 2-4-valved, usually 1-seeded.*Seed* 1, ovoid, 3-4mm long, dark brown, glabrous, shining to reticulate.

SPECIMENS EXAMINED – **Guinea**: Guinée Maritime: near Conakry, 20 Sept. 1897, *Maclaud 31* (BR0000018872606!, K001759755!).

DISTRIBUTION – Widespread in Africa; Meditterranean; west, central and southeast Asia; Australia, and South America. In Guinea, only one occurrence was found, near Conakry.

HABITAT – Found at altitudes of 0-175 m, globally.

CONSERVATION STATUS – LC (Least Concern) on the IUCN Red List (Ghogue & Lansdown 2020).

USES – *C. cretica* has medicinal uses as is antibilious, antitubercular, expectorant, anthelmintic so can be used to enrich blood and treat constipation, leprosy, asthma, diabetes, urinary problems and general weakness. In Sudan, the leaves are crushed and mixed with sugar to induce vomiting. The fruits have potential to be a source of edible oil (Ghogue & Lansdown 2020).

## 5. Bonamia Thouars

### 5.1. *Bonamia thunbergiana* (Roem. & Schult.) F.N.Williams

*Convolvulus thunbergianus* Roem. & Schult. in Syst. Veg., ed. 15[bis]. 4: 884 (1819). Type: Sierra Leone, *Don s.n.* (BM000930510!).

Perennial woody *liana*; *stem* twining, terete, 1-3mm thick in diameter, reaching 4m in height, covered in small black dots, pubescent with simple yellowish hairs, densely whilst young, becoming glabrous in age. *Leaf*: *petiole* 5-15mm long, with simple yellowish hairs; lamina entire, oblong to oblong-lanceolate, 2-8.5cm long, 1-3.5cm wide, rounded at the base, obtuse-mucronulate, rarely acute or acuminate at the apex, subcoriaceous or membranous, upper surface green and thinly puberulous, lower surface densely pubescent with golden brown hairs; 6-10 pairs of secondary veins, pinnate, sunken above, prominent beneath. *Inflorescence:* dense cymes, usually secund on short peduncles or congested into a terminal panicle, many-flowered; *peduncle* 5-15mm long, tomentose with golden brown hairs; bracteoles lanceolate, minute. *Flower*: *pedicel* 4-15 mm long, similar in indumentum to the peduncles; *sepals* unequal, oblong-lanceolate to ovate, apex acuminate or acute, ± 6-8mm long, not accresscent but persistent in fruit, outer sepals sparsely puberulous with simple yellowish hairs, inner sepals slightly shorter, coriaceous to glumaceous, densely silky tomentose on the outside; corolla cupuliform, obscurely lobed or subentire, 1.5-2.5cm long, white, pilose or hirsute externally on the lobes, glabrous on the midpetaline bands; *stamens* included, *filaments* unequal, filiform, broadened at the base, glabrous above, pilose along the edge near the base, *anthers* oblong, base cordate, dorsifixed; *disc* at the base of the ovary; *ovary* ovoid, pilose near the apex, 2-celled, 4-ovuled, with axillary placentation; styles 2, bifid above the middle, with scattered long hairs, *stigma* conical, rugose. *Fruit*: *capsule* ovoid, apiculate, ± 5-8mm long, glabrous, 2-celled, 8-valved, 2-4-seeded, dehiscing from the base in several clefts. *Seeds* 2-4, ovate-oblong, black, glabrous.

DISTRIBUTION – West and central tropical Africa. In Guinea: mostly in Guinée Forestière, a few occurrences in Guinée Maritime.

HABITAT – In Guinea: forest, fallow edge, mangrove beach; found at 3-680 m.

SPECIMENS EXAMINED – **Guinea**: Guinée Forestière: Nzérékoré Region: Lola Préfecture, Mts. Nimba, between Mifergui Camp and barrier at the border, 16 Nov. 2007, *Jongkind 8023* (K001764230!, P04472980!, MO, WAG1541761!, SERG); Yomou Préfecture, Koniyakoele, Bamakama, forêt médicinale, 19 Jan. 2017, *Haba 690* (K00139794!); Nimba, Bakolé à 450m d’altitude, Zouepo, 20 May 1949, *Adam 5140* (P03524475!); Zouhouromuai/Macenta, 1 Dec. 1949, *Adam 7298* (P03524474!); Guinée Maritime: Boké Region: Boké Préfecture, Dougoula, W of Dougoula Village, 20 Nov. 2007, *van der Burgt 1024* (HNG, K001061444!, WAG); Coyah Préfecture, near to Manéah, Fourré, 26 Dec. 1978, *Lisowski 51020* (BR0000018868388!, POZG-V-0057053); Kindia Préfecture, Friguiagbé, 14 Dec. 1936, *Chillou 40* (BR0000018867459!, P03524482!); “Plantes de Konakry”, 1897, *Maclaud s.n.* (P03524483!, P03524485!); Kindia, environs de Kindia, Oct. 1905, *Pobéguin 1310* (P03524484!).

ADDITIONAL SPECIMENS – **Guinea**: Guinée Forestière: Nzérékoré Region: Yomou Préfecture, Forêt Diécké, 21 Nov. 2019, *Yarwoah Konaté & Koïvogui 70* [7°29’N 8°50’W] (BRLU, MO, SERG); Nzérékoré Region: Macenta Préfecture, Sérédou, 23 Jan. 1993, *Lisowski B-7388* (POZG-V-0057054); Yomou Préfecture, Diéké, 27 Jan. 1993, *Lisowski B-7501* (POZG-V-0057055); Nzérékoré Préfecture, au S de Nzérékoré, lisère d’une forêt primaire [7° 45’ 13’N 8° 49’ 12’W], 27 Jan. 1993, *Lisowski B-7410* (POZG-V-0057056); Macenta, 15 Nov. 1962, *Lisowski 27187* (POZG-V-0057057); Macenta, 11 Nov. 1962, *Lisowski 27188* (POZG-V-0057058); Macenta, 4 Dec. 1962, *Lisowski 27189* (POZG-V-0057059).

CONSERVATION STATUS (PRELIMINARY ASSESMENT) – LC (Least Concern), following the global IUCN (2012) guidelines, known from 127 occurrence points worldwide.

USES – Pounded and moistened leaves were reported to be inserted into the nostrils of hunting dogs to improve their ability to scent (Burkill *et al*. 1985).

## 6. Neuropeltis Wall

**1a.** Leaves glabrous beneath or nearly so; when hairy, the indumentum closely appressed to the lamina’s surface **1. N. acuminata**

**1b.** Leaves velutinous beneath **2. N. occidentalis**

### 6.1. *Neuropeltis acuminata* (P.Beauv.) Benth

*Porana acuminata* P.Beauv. in Fl. Oware 1: 66 (1806).

Type: Nigeria, 1813, *Palisot de Beauvois s.n.* [holotype: G (G00023036!); isotypes: G (G00146308!, G00146344!)].

Large *liana* or sometimes *shrub*; *stem* with many medullary rays, terete, 1-2 mm in diameter, white, turning rapidly brown, up to 40 m in length, bumpy, striate, glabrous or pubescent on the younger stems. *Leaf*: *petiole* 5-15(−25) mm long, narrowly canaliculate above, mostly glabrous or slightly puberulous; lamina entire, elliptic or sometimes ovate, (3-)5-10(−12) cm long, (2-)3-6 cm wide, cuneate to rounded at the base, acute to acuminate 5-15 mm and mucronate 1-3 mm at the apex, glabrous on both surfaces, upper surface shiny, lower surface dull; 7-10 pairs of secondary veins ascending and anastomosing towards the edge. *Inflorescence* axillary, racemose, densely-flowered, glabrous or pubescent at least when young, with adpressed, sparse hairs; *peduncle* 15-30 cm long, glabrous or sparsely pubescent; bracteoles elliptical or slightly obovate, 1.5-2 mm long, absent or very poorly developed; *bracteoles* suborbicular, base cordate, apex acute to emarginate, ± as long as the calyx, 4-6 mm in diameter, papery, covered in similar indumentum to the calyx, fused to the pedicels, enlarged in fruit, strongly accrescent and ± scarious after anthesis, flower buds globular, hairy. *Flower*: *pedicel* 1.5-2(−4) cm long, accrescent, hairy like the inflorescence; *sepals* subequal, semi-orbicular to broadly elliptical, 1.5-3 mm long, persistant, brown pubescent, imbricate. *Corolla* campanulate, 3-9 mm long, white or sometimes tinged with green, sparsely pubescent on the outside, 5 refracted lobes, 2-3 mm long, with hyaline margins; *stamens* included, glabrous, filaments as long as, or longer, than the corolla, 4-9 mm long, filiform, broadened at the base, inserted on the corolla tube, anthers oblong, 1-1.5(−2) mm long, medifixed; *disc* poorly developed, 0.2-0.5 mm high; *pistil* (3-)4-7.5 mm long; *ovary* conical, 0.8-2 mm long, 1-2 mm wide, pubescent at the top, 2-celled, 2-ovuled per cell; *styles* 2, free, divergent, glabrous or sparsely pubescent, equal or subequal; each stigma 0.8-1 mm long, peltate, greenish white, with 2 flattened, wavy, and irregularly lobed branches. *Fruit*: capsule, ovoid, 7-8mm long, mm wide, sparsely pubescent, glabrescent towards the top, rarely glabrous, surrounded at the base by the persistent calyx, 1-2-seeded, indehiscenct, inserted ± in the middle of the midrib of the very enlarged bract. *Seed* black, glabrous, finely pustular.

DISTRIBUTION – From West tropical Africa to Angola. In Guinea: Guinée Maritime and Guinée Foréstière.

HABITAT – In Guinea: secondary forest, found at 28-500 m (elsewhere up to 1700 m).

SPECIMENS EXAMINED – **Guinea**: Guinée Maritime: Kindia Region: Forécariah Préfecture, Souganyah, 3km from Senguelen, 0.25km from new village, 23 Sept. 2015, *Molmou 910* (HNG!, K000874476!, MO).

ADDITIONAL SPECIMENS – **Guinea**: Guinée Forestière: Nzérékoré Region: Nzérékoré Préfecture, Nzérékoré, 27 Jan. 1993, *Lisowkski 90655* (POZG-V-0059273); au S de Nzérékoré, lisière d’une forêt primaire [7° 45’ 13“N 8° 49’ 12”W], 27 Jan. 1993, *Lisowski B-7395* (POZG-V-0059278).

CONSERVATION STATUS (PRELIMINARY ASSESSMENT) – Least Concern (LC), following the global IUCN (2012) guidelines; known from 301 occurrence points worldwide.

USES - *N. acuminata* is eaten as a vegetable in Gabon. It also has additional uses as ropes in house construction in Ghana and Ivory Coast, and the plant fibres as dish sponges and toilet gloves in Cameroon (Kambiré *et al*. 2022).

### 6.2. Neuropeltis occidentalis Breteler

Type: Guinea, Guinée Forestière, Nimba Mts, Nzérékoré, Lola Préfecture, between Miferguire Camp and barrier at the border, 16 Nov. 2007, 16 Nov. 2007, *Jongkind et al. 8025* [holotype: WAG (WAG0296936!, WAG0296937!); isotypes: BR (BR0000009959446!), K (K001310590!), MO, P (P04528876!)].

*Liana*; *stem* 2-4mm in diameter, up to 25m high, trichomes along the stem erect to somewhat adpressed. *Leaf*: petiole (5-)7-15(−18) mm long, slender, ± terete, grooved above, brown-velutinous; lamina entire, obovate-elliptic, (3-)4-10 cm long, 2-5.5 cm wide, rounded to subcordate at the base, acuminate, 1-4(−10) mm long, mucronate at the apex, papery to coriaceous, upper surface pale brown to ± white, softly hairy, longer persistent hairs, or glabrescent, on the impressed midrib, lower surface sparsely to densely brown-velutinous, more densely so on the prominent midrib; (4-)6-8(−11) pairs of secondary veins. *Inflorescence* axillary, usually an unbranched raceme, up to 10cm long, appearing compound at the branched apex; peduncle 2-5cm long; bracteoles elliptic, (2-)3-5(−6) mm long, including the 1-2 mm long acumen, velutinous, fused to the pedicels, strongly accrescent and ± scarious after anthesis. *Flower*: *pedicel* 2-3mm long, subappressed brown-velutinous; *sepals* subequal, ± free, broadly ovate-elliptic, 1.5-3mm long, 1.5-2(−3) mm wide, persistant, brown pubescent outside, glabrous inside, imbricate.; corolla campanulate, 5-6 mm long, tube 2-4mm long, white, 5-lobed, lobes ovate-triangular, with hyaline margins, pubescent outside, glabrous inside; *stamens* included, glabrous, filaments 8-10mm long, filiform, broadened at the base, united with the corolla tube, anthers oblong, ± 1.5 mm long, medifixed; *disc* poorly developed, up to 1.5mm high, glabrous; pistil ± as long as corolla; *ovary* enlarged, 1mm long, adpressed-pubescent on apical part, rarely glabrous, 2-celled, 2-ovuled per cell; styles 2, free, villous to glabrous, equal or subequal, stigma subpeltate. *Fruit*: calyx lobes enlarged, circular to broadly elliptic, mucronate, 3-5cm long, 3-4cm wide, up to 5 mm long, sparsely golden-brown pubescent; *capsule* ovoid, up to 0.5cm long, glabrous, rarely with a few hairs apically, indehiscent, inserted ± in the middle of the midrib of the very enlarged bract. *Seed* glabrous.

DISTRIBUTION – West Africa (Guinea, Sierra Leone, Liberia and Ivory Coast). In Guinea: Guinée Forestière.

HABITAT – In Guinea: moist evergreen to semi-deciduous, lowland to submontane forest, forest edge; found at 680-740m.

SPECIMENS EXAMINED – **Guinea**: Guinée Forestière: Nzérékoré Region: Youmou Préfecture, 23 Feb. 1949, *Adam 3820* (P00635947!).

ADDITIONAL SPECIMENS – **Guinea**: Guinée Forestière: Nzérékoré Region: Lola Préfecture, Nimba Mountains, SMFG iron mining concession, Zougué Valley, 30 Sept. 2011, *McPherson 21479* (MO, P01029252!).

CONSERVATION STATUS (PRELIMINARY ASSESMENT) – Least Concern (LC), following the global IUCN (2012) guidelines, known from 41 occurrence points worldwide.

NOTE – Lisowski (2009) lists *N. velutina* as present in Guinea, but the only specimen cited (*Adam 3820*) has been integrated in this later described species *N. occidentalis* (Breteler 2010). As currently circumscribed, *N. velutina* does not occur in West Africa, and is restricted to Central Africa (Cameroon, Central African Republic, Congo, Gabon, Nigeria, Democratic Republic of Congo) (POWO 2024).

## 7. Calycobolus Willd. ex Schult

**1a.** Young stems and petiole hairy; anthers pendulous at anthesis **1. C. africanus**

**1b.** Young stems and petiole glabrous; anthers erect at anthesis **C. heudelotti**

### 7.1. Calycobolus africanus (G.Don) Heine

*Codonanthus africana* G.Don in Gen. Hist. 4: 166 (1837).

Type: Sierra Leone, *Don s.n.* [holotype K (K000405768!); isotypes: BM (BM000930522!), K (K000405769!), MPU (MPU010871!)].

*Liana* reaching the canopy of large trees; *stem* terete, up to 16 cm in diameter, dark brown, up to 10m high, pubescent with appressed hairs, to glabrescent. *Leaf*: petiole (5-)6-10(−16)mm long, subterete, grooved above, slightly pubescent with appr=essed hairs; lamina entire, obovate-elliptical to obovate, 2-2.5(−3.5) times as long as wide, (7-)10-18(−21)cm long, (3-)4-6(−8.5)cm wide, round to obtuse at the base, acuminate and usually mucronate at the apex, papery to subcoriaceous, appressed-pubescent on both surfaces when young, later upper surface glabrous, lower surface slightly pubescent; midrib depressed above, prominent beneath, 8-10 pairs of secondary veins, ± plane above, prominent beneath, arched and at the end going up on the preceding vein. *Inflorescence* axillary, in a fascicle in the leaf axils, up to 6-flowered, appressed-pubescent; peduncle 2-4 mm long, adpressed-pubescent; bracteoles 2, opposite, ± elliptical, inserted 1-2 mm above base of pedicel, not accrescent in fruit; bracteoles ± elliptical, 1-3mm long. *Flower*: pedicel (8-)10-16(−25) mm long, <0.5 mm in diameter, filiform, appressed-pubescent; sepals very unequal, 2 outer sepals strongly accrescent and membraneous in fruit, ± circular, base deeply cordate, apex obtuse, 7-10mm long and wide, with short appressed hairs, 3 inner sepals ovate-elliptic, apex acute, 3-4mm long, 4-7mm wide, margin ciliate, sparsely puberulous; corolla ± urceolate, 17-20mm long, usually with a narrow tubular part 3mm long at the base, white to pale pink, ± tomentose on apex, obtusely lobed ± 2 mm apically; stamens included, (5-)9-13mm long, filaments usually unequal, flattened at the base, ± glabrous to ± pustular-puberulous in a 2mm long zone above, 3 mm adnate to corolla tube, anthers pendulous, base sagittate, 2-4mm long, glabrous; disc 0.5 mm long, glabrous; pistil 10-17 mm long, glabrous; ovary 1-2mm long, 2-celled, 2-ovuled; styles 2, united in the lower half to slightly above, bifid, equal to slightly unequal, stigmas large, ellipsoidal, peltate. *Fruit*: calyx lobes, largest 5-6.5cm long, 5-5.5cm wide, smallest 2-2.5cm long, 2cm wide, glabrous; fruit indehiscent, ellipsoid, 10-13mm long, 6-7mm wide, glabrous, crowned by the basal part of the styles, 1-seeded, with a 1-2mm long style remnant at the apex, thin walled. *Seed* 1, glabrous, smooth.

DISTRIBUTION – West and West Central African. In Guinea: Guinée Forestière. HABITAT – In Guinea: forest, forest edge, roadside; found at 760-839 m.

SPECIMENS EXAMINED – **Guinea**: Guinée Forestière: Nzérékoré Region: Macenta Préfecture, Mts. Ziama, Papo sur la route de l’antenne à Seredou, 7 Feb. 2010, *Goman 227* (K000683678!, HNG, WAG1541912!); near to Macenta, 13 May 1963, *Lisowski 10632* (BR0000018868982!, BR0000018868999!, POZG-V-0057077); Lola Préfecture, Nimba

Mountains, between Mifergui Camp and Zougué River, 11 Dec. 2006, *Jongkind 7603* (WAG1541913!).

ADDITIONAL SPECIMENS - **Guinea**: Guinée Forestière: Nzérékoré Region: Macenta Préfecture, Macenta, 6 Jan. 1963, *Lisowski 91008* (POZG-V-0057078).

CONSERVATION STATUS (PRELIMINARY ASSESSMENT) – Least Concern (LC) following the global IUCN (2012) guidelines, known from 295 occurrence points worldwide.

### 7.2. *Calycobolus heudelotii* (Baker ex Oliv.) Heine

*Breweria heudelotii* Baker ex Oliv. in Hooker’s Icon. Pl. 23: t. 2276 (1894).

Type: Guinea, Fouta Djallon, 1847, *Heudelot 864* [lectotype: K (K000405774!); isolectotypes: A (A01154502!), BR (BR0000008885388!), G-DC (G00023114!), K (K000405772!, K000405773!, K000405774!), MPU (MPU010872!).

Small to stout *liana* reaching the crown of tall trees, or shrub; *stem* woody, terete, 3-6mm in diameter, striate, glabrous, sometimes sparsely puberulous when very young. *Leaf*: *petiole* (0.5-)1-3(−4)cm long, grooved to canaliculate above, glabrous; lamina entire, ovate-elliptic, 1.5-2 times as long as wide, (3.5–)8–15(17)cm long, (2.5–)5–7(–11)cm wide, obtuse or rounded to shortly cuneate at the base, acuminate,, rarely obtuse or rounded at the apex, papery to coriaceous, glabrous on both surfaces, except for some hairs at the entrance of the domatia, sometimes glandular-punctate on the lower surface; (6-)7-9 pairs of secondary nerves, slightly prominent above, distinctly so beneath, midrib impressed above, prominent beneath. *Inflorescence* axillary, 1-16-flowered panicle, rarely very shortly racemose; *peduncle* 2-4mm long, tomentose with whitish hairs; *bracteoles* obovate-elliptic to almost circular, 2-4mm long, glabrous above, opposite, attached to pedicel 1-5mm from the base, not accrescent in fruit; *bracteoles* elliptic, 1-1.5mm long. *Flower*: *pedicel* 6-13mm long, tomentose with whitish hairs; *sepals* unequal, broadly elliptic to circular, base deeply cordate, tomentose with whitish hairs outside, ± glabrous inside, 2 outer sepals strongly accrescent and membranous in fruit, 5-6mm long, 4-5mm wide, 3 inner sepals slightly narrower; corolla campanulate to funnel-shaped with 5 lobes, (16-)20-27 mm long, basally constricted for 1-2mm, white, puberulous with whitish hairs mainly on the upper half, outside, lobes oblong-triangular, 7-15mm long, ± spreading at anthesis; *stamens* included, (7-)11-13mm long, glabrous, *filaments* (2-)4-5 mm long, flattened at the base, united with the corolla tube, anthers longitudinally dehiscing, 2.5-4mm long; disc 1-2mm in diameter, 1mm thick, glabrous; pistil 14-20mm long, glabrous; ovary 1-2mm long, 2-celled, 2-ovuled; styles 2, bifid, 2/3 to almost completely united, very unequal in length, stigma ellipsoidal, peltate. *Fruit*: enlarged calyx lobes, ± hyaline, broadly elliptic to circular, deeply cordate at base, largest 3.5-6.5cm wide, smallest 1.5-3cm wide, inner ones 4-6mm long, ± glabrous to sparsely puberulous outside; fruit indehiscent, ellipsoid to oblongoid, 10-15mm long, 4-6 mm wide, glabrous, light green, glandular-punctate or not, crowned by the basal part of the styles, 1-seeded, thin walled. *Seed* 1, glabrous, smooth.

DISTRIBUTION – West Tropical Africa. In Guinea: mostly occurring in Guinée Forestière, with a few occurences in Guinée Maritime and Moyenne Guinée.

HABITAT – In Guinea: forest, forest edge, savannah shrub, roadside; found at 339-1260 m.

SPECIMENS EXAMINED – **Guinea**: Guinée Maritime: Kindia Region: Télimélé Préfecture, Pellel, ± 8k E of Sangaredi, 15 Feb. 2011, *Gandeka 158* (K001759465!, HNG); Kindia Préfecture, Kindia, *Morvan s.n.*, Jul. 1913 (L.2723783!); Moyenne Guinée: Labé Region: Veudou, 31 Mar. 1956, *Georges 11746* (BR0000018870053!, WAG0034430!,

WAG.1542044!); Guinée Forestière: Nzérékoré Region: Macenta Préfecture, Simandou Range, S. of Pic de Fon, 24 Mar. 2008, *van der Burgt & Couch 1152* (K, WAG0393379!); Lola Préfecture, Mounts Nimba, SMFG Mining Concession, 14 Feb. 2012, *Bidault, Diabaté & Mas 303* (BR0000018870060!, K, MO-2450682, P00853500, SERG, WAG); Lola Préfecture, Mounts Nimba, SMFG Mining Concession, Gouan Valley, 19 Feb. 2012, *Diabaté, Bidault & Mas 1259* (BR, P00853724!, MO, SERG, WAG); Lola Préfecture, Monts Nimba, April 1949, *Portères s.n.,* (P01183713!); Mount Nimba, between Mifergui Camp and Zougué River, 11 Dec. 2006, *Jongkind*, *Nema & Holië 7602* (WAG0237380!, WAG.1542042!, WAG0049652!); Mount Nimba, N. of Mifergui Camp, 12 Dec. 2006, *Jongkind*, *Nema & Holië 7606* (WAG0237189!, WAG.1542043!, WAG0049652); West slope of Mont Yonon, 6 Feb. 2012, *Yonon Botanic Team 120* (WAG0405619!, WAG1542041!).

ADDITIONAL SPECIMENS - **Guinea**: Guinée Forestière: Nzérékoré Region: Macenta Préfecture, Ziama Biosphere Reserve on the Sibata II side, 23 Feb. 2019, *Diabaté 3585* (SERG, MO); Macenta, 23 Jan. 1993, *Lisowski B-7320* (POZG-V-0057081); *loc. cit*., 23 Jan. 1993, *Lisowski B-7322* (POZG-V-0057082); Macenta Préfecture, Sérédou, *Lisowski B-7434*, 23 Jan. 1993 (POZG-V-0057080).

CONSERVATION (PRELIMINARY ASSESSMENT) –Least Concern (LC) following the global IUCN (2012) guidelines, known from 385 occurrence points worldwide.

## 8. Aniseia Choisy

### 8.1. Aniseia martinicensis (Jacq.) Choisy

C*onvolvulus martinicensis* Jacq. in Select. Stirp. Amer. Hist.: 26 (1763).

Type: Martinique, ‘circa vicum Roberti’, *Jacquin s.n.* [holotype: BM (BM000953205)].

Perennial *herb*; *stem* climbing or creeping, terete, 1-2mm in diameter, longitudinally striate, thinly pubescent or glabrescent. *Leaf*: *petiole* 0.5-2cm, thinly pubescent or glabrescent; lamina entire, narrowly elliptical, 1.5-9(−10.5)cm long, 0.5-3cm wide, cuneate or sometimes rounded at the base, obtuse but minutely mucronate at the apex, glabrescent or thinly pubescent on both surfaces, particularly at the margins and on the nerves below; midrib protruding below, depressed above, 5-7 pairs of secondary veins. *Inflorescence* axillary cyme, often solitary, sometimes in 3-flowered cyme; *peduncle* 1-4 cm long, glabrous to sparsely hairy; *bracteoles* narrowly triangular, 2mm long, 1mm wide, inserted 0.5-5mm below the calyx. *Flower*: *pedicel* 1-6mm long, pilose; *sepals* unequal, ovate-elliptical, apex acute, 8-16mm long, enlarging to 22mm in fruit, 6-9mm wide, decurrent on the pedicel, margin sparsely pubescent on the outside, becoming scarious, and slightly accrescent in fruit, 3 outer sepals larger, ovate, 8-15(−23)mm long, 6-9mm wide, margin reticulately veined, inner sepals narrowly elliptical-oval, apex acuminate, 8-15mm long, 3-4mm wide; corolla infundibuliform, 2cm long, 2.5cm wide, white, limb hairy, barely lobed at apex, midpetaline bands well-defined, puberulous; *stamens* 5, included, filaments equal, 4-6.5mm long, filiform, broadened and covered with glandular hairs at the base, inserted ± 2mm from the bottom of the tube, anthers erect, base sagittate, 2mm long; pollen 3-colpate; ovary disc annular, 0.3mm high, not very prominent; *ovary* ovoid, 2mm long, glabrous, 2-celled, 4-ovuled; style 1, 8.5mm long, simple, obscurely articulated in the lower quarter, glabrous, included; stigmas 2, globular or oblong. *Fruit*: capsule elongate-ovoid or globose, 1.5cm long, glabrous, septum incomplete, reduced to 2 membranous fimbriate wings, surrounded by accrescent sepals up to 22mm long, 2-celled, dehiscent by 4 valves. *Seeds* 4, trigonal or globose, 4.5mm in diameter, black, scattered hairs on the surface, tuft of brown hairs near the hilum.

SPECIMENS EXAMINED – **Guinea**: Guinée Maritime: Kindia Region: Kindia Préfecture, Friguiagbé, 29 Jan. 1943, *Jacques-Georges 26768* (WAG1541551!); Kafutchez: Nov. 1956, *Jacques-Felix 7294* (BR0000018864052!); Boké Region: Boké Préfecture, near to Kamsar, 20 Jan. 1979, *Lisowski 51188* (BR0000018864069!, POZG-V-0057030); Forécariah Préfecture, Mafarenya-Forecariah, river crossing on main road, 8 May 2012, *Sow 875* (HNG0000815!, K).

ADDITIONAL SPECIMENS – **Guinea:** Haute Guinée: Kankan Region: Kankan Préfecture, Kankan, *Lisowski 10639,* 20 Dec. 1966 (POZG-V-0057027); *loc. cit*., 2 Jan. 1967, *Lisowski 10638* (POZG-V-0057029); *s. loc*., 20 Jan. 1979, *Lisowski 51188* (POZG-V-0057030).

DISTRIBUTION – Native to tropical and subtropical America, introduced to tropical Africa, Asia and Australia. In Guinea: Guinée Maritime and Haute Guinée.

HABITAT – In Guinea: humid rice field, river edge, gallery forest, palm forest; found at altitudes of 10-400 m.

CONSERVATION STATUS (PRELIMINARY ASSESSMENT) –LC (Least Concern) on the IUCN Red List (Kumar 2011).

USES – The leaves of *A. martinicensis* are eaten in Malaysia and Indonesia (Kumar 2011).

## 9. Hewittia Wight & Arn

### 9.1. Hewittia malabarica (L.) Suresh

*Convolvulus malabaricus* L. in Sp. Pl.: 155 (1753).

Type: Rheede, Horti malabarici pars undecima, 1962, *11: 105, t. 51*.

Perennial *herb*; *stem* protruding or climbing, terete, 1-2mm in diameter, up to 2m long, sometimes rooting at the base, puberulous. *Leaf*: *petiole* 0.5-10cm long, puberulous; lamina entire or basally lobed, ovate very variable, 2-17cm long, 1.5-14cm wide, cordate or hastate at the base, obtuse to acute or acuminate and mucronate at the apex, ± pubescent to puberulous on both surfaces; 3 basal veins, 3-5 pairs of secondary veins. *Inflorescence* axillary, compressed cyme, 1-few-flowered, puberulous; *peduncle* 0.5-19cm long, angular, pubescent; numerous bracteoles, ovate to elliptic, apex 8-10(−13)mm long, 2-4(−5)mm wide, margin ciliate, situated at the top of the peduncle, persistent. *Flower*: *pedicel* 2-10mm long, pubescent; *sepals* unequal, apex acute, with clearly visible veins, margin hyaline, pubescent, 3 outer sepals accrescent in fruit, ovate to broadly elliptical, 10-18mm long, 6-9mm wide, 2 inner sepals narrowly elliptical to narrowly elliptical-ovate, 0.9-12mm long, 2-2.5mm wide. corolla infundibuliform, 2-2.5cm long, yellow or yellowish to whitish with a dark red funnel, pubescent on the outside, unlobed, mid-petaline bands hairy; *stamens* subequal, filament 5-9mm long, flattened and broadened at the base, with small capitate hairs at the insertion, fused approximately 4mm from the base of the corolla tube, free, anthers ellipsoid, 2mm long; pollen grain smooth 12-15 pantocolpate; *disc* annular, 0.4mm high, finely papillose; *ovary* globose to ovoid, 1mm long, with long erect hairs, 1-celled, 4-ovuled; *style* 1, 10mm long, filiform, pubescent at the base, *stigmas* 2, globular, very finely papillate. *Fruit*: capsule globose, 8-10mm long, hairy, crowned by accrescent sepals, 1-celled, 4-valved. *Seeds* 4, obovate-globose, 5-6mm long, black, glabrous, with a warty integument enveloping the seed.

SPECIMENS EXAMINDED – **Guinea**: Guinée Forestière: Nzérékoré Region: Lola Préfecture, W of Nimba Mountains, near Gbakoré, 6 Aug. 2008, *Jongkind 8302* (MO-3259968!, WAG127116!).

ADDITIONAL SPECIMENS – Guinea: Guinée Forestière: Nzérékoré Region: Guéckédo Préfecture, Guéckédo, 17 Dec. 1962, *Lisowski 90660* (POZG-V-0057294); Guinée Maritime: Kindia Region: Coyah Préfecture, Maneah Tanene, 31 Jul. 2011, *Koumassadouno 255* (HNG0000243!); Guinée Maritime: Kindia Region: Coyah Préfecture, Friguiady, 31 Jul. 2011, *Sow 187* (HNG0000179!).

DISTRIBUTION – Tropical Africa, Asia and Polynesia. In Guinea: across the southern border, in Guinée Maritime and Guinée Forestière.

HABITAT – In Guinea: disturbed open vegetation, roadside, river edge; found at altitudes of 139-505 m.

CONSERVATION STATUS (PRELIMINARY ASSESSMENT) – LC (Least Concern) following the global IUCN (2012) guidelines, known from 909 occurrence points worldwide.

USES - The leaves are eaten as a vegetable by being boiled and pounded with groundnuts or coconut milk, either alone or with other vegetables, and eaten with a staple food (Ruffo *et al*. 2002). The leaves are used for a traditional dish called ‘onyebe’ in Uganda by the Langi people (Grubben & Denton 2004). *H. malabarica* has medicinal uses, particularly in folk medicine. The leaves are rubbed into sores and a root decoction is used for treatment of Oxyuris threadworms (Burkill *et al*. 1985). It is used as an ornamental plant and is grown for ground cover, particularly in plantations of ylang-ylang *Cananga odorata* in Madagascar (Grubben & Denton 2004). In Benin, it is used for cattle fodder and the inner bark fibres are used for rope making (Ouachinou *et al*. 2018; Grubben & Denton 2004).

## 10. Camonea Raf

### 10.1. *Camonea umbellata* (L.) A.R.Simões & Staples

*Convolvulus umbellatus* L. in Sp. Pl.: 155 (1753).

Type: Icon in Pluknet, Phytographia plate 167, fig. 1. (1979).

Climbing or creeping, twining, slightly woody *vine*; *stem* slender, cylindrical, 1-2mm in diameter, green or copper, reaching 5m, wrinkled, striate, glabrous or sparsely pubescent with simple greyish hairs, with scarce milky latex and a pair of spiniform projections at the nodes. *Leaf*: *petiole* 1-5cm long, cylindrical, pubescent with simple whitish hairs, two paired auricles at the base; lamina entire, ovate or lanceolate, 3-17cm long, 2.5-12cm wide, cordate or sagittate at the base, obtuse, acute or acuminate and mucronate at the apex, margin undulate, chartaceous, upper surface yellowish green, puberulous with minute white hairs, lower surface glabrous but puberulous with white hairs on the veins; 5-7 pairs of basal veins, 5-7 pairs of secondary veins, sunken beneath. *Inflorescence* umbelliform axillary cymes; *peduncle* 0.5-2cm long, angular or cylindrical, pubescent with whitish hairs. *Flower*: *pedicel* 1.5-2.5cm long, same hairiness as peduncles; sepals unequal, ovate or rounded, 0.5-1 cm long, green, glabrous; corolla infundibuliform, 2.5-3cm long, yellow, limb ± 3 cm in diameter, glabrous, lobes obtuse; *stamens* included, white, filaments unequal, longest 9mm long, shortest 5mm long, anthers oblong, 3.5mm long, 1mm wide; pollen6-zonocolpate; style 1, filiform; stigmas 2, globose greenish, slightly exserted. *Fruit*: capsule globose, 1cm long, brown, covered by the persistent sepals, 4-valved, exocarp not delaminating during dehiscence. *Seed* obtusely trigonal, 5-8mm long, brown, velvety, with a line of longer hairs on two of the margins.

DISTRIBUTION – Widespread across tropical and subtropical regions. In Guinea: across the southern border, in Guinée Maritime and Haute Guinée.

SPECIMENS EXAMINED – **Guinea**: Guinée Maritime: Télimélé Préfecture: Kérékéré, next to River Kogon, E of gallery forest, 16 Feb. 2011, *Molmou 14* (HNG, K001754703!); Haute Guinée: Mamou Region: Dalaba Préfecture, Kaba, 16 Feb. 2011, *Chevalier 20391* (K001754704!, P00434245!, P00434246!, P00434247!).

ADDITIONAL SPECIMENS – **Guinea**: Guinée Maritime: Kindia Region, Forécariah Préfecture, Forécariah, 30 Dec. 1978, *Lisowski 51409* (POZG-V-0059274); Haute Guinée: Mamou Region: Dalaba Préfecture, Kaba, 22 Feb. 1979, *Lisowski 51852* (POZG-V-0059275).

CONSERVATION STATUS (PRELIMINARY ASSESSMENT) – LC (Least Concern) following the global IUCN (2012) guidelines, known from 3,586 occurrence points worldwide.

USES – The leaves of *C. umbellata* are eaten as a vegetable in Asia but it is also known for medicinal, environmental and material uses (POWO 2024; Ooststroom & Hoogland 1954). A poultice or paste of the leaves sooth the skin so is used to treat burns, abscesses, ulcers, sores and, combined with Curcuma powder, is used to treat cracks in the hands and feet (Ooststroom & Hoogland 1954).

## 11. Distimake Raf

**1a.** Plant hirsute with yellowish hairs, 3-3.5 mm long, with tuberculate base; leaves palmately compound, leaflets 5; corolla completely white **1. D. aegyptius**

**1b.** Plant glabrous or rarely slightly pubescent; leaves palmately lobed, lobes 5-7; corolla white to cream with purple middle, or yellow **2**

**2a.** Corolla white to dull pale yellow, with dark purple centre; stems distinctly muriculate with reddish papillae; external sepals shorter **2. D. kentrocaulos**

**2b.** Corolla golden yellow; stems smooth; external sepals longer **3. D. tuberosus**

### 11.1. *Distimake aegyptius* (L.) A.R.Simões & Staples

*Ipomoea aegyptia* L. in Sp. Pl.: 162 (1753).

Type: America calidiore, Herb. Linn. No. 218.35 [lectotype: LINN (LINN-HL218-35!), as *Convolvulus pentaphyllus* L., nom illeg., replaced by *Ipomoea aegyptia* L.].

Annual *herb*; *stem* twining or prostrate, branched, slender, cylindrical, 2-4mm in diameter in diameter, up to 2m, with black dots, hirsute, with yellowish hairs 3-3.5mm long, with tuberculate base. *Leaf*: *petiole* 1.4-15cm long, hirsute like the stem; lamina palmately 5-lobed, lobes elliptical to obovate, 2.5-13cm long, 1-4(−4.3)cm wide, acuminate and mucronate at the apex, margin ciliate, sparsely pubescent on both surfaces, especially on the veins; pinnate segmental venation, 9-11 pairs of secondary veins. *Inflorescence* axillary cyme, (1 or)2-10-flowered, hirsute; *peduncle* 1.5-6cm long, cylindrical, hirsute with the same yellowish hairs as stem; *bracteoles* narrowly oblong to narrowly elliptical-ovate, 2-3.5(−6)mm long, villous. *Flower*: *pedicel* 1-2(−2.5)cm long, ± inflected, hirsute; *sepals* unequal, free, apex obtuse, 10-15mm long, reaching 30mm in fruit, 3 outer sepals oval, covered with long golden hairs up to 5mm, coriaceous, 2 inner sepals largely elliptical, entirely glabrous, less coriaceous; corolla infundibuliform, with a very short and rapidly widening tubular base, 2-2.5(−3.5)cm long, white, glabrous; *stamens* 5, included, filaments unequal, flattened and broadened at the base, anthers sagittate, slightly rolled up; pollen 3-colpate; *disc* annular; *ovary* ± 1mm long, glabrous, 4-celled, 4-ovuled; *style* 1, filiform, included, *stigmas* 2, capitate. *Fruit*: capsule ovoid, acute at apex, 12-14mm long, 8-13mm wide, glabrous, rigid, surrounded by persistent, hirsute sepals, 4-valved. *Seeds* 4, ± 5 mm long, light brown, glabrous.

SPECIMENS EXAMINED – **Guinea**: Guinée Maritime: Conakry, 27 Apr. 1902, *s.c. 314,* (K001743067!); Boké Préfecture: Kabata, N to NW of village Kabata, 21 Nov. 2007, *Fofana 93* (HNG, K001061442!).

DISTRIBUTION – Widespread in tropical and subtropical regions. In Guinea: Guinée Maritime.

HABITAT – In Guinea: forest, roadside; found at altitudes of 10 m.

CONSERVATION STATUS (PRELIMINARY ASSESSMENT) – LC (Least Concern) following the global IUCN (2012) guidelines, known from 549 occurrence points worldwide.

USES – The plant is grown for ornamental use but is also known to be used to feed fodder, as medicine and in construction, as fibre; the dried leaves are applied in Nigeria as a dressing for burns, and the stems used as ties in Senegal (Burkill, 1985; POWO 2024; Mwanga Mwanga *et al*. 2022).

### 11.2. *Distimake kentrocaulos* (C.B.Clarke) A.R.Simões & Staples

*Ipomoea kentrocaulos* C.B.Clarke in J.D.Hooker, Fl. Brit. India 4: 213 (1883).

Type: Ethiopia, Takkaza River, *Schimper 800* [syntypes: BR (BR0000008250827!), S (S11-38674!), K (K000097236!), P (P00434239!), P (P00434240!)].

Perennial, twining *liana*, glabrous; *stem* becoming woody, herbaceous but firm, terete, younger stems slender, 1-2mm in diameter, up to 15m long, usually distinctly muriculate with reddish papillae. *Leaf*: *petiole* 2-6cm long; lamina 5-7-lobed, palmatisect, pentagonal in outline, 4-15cm long and wide, cordate with a narrow sinus at the base, lobes oblong to lanceolate, apex obtuse to subacute or attenuate, lobes margin entire or crenate; 5-7 basal veins, each reaching to a lobe, numerous pairs of secondary veins. *Inflorescence* axillary cymes, up to 20cm in length, 1-few-flowered; *peduncle* 1-9cm long, patent to suberect, usually distinctly muriculate with reddish papillae; *bracteoles* ovate, acute, concave, 3-5mm long, early deciduous, occasionally larger and dissected like leaves. *Flower*: *pedicel* 2-10mm long, at first deflexed, patent to suberect when flowers open and ultimately cernuous in fruit, usually distinctly muriculate with reddish papillae; *sepals* subequal, free, ovate-oblong or elliptic, concave, apex obtuse or rounded and minutely mucronate, up to 30mm long, ± 12mm wide, coriaceous with thinner submembraneous edges, glabrous, accrescent in fruit, outer sepals coriaceous, inner sepals longer, less coriaceous; corolla infundibuliform, 4-6 cm long, 6-8cm wide, white to dull pale yellow or buff with dark purple centre, limb faintly 5-angled, plicate, glabrous, mid-petaline bands not sharply defined; *stamens* 5, included, *filaments* flattened and broadened at the base; *pollen* 3-colpate; *disc* annular; *ovary* 4-ovuled; *style* 1, filiform, included, *stigmas* 2, globose. *Fruit*: capsule narrowly ellipsoid, 12-15 mm wide, pale brown at first, enclosed in accrescent brown calyx but ultimately exposed just before dehiscence when sepals spread out, 4-valved, circumscisile at the base. *Seeds* 4, 8-9 mm long, ± 6 mm broad, brown-black, minutely pubescent.

SPECIMENS EXAMINED – **Guinea**: Moyenne Guinée: Tougué Préfecture: Kansangui, Bani Misside, Waladji, ± 2k W of Misside, 20 Nov. 2018, *Molmou 1975* (HNG, K000875237!, MO).

DISTRIBUTION – Tropical Africa and India. In Guinea: Moyenne Guinée, in Kansanguin, Tougué Préfecture.

HABITAT – In Guinea: herbaceous savannah; found at the altitude of 783m.

CONSERVATION STATUS (PRELIMINARY ASSESSMENT) – LC (Least Concern) following the global IUCN (2012) guidelines, known from 217 occurrence points worldwide.

**1.3. *Distimake tuberosus*** (L.) A.R.Simões & Staples

*Ipomoea tuberosa* L. in Sp. Pl.: 160 (1753).

Type: Jamaïque, *Herb. Linn. 219.4* [lectotype: LINN (LINN-HL219-4!)].

Perennial *liana*, glabrous or rarely slightly pubescent; *stem* reaching treetops, branched, cylindrical, slightly striate, 4mm in diameter, with underground tubercule. *Leaf*: *petiole* 6-15(−18)cm long; lamina 5-7-lobed, palmatifid to palmatipartite, 6-16cm long and wide, acuminate at the apex, lobes narrowly elliptical-ovate to narrowly elliptical-obovate or oblong, lower lobes smaller than the terminal, venation pinnate, clearly visible on both sides, basal veins in each lobe, numerous secondary veins. *Inflorescence* axillary, a loose cyme, few-flowered or sometimes reduced to a single flower; *peduncle* 15(−20)cm long; *bracteoles* triangular, ± 2 cm long. *Flower*: *pedicel* 1.6-1.8cm long, elongating to 5cm and becoming clavate in fruit; *sepals* unequal, free, glabrous, accrescent in fruit, 2 outer sepals larger, ovate, apex obtuse and mucronate, 2-3cm long, coriaceous, 3 inner sepals narrower, oblong, 2-2.5cm long, less coriaceous; corolla infundibuliform, long tubular at the base, crenate at the top, up to 5.5 cm long, 6cm wide, golden yellow, glabrous; stamens 5, included, filaments unequal, flattened at the base, anthers spiralling twisting at dehiscence; pollen 12-zonocolpate; *disc* annular; *ovary* 4-ovuled; *style* 1, filiform, glabrous, included, *stigmas* 2, globose. *Fruit*: capsule subglobose to ellipsoidal, 3.5cm in diameter, glabrous, thin pericarp, surrounded by hard accrescent sepals, reaching 6cm in length, 2-celled, 2-4-seeded, irregularly dehiscing. *Seeds* 2-4, ± 17mm long, 14mm wide, blackish, ± pubescent with short black trichomes covering the entire surface, more densely so along the margins.

DISTRIBUTION – Native to tropical America, introduced to tropical Africa, America, Asia and Australia.

HABITAT – Found at altitudes of 200-1600 m, globally.

CONSERVATION STATUS (PRELIMINARY ASSESSMENT) – LC (Least Concern following the global IUCN (2012) guidelines, known from 549 occurrence points worldwide. VERNACULAR NAMES – *rose de yaoundé*.

USES – *D. tuberosus* has ornamental use for its flower, capsule and calyx; the tuber can be used medicinally as a purgative (Mwanga Mwanga *et al*. 2022).

SPECIMENS EXAMINED – **Guinea.** Guinée Maritime: Kindia Region: Coyah Préfécture, plantantion Jacquard, cultivée, 14 Feb. 1939, *Chillou 1165* (P03544932!, P03544935!).

## 12. Stictocardia Hallier f

### 12.1. Stictocardia beraviensis (Vatke) Hallier f

*Ipomoea beraviensis* Vatke in Linnaea 43: 514 (1882).

Type: Madagascar, Beravi interior Gebirge, juil. 1879, *Hildebrandt 3094* [holotype: K (K000097044!); isotypes: A (A00054662!), L (L0004255!), M (M0109983!), P (P00427176!)].

*Liana*; *stem* twining, older twigs reaching 1cm in diameter in diameter, wrinkled surface and densely covered with tuberous lenticels, densely pubescent young stems, later yellowish glabrescent. *Leaf*: *petiole* 3-15cm long, canaliculate, grey tomentose; lamina entire, ovate, up to 16(−23) cm long and 15(−20.5) cm wide, cordate or truncate at the base, acute or blunt and mucronate at the apex, upper surface glabrous or very sparingly pilose, brownish green, covered with small glands, lower surface very densely grey-velvety to glabrous or glabrescent; venation pinnate, depressed above, protruding below, 3-5 basal veins, 9-13 pairs of closely parallel secondary veins. *Inflorescence* axillary cymes, many-flowered; *peduncle* subsessile, 0.8-1(−2.5)cm long; bracteoles small, caducous, flower buds long and conical, finely pubescent at the tips. *Flower*: *pedicel* 0.6-1cm long, tomentose; *sepals* subequal, free, elliptic or sub-orbicular, apex blunt or emarginate, mucronulate, persistent and accrescent in fruit, glabrescent above, pubescent towards the base, 2 outer sepals orbicular, 7-13mm long, 11-12mm wide, pubescent, more densely so at base and along the midrib, 3 inner sepals elliptical, apex rounded and slightly notched, 10-11mm long, 5-8mm wide, glabrous; corolla infundibuliform, gradually tapering towards the base, the basal part narrower, 4.5-5.5 cm long, very bright crimson with base of tube orange-yellow, with 5 barely marked lobes ending in a tuft of hairs like those of the flower buds, mid-petaline bands often somewhat pubescent and with minute glands as the leaves; *stamens* 5, included, *filaments* subequal, 2.1-3.5cm long, free, inserted near the base of the corolla tube, *anthers* narrowly ovoid, base sagittate, 4-5mm long; pollen spinulose pantoporate; *ovary* glabrous, 4-celled, 1-ovule per cell; *style* 1, 3.5-3.8cm long, simple, glabrous, stigmas 2, globose. *Fruit*: capsule globose, 1.4cm long, 1.3cm wide, straw-coloured, woody, pericarp thin, irregularly dehiscing, completely enclosed by the accrescent calyx.. *Seeds* 4, 6mm long, 4mm wide, black, pubescent, ± granular.

DISTRIBUTION – Widespread in Tropical Africa, Madagascar and India. In Guinea: Guinée Forestière and Guinée Maritime.

HABITAT – In Guinea: palm forest, edge of disturbed forest, river edge; found at 126-650m (elsewhere up to 1800m).

CONSERVATION STATUS (PRELIMINARY ASSESSMENTS) – LC (Least Concern) following the global IUCN (2012) guidelines, known from 222 occurrence points worldwide. USES – Ornamental. (Mwanga Mwanga *et al*. 2022).

SPECIMENS EXAMINED – **Guinea**: Guinée Maritime: Télimélé Préfecture: Lamba, Bank of river Tinguilinta, 20 Feb. 2011, *Molmou 74* (HNG, K001587409!); Guinée Forestière: Nzérékoré: Mts. Nimba, between Mifergui Camp and barrier at the border, 16 Nov. 2011, *Jongkind 8042* (BR0000009959224, K001587410!, MO-2894509!, WAG1748231); Macenta Préfecture, Sérédou, 2 Dec. 1964, *Fora 21* (BR0000018302929!).

## 13. Lepistemon Blume

### 13.1. *Lepistemon owariense* (P.Beauv.) Hallier f

*Ipomoea owariensis* P.Beauv. in Fl. Oware 2: 40 (1816).

Type: Nigeria, près d’Oware, *Palisot de Beauvois s.n.* (P).

Perennial *herb*; *stem* branched, twining, tuberculate base, robust, up to 3m, covered with long appressed or spreading yellow-brown bristly hairs with a tuberculate base. *Leaf*: *petiole* 5-15cm long, hairy like the stem; lamina entire, sinuate or shallowly lobed, broadly ovate, 5-16cm long, 4-18cm wide, deeply cordate at the base, acute to emarginate at the apex, lobes deltoid, about 1cm long and 2cm wide, pilose hairs on both surfaces, the same as the stem; midrib depressed above, projecting below, 5-6 palmate basal veins, 4-5 pairs of secondary veins. *Inflorescence* compact cymes, forming pauciflorous; sessile or shortly peduncled; *bracteoles* minute. *Flower*: *pedicel* 3-20(−25)mm long, angular, glabrous to pilose; *sepals* subequal, free, ovate to elliptic, apex obtuse, ± cuspidate, 4.8-6mm long, 3.5mm wide, herbaceous or subcoriaceous, pilose with long yellow hairs to glabrous, inner sepals slighter longer; corolla urceolate, tube at first subcylindric, subcampanulate, becoming ovoid as the ovary expands, up to 1.8cm long, 16-25mm in diameter, white, limb up to 3cm in diameter, glabrous; *stamens* included, filaments 4-5mm long, inserted low down in the corolla tube, with triangular basal scale, 3-4mm long, glabrous, anthers obloid, base deeply sagittate, 2-2.5mm long, surrounded by staminal scales; *pollen* spinulose, pantoporate; *disc* annular; *ovary* ovoid to globose, 2-celled, 2-ovuled per locule; style 1, 2mm long, stigmas 2, globular. *Fruit*: capsule ovoid-globose, ± 15mm long, 13mm wide, hirsute with very long and rigid yellow hairs, setose except in the upper part, coriaceous, 4-valved or almost indehiscent, bursting irregularly 3-4-seeded. *Seeds* 3-4, globose or nearly so, 3.5-4(−5)mm in diameter, grey-black, glabrous or sparsely hairy, minute perforations in the pericarp.

SPECIMENS EXAMINED – **Guinea**: Guinée Forestière: Nzérékoré Region: Macenta (‘N’zoiaro’), *Adam 6852*, 6 Nov. 1949 (P03867179!); Haute Guinée: Kouroussa, *Pobéguin 1068,* Oct. 1902 (P03867177!, P03867178!).

ADDITIONAL SPECIMENS – **Guinea**: Guinée Forestière: Nzérékoré Region: Guéckédou Préfecture, Guéckédou, 27 Dec. 1962, *Lisowski 27175* (POZG-V-0059112); Haute Guinée: Faranah Préfecture, between Tiro and Sandankoro, 28 May 1967, *Lisowski 27174* (POZG-V-0059110, POZG-V-0059111).

DISTRIBUTION – Tropical Africa, Asia, Australia and Pacific Islands. In Guinea: Guinée Forestière and Haute Guinée.

HABITAT – In Guinea: savannah, forest edge, river edge; found at altitudes of 440-450 m in Guinea, elsewhere at 0-1350 m.

CONSERVATION STATUS (PRELIMINARY ASSESSMENT)– LC (Least Concern), following the global IUCN (2012) guidelines, known from 152 occurrence points worldwide. USES – The leaves are eaten as a vegetable (Burkill *et al*. 1985).

## 14. Argyreia

### 14.1. *Argyreia nervosa* (Burm.f.) Bojer

*Convolvulus nervosus* Burm.f. in Fl. Indica: 48 (1768).

Type: Inde [Coromandel], *Outgaerden s.n* [lectotype: G-PREL (G00816025!)].

Voluble *liana*; stem reaching 10 m, dense silvery white or tawny tomentose. *Leaf*: petiole 6.5-15(−20) cm long; lamina entire, ovate to orbicular, 10-30 cm long, 8-25 cm wide, cordate at the base, obtuse or acute to shortly acuminate and mucronate at the apex, upper surface glabrous to glabrescent, lower surface densely silky with silvery white to tawny grey and shiny tomentose; venation pinnate, prominent below, 4-5 pairs of subpalmate basal veins, 11-15 pairs of secondary veins. *Inflorescence* axillary, in a dense cyme, subcapitate, 6-12-flowered; peduncle 10-20 cm long, tomentose like the stem; bracteoles oval to elliptical or oblong, apex broadly acuminate, 3-5 cm long, glabrous above, tomentose below, deciduous, foliaceous. *Flower*: pedicel 1-2 cm long, angular; sepals unequal, white tomentose on the outside, 2 outer sepals broadly elliptical, apex slightly acute, ± 15 mm long, inner sepals broadly elliptical to suborbicular, apex obtuse, 10-12 mm long; corolla infundibuliform, the tube widened in the middle, with shallow lobes, (4-)6-7 cm long, mauve or purple, darker on the throat, tube and midpetaline bands, silky pubescent on the outside; 5 stamens included, filaments unequal, 2 longer and 3 shorter, broadened at the base, glabrous, inserted 8 mm from the bottom of the tube, anthers sagittate at the base, 5 mm long; pollen grain echinulate; disc annular, strongly undulated, 1.5 mm high, glabrous; ovary obpyriform, glabrous, 4-celled, 4-ovuled; 1 style 18 mm long, filiform, glabrous, stigmas 2, globose. *Fruit*: calyx lobes silvery pubescent on the outside outside; capsule ellipsoidal to subglobose, depressed, with short apex, ± 18 mm long, glabrous, incomplete septa, transformed into ossified tissue, grainy in appearance, surrounded by accrescent sepals, ≤ 4-seeded, indehiscent. *Seed* ellipsoidal, 10-12 mm long, 7-9 mm wide, beige, velvety with white hairs, longer around the hilum.

DISTRIBUTION – Native from India to Myanmar, cultivated in the tropics worldwide.

CONSERVATION STATUS (PRELIMINARY ASSESSMENT) – LC (Least Concern) following the global IUCN (2012) guidelines, known from 147 occurrence points worldwide.

USES – *A. nervosa* is cultivated for ornamental use due to its large purple flowers; the seeds contain LSA (D-lyseric acid amino) causing a hallucinogenic affect for which the plant is also cultivated (Mwanga Mwanga *et al*. 2022).

OBSERVATIONS – Between Sérédou and Nzérekoré, village on the riverbank of the Diani; Nzérekoré (Lisowski 2009).

NOTES – No specimens were found to document the presence of this species in Guinea; this record is based only on the observations by S. Lisowski (2009).

## 15. Ipomoea L

Key to the species of *Ipomoea* in Guinea-Konakry

**1a.** Leaves deeply 5-7 palmately lobed **2.**

**1b.** Leaves entire, variable in shape, ovate, triangular to narrowly lanceolate or linear in outline, simple or shallowly 3-lobed **3.**

**2a.** Pair of bracts inserted at the base of the petiole, deeply palmately lobed like the leaves; leaves palmately 5-lobed nearly to base, the outer lobes often lobulate, acutely mucronate; corolla 4.5 cm long, not contracted at the base **1.I. cairica**

**2b.** No bracts at the insertion of the petiole; leaves usually 5-7-lobed to about the middle of the lamina, lobes acuminate; corolla c. 7 cm long, contracted at base **2**. **I. mauritiana**

**3a.** Stoloniferous plants, with sub-succulent leaves, often in aquatic environments, swamps or river banks, with a floating thick, fistulose or spongy stem **4.**

**3b.** Plant without stolon, occurring in terrestrial environments **7**

**4a.** Leaves narrowly to broadly triangular, linear-oblong, lanceolate or linear; corolla infundibuliform with a narrow tube, widening at the top, up to 8.5 cm long, purple, pink or white with a dark centre 3. **I. aquatica**

**4b.** Leaves broadly orbicular or reniform; corolla broadly infundibuliform, up to 6.5 cm long, white, yellow, purple or red-purple, with or without dark centre **5.**

**5a.** Leave broadly orbicular to reniform; apex obtuse to shortly acuminate 4. **I. asarifolia**

**5b.** Leaves reniform; apex emarginate to deeply bilobed **6.**

**6a.** Corolla white or yellow, without dark centre, 3.5-5cm long **I. imperati**

**6b.** Corolla pink or red-purple with darker centre, 3-6.5 cm long 6. **I. pes-caprae**

**7a.** Inflorescences in densely clustered, pedunculate, heads; entire plant covered in yellowish and whitish indumentum **8.**

**7b.** Inflorescences not in densely clustered, pedunculate, heads **11**

**8a.** Bracteoles subtending the inflorescence united into an oval involucre, 3-6 cm long, up to 5 cm in diameter 7. **I. involucrata**

**8b.** Bracteoles subtending the inflorescence not united, 1.4-3 cm long **9**

**9a.** Plant erect; inflorescences and leaves sessile to subsessile; leaves oblong-lanceolate, mucronate, truncate or subcordate at the base 8. **I. argentaurata**

**9b.** Plant twining; inflorescences long pedunculate, peduncle and petioles conspicuous; leaves ovate-subcordate **10.**

**10a.** Outer bracteoles foliaceous, partly covering the inflorescence, up to 3 cm long and 1 cm wide, ovate-elliptical to narrowly oblong; corolla 1.3-2.4 cm long, purple or white………... 9. I. heterotricha

**10b.** Outer bracteoles not foliaceous, not covering the inflorescence, up to 2.5 cm long, ovate-elliptical to narrowly elliptical, apex acute to acuminate; corolla 3-5.5 cm long, mauve to red-violet 10. **I. chrysochaetia var. velutipes**

**11a.** Inflorescences subsessile, many-flowered or sub-solitary **12**

**11b.** Inflorescences supported by a conspicuous peduncle, not subsessile **15**

**12a.** Leaves oblong ovate, narrowly cordate, lanceolate or linear **13**

**12b.** Leaves very variable, from ovate-triangular to narrowly lanceolate, widely cordate, mucronate **14**

**13a.** Leaves oblong-lanceolate; sepals 0.6 cm long; corolla the same length or slightly longer than the calyx, 0.5-0.8cm long, red or white **11. I. coscinosperma**

**13b.** Leaves linear or oblong-linear; calyx 1.5-1.8cm long; corolla thrice as long as the calyx, up to 5.5 cm long, mauve with darker centre **12**. **I. blepharophylla**

**14a.** Leaves ovate to linear-oblong, sub-hastate with rounded lobes at the base, long-attenuate to acuminate, 2.5-8cm long, 0.6-4cm wide, flowers generally clustered, rarely solitary or paired; petiole up to 4.5cm; sepals lanceolate, 0.7cm long, pubescent; corolla1 cm long; fruit pilose **13**. **I. eriocarpa**

**14b.** Leaves ovate, widely cordate at the base, rounded and mucronate at the apex, 3-5 cm long, 1.5-3.5 cm wide, long-petiolate, petiole up to 9cm long; flowers solitary or paired; sepals oblong, 0.3-0.4 mm long, thinly pilose; corolla 0.6-0.8cm long; fruit glabrous……………….. **14**. **I. biflora**

**15a.** Bracteoles conspicuous, 1.2-2 cm, ovate, long mucronate, outer sepals finely aristate, abaxially 5-winged; fruit enclosed by the enlarged sepals **15**. **I. setifera**

**15b.** Bracteoles inconspicuous, not as long; sepals subequal, or outers sepals not aristate nor winged; fruit not enclosed by the sepals **16.**

**16a.** Sepals distinctly awned at or below the apex; awn straight or curved; corolla salver-shaped with a long and narrow tube; stamens and style mostly exserted **17**

**16b.** Sepals obtuse, acute or acuminate, whether mucronulate or not, but not distinctly awned at or below the apex; corolla mostly funnel-shaped, or campanulate, sometimes salver-shaped; stamens and style mostly included, sometimes exserted **19**

**17a.** Corolla scarlet, rarely pure white, rather small, 3-4·5 cm. long; outer sepals 2-4·5 mm. long (without awn); inner ones 3-6 mm. (without awn); leaves pinnately parted into numerous linear or filiform segments, rarely less deeply pinnately cut 16. **I. quamoclit**

**17b.** Corolla white with greenish bands or pale bluish-purple, 5-15 cm. long; outer sepals 5-12

mm. long (without awn), inner ones 7-15 mm (without awn) **18**

**18a.** Corolla white; tube not or slightly widened above, 7-12 cm. long; limb rotate; stamens and style exserted **17**. **I. alba**

**18b.** Corolla purplish; the tube distinctly widened above, 3-6 cm. long, the limb funnel-shaped to rotate; stamens and style not or scarcely exserted **18**. **I. muricata**

**19a.** Petiole carrying a pair of glands (extra-floral nectaries) at the insertion of the leaf (observed on the abaxial face); sub-shrub or shrub **20**.

**19b.** Petiole without glands at the insertion of the leaf; climbing or prostrate herb **21**.

**20a.** Corolle hypocrateriform **19**. **I. verbascoidea**

**20b.** Corolle infundibuliform **20**. **I. carnea**

**21a**.Leaves oblong ovate, narrowly cordate, lanceolate or linear **22**

**21b.** Leaves ovate, triangular or cordate, apex acuminate, not linear or narrowly cordate or lanceolate **24**

**22a.** Calyx verrucose outside **I. barteri**

**22b.** Calyx not verrucose outside **23**

**23a.** Leaves oblong-ovate to triangular-lanceolate, base retuse to shallowly cordate, apex obtuse, mucronate; bracteoles conspicuous, linear, up to 0.7 cm long; sepals up to 12 mm. long. **22**. **I. pyrophila**

**23b.** Leaves cordate-ovate, acute, mucronate; bracteoles inconspicuous, minute, triangular 0.2-0.3cm long **23**. **I. tenuirostris**

**24a.** Stems and leaves mostly glabrous **25**

**24b.** Stem or leaves softly tomentose or pubescent, sometimes becoming glabrescent **27**

**25a.** Corolla salver-shaped, with a long narrow tube, 7-8 cm long, white and/or pale greenish-yellow **24**. **I. violacea**

**25b.**Corolla infundibuliform, 1.5-5 cm long, mauve, violet to white with purple centre **26**.

**26a.** Corolla 1.5-2.5 cm long **25**. **I.triloba**

**26b.** Corolla 3-5cm long **26**. **I.batatas**

**27a.** Corolla completely glabrous **28**

**27b.** Corolla pubescent outside **30**

**28a.** Ovary 2-4-locular, fruit 4-valved. Leaves entire, variable, ovate to triangular or narrowly ovate-elliptical, 4-16.5 cm long, 1-15 cm wide, cordate, sagittate, hastate or truncate at the base, acuminate to acute and apiculate at the apex; corolla white to mauve, sometimes purple or violet inside **27**. **I. sagittifolia**

**28b.** Ovary 3-locular, fruit 3-valved; corolla blue, purple, mauve or magenta **29**

**29a.** Leaves entire or 3-lobed, palmatifid, ovate to circular, 4-14 cm long, 3-13.5 cm wide, cordate at the base, acuminate at the apex, lobes ± acuminate at the apex, middle lobe ovate to oblong, acuminate, lateral ones obliquely ovate to broadly falcate; corolla magenta to mauve with paler tube, often white inside, glabrous **28**. **I. nil**

**29b.** Leaves entire or 3-lobed, palmatifid, ovate in outline, 5-12cm long, 3-15cm wide, cordate at the base, acuminate at the apex, lobes acuminate; corolla blue or mauve-purple, often red-tinged, tube whitish at the bas **29**. **I. indica**

**30a.** Corolla yellow, 1.5-2.5 cm long **30**. **I. obscura**

**30b.** Corolla purplish-red, pink, lilac or white, 4-5 cm long **31**. **I. rubens**

### 15.1. *Ipomoea cairica* (L.) Sweet

*Convolvulus cairicus* L. in Syst. Nat., ed. 10. 2: 922 (1759).

Type: Icon, t.70, Vesling in De Plantis Aegypti (Alpino 1640) (lectotype, designated by Bosser and Heine 2000: 32).

Perennial *herb*, glabrous or nearly so; *stems* twining or prostrate, terete, 1-2mm in diameter, up to 2m long, from a storage rootstock, smooth or muriculate, glabrous or villous at the nodes. *Leaf*: *petiole* 2-6cm long, glabrous, usually pseudostipulate by small leaves of developing or supressed axillary shoots; lamina 5-7-lobed, palmatisect, ovate to circular in outline, (2-)3-10cm long, 3-10cm wide, lobes lanceolate to ovate, elliptic or somewhat oblanceolate, acute or obtuse and mucronulate at the apex, up to 5cm long and 1.6cm wide, outer lobes often bifid, glabrous on both surfaces; 10 pairs of secondary veins per segment. *Inflorescence* lax axillary cymes, dichotomous, 1-many-flowered; *peduncle* 0.5-8cm long, glabrous, branched; *bracteoles*1.5 mm long, caducous. *Flower*: *pedicel* 1.2-3 cm long, thickened towards apex, glabrous; *sepals* subequal, ovate, apex obtuse to acute and mucronulate, 4-6.5mm long, 2.5-3(−3.2)mm wide, margin membranous, glabrous, sometimes verruculose, persistent in fruit, outer sepals slightly shorter, oblong-ovate, apex acute, often abasically rugose, 5-7mm long, 4mm wide, margin membranous and pale, inner sepals broadly ovate-elliptic, apex obtuse, 6-8mm long; corolla broadly infundibuliform, 4.5-6 cm long, 4.5-5 cm wide, purple, red or white with purple centre and purple tinge on outside of the limb or rarely entirely white, limb 4cm in diamater, glabrous, unlobed; *stamens* 5, included, *filaments* unequal, 10-20mm long, broadened and pubescent at the base; pollen spinulose, pantoporate; *disc* annular, ± 1 mmlong; *ovary* 2-celled, 4-ovuled; *style* 1, ± 18 mm long, filiform, glabrous, included, *stigmas* 2, globose. *Fruit*: capsule subglobose, 0.9-1.3cm long, 1-1.2cm wide, glabrous, 4-valved. *Seeds* 4, sub-globose or ovoid, 3.5-6 mm long, ± 4 mm wide, blackish-brown, densely short-tomentose with long silky hairs along the edges.

DISTRIBUTION – Widely spread throughout tropical and subtropical regions. In Guinea: Guinée Maritime, near the coast.

HABITAT – In Guinea: near settlements; found at 10-120 m (elsewhere up to 2,600 m). CONSERVATION – LC (Least Concern) on the IUCN Red List (Allen 2017).

USES – This plant is cultivated for ornamental use and has medicinal uses to treat rheumatism and inflammation, and in Brazilian folk medicine (Wood et al. 2020; Meira et al. 2012; Allen, 2017).

SPECIMENS EXAMINED – **Guinea**: Guinée Maritime: Conakry, 23 Mar. 1949, *Portères s.n.* (P01183706!).

ADDITIONAL SPECIMENS – **Guinea**: Guinée Maritime: Conakry, 23 Feb. 1967, *Lisowski 90302* (POZG-V-0057441); *loc. cit*., 24 Feb. 1967, *Lisowski 27181* (POZG-V-0057442).

### 15.2. Ipomoea mauritiana Jacq

Type: Plant, reputedly from Maurice (Mauritius), cultivated in Vienna, probably not preserved, possible type tab. 200 in Hort. Schoenb. (Jacquin, 1797).

Perennial large *liana*, glabrous; stem twining, occasionally prostrate, becoming woody, branched, terete, sometimes winged when old, 2-6 mm in diameter, up to 8 m long, fistulous glabrous, with large, inedible storage roots. *Leaf*: petiole 3-11 cm long, smooth or muriculate, puberulous; lamina entire or 3-9-lobed, palmatipartite, orbicular in outline, 6-24 cm long, 6-18 cm wide, cordate or truncate at the base, acuminate at the apex, lobes lanceolate to ovate, acuminate at the apex, upper surface glabrous to puberulous, lower surface glabrous, sometimes sparsely hairy on the veins; venation webbed, 5-7(−9) basal veins corresponding to the lobes, 2-8 pairs of secondary veins. *Inflorescence* pedunculate axillary, occasionally compound cymes, few-many-flowered, puberulous; peduncle 2.5-20 cm long, terete but often angular and cymosely branched near the apex, glabrous; bracteoles oblong, 2-4 mm long, 1-2 mm wide, margin caducous, flower buds globular. *Flower*: pedicel 0.9-2.5 cm long, terete, cylindrical, puberulous with very short hairs; sepals subequal, ovate, convex, clasping the corolla-tube, orbicular or elliptic, 6-12 mm long, coriaceous, margin hyaline, glabrous, persistent in fruit; corolla infundibuliform, with spreading limb and the tube narrow below, 5-6 cm long, 3.2-7 cm wide, reddish-purple or mauve with darker centre, glabrous, lobes deep and spreading; stamens included, filaments unequal, 14-18(−20) mm long, broadened and hairy at the base, anthers obloid, base sagittate, 3-4 mm long, white; pollen grain echinate; disc annular, cupuliform, slightly lobed, glabrous; ovary conical, glabrous, 2-celled; 1 style filiform, puberulous, 2 stigma globular. *Fruit*: capsule ovoid or globose, obtuse at apex, 1.2-1.4 cm long, 0.8-1.5 cm wide, glabrous, 2-celled, with cotton around seeds. *Seed* 6-7 mm long, black, covered with ± 7 mm long silky hairs.

DISTRIBUTION – Native to southern Africa and tropical America, introduced in Asia and Australia. In Guinea: Guinée Maritime, Hautê Guinée and Guinée Forestière.

HABITAT – Bushland at edge of bowal, savannah with shrubs, trees, near agricultural fields, savannah-grassland, inselberg belt forest, roadside, river bank; found at altitudes of 2-562 m in Guinea, elsewhere at 3-2450 m.

CONSERVATION STATUS (PRELIMINARY ASSESSMENT) – LC (Least Concern) following the global IUCN (2012) guidelines, known from 1,871 occurrence points worldwide.

USES - Once pounded and stewed, the grated tuber is used medicinally for abscesses and as an enema against constipation or venereal diseases (Mwanga Mwanga *et al*. 2022). It is known for treating an enlarged liver or spleen, heavy menstruation and gastrointestinal disorders. It also a lacto-stimulant and libido enhancer (Srivastava & Rauniyar 2020).

NOTE - One specimen [Guinée Maritime: Conakry, *Morvan s.n.*, Jun. 1913 (L2740748)] is of mixed content: *I. alba* (inflorescence and fruit) and *I. mauritiana* (leaves). There is no other record of *I. alba* in Guinée Maritime, suggesting that the occurrence may refer to *I. mauritiana*, to which elements of *I. alba* from a different collection were mistakenly added.

SPECIMENS EXAMINED – **Guinea**: Guinée Forestière: Nzérékoré Region: Lola Préfecture, Mts. Nimba, right bank of River Cavaly, 10 Jun. 2012, *Bilivogui 188* (K001740119!, MO-2451393!, BR, SERG, P, WAG); Lola Préfecture, Mts. Nimba, savanne entre les rivières Gba et le Yé et la piste menant à Serengbara, 6 Jul. 2012, *Mas 1368* (MO-2452107, P, SERG, WAG); Guéckédou Préfecture, Guéckédou, bord d’un chemin, *Lisowski 90305,* 28 Dec. 1962 (BR0000016092600!, POZG-V-0057763); Haute Guinée: Faranah Préfecture, Kouria, 4 Jul. 1905, *Chevalier 14944* (P00434182!, P00434183!); *s. loc*., 15 Jul. 1905 *Chevalier 14945* (P00434184!); Guinée Maritime: Boké Préfecture: Sangaredi, Para Gogo, Dalagala bowal, 19 Nov. 2013, *Fofana 205* (HNG, K000749898!); Tarensa village, 1k SE of village, nearest town Kamsar, 16 Nov. 2007, *van der Burgt 962* (HNG, K001061439!); Tarensa village, outskirts of village, 18 Nov. 2007, *Couch 462* (HNG, K001061440!); Bindêlya, 4 Dec. 1901, *Paroisse 31* (K001740120!); Kindia Region: Forécariah Préfecture, Senguelen Moofanyi inselberg, 28 Sep. 2012, *Cheek 1597* (HNG0001557, K).

ADDITIONAL SPECIMENS – **Guinea:** Guinée Maritime: Conakry: Cite de Kassa, 1 Dec. 1967, *Glaess s.n.* (POZG-V-0057760); Haute Guinée: Kankan Region: Kankan, 8 Jul. 1967, *Lisowski 90308* (POZG-V-0057761); Faranah Préfecture, Bordou, *Lisowski 10624,* 9 Jun. 1967 (POZG-V-0057765); Guinée Forestière: Nzérékoré Region: Macenta Préfecture, Seredou, 10 Dec. 1962, *Lisowski 90304* (POZG-V-0057762); Macenta, 28 Nov. 1962, *Lisowski 90307* (POZG-V-0057764).

### 15.3. Ipomoea aquatica Forssk

Type: Yémen, ad Zebid, 5 April 1763, *Forsskål 447* (holotype C [C10002419!]).

Perennial aquatic *herb*; *stems* several from a stout woody base, prostrate or floating, semi-succulent, terete, hollow, 4-6mm in diameter, 2-3m long, rooting at the nodes, striate, glabrous or slightly puberulous, with simple hairs. *Leaf*: *petiole* 3-25cm long, glabrous; lamina entire or coarsely dentate, very variable, narrowly to broadly triangular, linear-oblong, lanceolate or linear, 3-15cm long, 0.5-9cm wide, truncate, cordate or rarely rounded or sagittate to hastate at the base, acute, acuminate or rarely obtuse and mucronulate at the apex, glabrous on both surfaces with black-brown dots; 5-9 indistinct basal veins, 6-8 pairs of secondary veins. *Inflorescence* lax, pedunculate axillary cymes, (1-)2-5-flowered; peduncle 1-14cm long, thinner than petiole, glabrous expect for the pilose base; bracteoles narrowly ovate, apex acute, 1.5-2mm long. *Flower*: *pedicel* 2-6.5cm long, variable in length in the same plant, glabrous; *sepals* subequal, lanceolate-ovate, apex blunt or ± acute, 6-12mm long, 3-6mm wide, ± tuberculate, margin thin and pale, persistent in fruit, outer sepals slightly shorter, ovate-oblong, apex obtuse, often mucronulate, 7-8mm long, margin pale, glabrous, inner sepals ovate-elliptical, apex mucronulate, ± 8 mm long; corolla infundibuliform with a narrow tube (2.5-)4.5-8.5 cm long, purple or pink, or white with a deeper centre, limb 2.5cm in diameter, glabrous; *stamens* 5, unequal, *filaments* longest up to 10mm long, shortest 4-5mm long, broadened and hairy at the base, *anthers* obloid, base sagittate, ± 2 mm long; pollen spinulose, pantoporate; *disc* annular; *ovary* obpyriform, 3mm long, glabrous, 2-celled, 4-ovuled; *style* 1, up to 13mm long, articulated, filiform, *stigmas* 2, globular. *Fruit*: capsule globose to ovoid, 8-10mm in diameter, glabrous, tardily dehiscent, 4-valved. *Seeds* 4, ovoid, 6-6.5mm long, 5-5.5mm wide, densely pubescent.

DISTRIBUTION – Widespread in Tropical Africa and Asia, introduced in tropical America and Pacific. In Guinea: across all four provinces.

HABITAT – In Guinea: swamp, roadside, savannah; found at 65-538 m (elsewhere up to 1900 m).

CONSERVATION STATUS – LC (Least Concern) (Gupta & Sayer 2018).

USES – The young shoots, leaves and sometimes roots are eaten as a vegetable (Mwanga Mwanga *et al*. 2022). The leaves contain adequate amounts of most of the essential amino acids so could have potential use a food supplement and also provide a good source of calcium, magnesium, iron, zinc and copper (Meira *et al*. 2012). It is also known to provide fodder for livestock (Mwanga Mwanga *et al*. 2022) and has uses for medicine, fuel, gene sources, materials and social uses (POWO 2024). The leaves and stems are used for a laxative in India (Gupta & Sayer 2018). Other medicinal uses may be as a scorpion venom antidote, emetic, diuretic, purgative, and to treat liver problems, ringworm, leucoderma, leprosy and fever (Meira *et al*. 2012).

SPECIMENS EXAMINED – **Guinea**: Guinée Forestière: Macenta Region: near to Koenkan, 8 Dec. 1962, *Lisowski 95433* (BR0000016073449!, POZG-V-0057332); *Oudney 5* (BM000930424); Guinée Maritime: Kindia Region: Télimélé Préfecture, Koba, Nov. 1956, *Jacques-Felix 7236* (BR0000016073418!, K001594022!, K001594023!); Haute Guinée: Kankan Region: Kankan Préfecture, 1km NE. of Kankan, 20 Nov. 1966, *Lisowski 95432* (BR0000016073432!, POZG-V-0057329); Moyenne Guinée: Koundara Region: 23k S of Koundara, 12 Jan. 1979, *Lisowski 51931* (BR0000016073425!, K001594024!, POZG-V-0057326).

ADDITIONAL SPECIMENS – **Guinea**: Haute Guinée: Farannah Region, *Kawalec s.n.,* (POZG-V-0057311, POZG-V-0057312, POZG-V-0057313, POZG-V-0057314).

### 15.4. *Ipomoea asarifolia* (Desr.) Roem. & Schult

*Convolvulus asarifolius* Desr. in J.B.A.M.de Lamarck, Encycl. 3: 562 (1792).

Type: Senegal, *Roussillon s.n.* (holotype: P-LA[P-LAM00357544]; isotype: P-JUSS[P-JUSS6798]).

Perennial *herb*; *stem* prostrate or sometimes twining, much branched, thick, terete or angular, 1-5mm in diameter, rooting at the nodes, caniculate, glabrous. *Leaf*: *petiole* 3-8.5cm long, thick, deep longitudinal groove above, smooth or minutely muricated, glabrous; lamina entire, circular to reniform, 3-7cm long, 3.5-8.5cm wide, truncate to shallowly cordate with rounded auricles at the base, rounded to obtuse at the apex, subcoriaceous, glabrous on both surfaces; 5-7 basal veins, 5-7 pairs of secondary veins. *Inflorescence* axillary cymes, sometimes umbellate from apex of peduncle, 1-few-flowered, often together with an axillary leaf shoot; *peduncle* 1-2.5cm long, angular, glabrous; *bracteoles* ovate, 1-2mm long. *Flower*: *pedicel* 1.5-3cm long, glabrous; *sepals* unequal, elliptic-oblong, apex obtuse, mucronulate, glabrous, persistent in fruit, outer sepals shorter, 5-8mm long, 4mm wide, ± muricate, inner sepals 8-11mm long, 6-7mm wide; corolla infundibuliform, up to 6.5 cm long, red-purple, limb 3.5-4cm in diameter, glabrous, unlobed; pollen spinulose, pantoporate; *disc* annular; *style* 1, filiform, stigmas 2globose. *Fruit*: capsule globose, 10-12mm long, 8-10mm wide, glabrous, 4-valved. *Seeds* 4, 5-7mm long, 4 mm wide, minutely tomentellous.

DISTRIBUTION – Tropical West Africa, Asia and America. In Guinea: Guinée Maritime, Moyenne Guinée and Guinée Forestière.

HABITAT – In Guinea: forest edge, near the coast, roadside; found at 500-886 m (elsewhere up to 1200 m).

CONSERVATION STATUS (PRELIMINARY ASSESSMENT) – LC (Least Concern) following the global IUCN (2012) guidelines, known from 1737 occurrence points worldwide.

USES – This plant has medical uses to treat itch, urinary problems in pregnancy, haemorrhages, neuralgia, headaches, arthritis, and stomach ache (Meira *et al*. 2012). It can be used externally for dressing wounds, treating ophthalmia and guinea-worm sores, and combined with bulrush millet to create a steam bath to treat fever chills and rheumatic pain. The leaf and stems are combined with citron and water to be used as an oxytocic to stimulate contractions during labour. The leaves can be consumed to regulate blood pressure and the flowers are consumed to treat syphilis (Srivastava & Rauniyar 2020).

SPECIMENS EXAMINED – **Guinea**: Guinée Maritime: Ile du Kabak, Dec. 1956, *Jacques-Felix 7390* (K001591352!); Dec. 1956, *Pobéguin 1846* (K001591353!); Guinée Forestière: Nzérékoré Region: Lola Préfecture, Nimba Mountains, between Mifergui Camp and barrier at the border, 16 Nov. 2007, *Jongkind 8021,* (P, SERG, WAG1209032!).

ADDITIONAL SPECIMENS – **Guinea**: Moyenne Guinée: Labé Region: Mali Préfecture, Bandéa [11° 43’ 39“N 12° 13’ 27”W], 18 Feb. 1979, *Lisowski 51765* (POZG-V-0057353).

### 15.5. *Ipomoea imperati* (Vahl) Griseb

*Convolvulus imperati* Vahl in Symb. Bot. 1: 17 (1790).

Type: Imperato, unnumbered illustration by Imperati cited as “*Convolvulus marino*”, Hist. Nat. ed. 2: 671 (1672).

Perennial *herb*, glabrous; *stem* subsucculent,stoloniferous, creeping, sometimes twining, cylindrical, 1-3mm in diameter, reaching 5 m long, rooting at the nodes, striate, glabrous. *Leaf*: *petiole* 1-18cm long, canaliculate above, glabrous or sometimes with very few tip hairs; lamina entire or 3-5-lobed, linear to oblong or narrowly ovate-elliptical to ovate, 1.5-15cm long, 1-3.5(−7)cm wide, obtuse, truncate or cordate at the base, obtuse or emarginate to 2-lobed at the apex at the apex, glabrous on both surfaces, subsucculent, thick and fleshy, very variable in form and dimension, even on the same plant; venation pinnate, 5-10 pairs of secondary veins. *Inflorescence* axillary 1(to 3)-flowered, glabrous; *peduncle* 1-10cm long, glabrous; *bracteoles* very narrowly ovate-elliptical to linear or subulate, 2-4mm long, inserted at the base of the pedicel. *Flower*: *pedicel* 2-4.5cm long, slightly clavate, glabrous; *sepals* subequal, elliptical, apex obtuse and shortly cuspidate or mucronulate, 8-10mm long, 3mm wide, subcorieacous, glabrous, persistent in fruit, outer sepals 8-9mm long, 3-4mm wide, inner sepals 10-11mm long, 3-3.2 mm wide; *corolla* infundibuliform, 3.5-5cm long, white to yellow, with purple centre, glabrous externally, hairy internally at the throat, lobes broadly ovate, acute at apex, margin finely denticulate; *stamens* 5, included, *filaments* unequal, longest 8-10mm long, shortest 5-6(−7)mm long, pubescent at the base, *anthers* obloid, base sagittate, apex obtuse, 3-4mm long, longitudinally grooved, basifixed; pollen spinulose, pantoporate; *disc* annular; *pistil* included, 12-15mm long, glabrous; *ovary* ovoid, 2.2-2.5mm long, glabrous; *style* 1, 12-13mm long, filiform, *stigmas* 2, globose. *Fruit*: capsule subglobose, 10-12 mm long, glabrous, 2-celled, 4-valved.*Seeds* 4, ovoid-trigonal, ± 8 mm long, tomentose with wavy brown hairs.

DISTRIBUTION – Tropical and subtropical coasts worldwide. In Guinea: Guinée Maritime, near the coast.

HABITAT – In Guinea: white sand beach; found at 2-1,380 m globally.

CONSERVATION STATUS (PRELIMINARY ASSESSMENT) – LC (Least Concern) following the global IUCN (2012) guidelines, known from 4,233 occurrence points worldwide.

USES - *I. imperati* has medicinal use as a diuretic and to treat stomach issues, inflammation, swelling, wounds and pain after childbirth (Meira et al. 2012).

SPECIMENS EXAMINED – **Guinea**: Guinée Maritime: Boké Region: Boké Préfecture, Kamsar, Beach SW. of Taide Island, 26 Nov. 2013, *Guilavogui 675* (HNG, K000749899!, WAG); Boffa Préfecture, Bel-Air beach, 5 Feb. 1979, *Lisowski 51361* (K001594359!); near to Dupuru, Bongolondi, 5 Feb. 1979, *Lisowski 51430* (BR0000017392839!); Kindia Region: Télimélé Préfecture, Koba, 1 Nov. 1956, *Jacques-Félix 7265* (BR0000017392822!, K001594361!, L2729659!, WAG1744493!); Moyenne Guinée: Labé Region, Fouta Djallon, Mali, 23 Feb. 1949, *Portères s.n.* (P01183717!).

### 15.6. Ipomoea pes-caprae (L.) R.Br

*Convolvulus pes-caprae* L. in Sp. Pl.: 159 (1753).

Type: Herb. Linn. No. 218.59 (lectotype: LINN [LINN-HL218-59]).

Perennial herb, woody at base, glabrous; stem prostrate, often forming tangled mats, containing abundant white sap, fistulous, striated, cylindrical to angular or flattened, 3-7 mm in diameter, often purplish, 5-30 m long, rooting at the nodes. *Leaf*: petiole 3-6 cm long, purplish, with two glands at apex; lamina vary rarely rounded and entire, usually deeply 2-lobed, orbicular, obreniform, quadrangular or elliptic-3-10 cm long, 3-10.5 cm wide, rounded, cuneate or cordate at the base, conspicuously emarginate at the apex, coriaceous and somewhat succulent, held erect, often secund, glabrous on both surfaces, lower surface paler; prominently veined with glands near base of midrib, venation pinnate, 7-10 pairs of secondary veins. *Inflorescence* shortly pedunculate axillary cymes, 1-5-flowered; peduncle 3-16 cm long, erect, angular or flattened, secondary peduncle ending in 2 caducous bracteoles; bracteoles ovate-lanceolate, apex acuminate, 2-3.5 mm long, caducous. *Flower*: pedicel 1.2-4.5 cm long, thickened upwards; sepals subequal, ovate to elliptic-ovate, very concave, apex obtuse and mucronulate, 5-12 mm long, 5-7 mm wide, coriaceous, pale green, glabrous, persistent in fruit, outer sepals ovate to slightly elliptical, 6-12 mm long, 6 mm wide, margin distinctly 3-5-ribbed, inner sepals slightly larger, 8-15 mm long, 7-9 mm wide, margin scarious; corolla infundibuliform, 3-6.5 cm long, pink or red-purple with a darker centre, limb 4-5 cm in diameter, glabrous; stamens unequal, filaments 5-9 mm long, broadened and hairy at the base, anthers obloid, base sagittate, apex obtuse, 4-4.5 mm long; pollen grain echinate; disc annular, 0.6 mm high; ovary globose to ovoid, 2-2.5 mm long, glabrous, 2-celled; 1 style 12-20 mm long, filiform, somewhat persistent, 2 stigma globular. *Fruit*: capsule globular, 1.2-1.8 cm in diameter, glabrous, 4-valved, 4-seeded, pedicel often persistent on fallen capsule. *Seed* trigonal, 6-10 mm long, blackish-brown tomentose-villous.

DISTRIBUTION – Tropical and subtropical coasts worldwide. In Guinea: Guinée Maritime, near the coast.

HABITAT – In Guinea: sandy beach, coastal sites; found at 2-120 m (elsewhere up to-750 m).

SPECIMENS EXAMINED – **Guinea**: Guinée Maritime: Boké Préfecture: Kamsar, Beach SW. of Taide Island, 26 Nov. 2013, *Guilavogui 671* (HNG, K000749900!); Koba, 1 Nov. 1956, *Jacques-Felix 7264* (BR0000016100459!, BR0000016100442!, K001591310!).

ADDITIONAL SPECIMENS – **Guinea**: Guinée Maritime: Boké Region: Boffa Préfecture, Bel-Air [10° 14’ 16“N 14° 27’ 13”W], 5 Feb. 1979, *Lisowski 51402* (POZG-V-0057832); Conakry [9° 37’ 57“N 13° 35’ 15”W], 1 Oct. 1967, *Glaes s.n.* (POZG-V-0057831); Conakry, *Kawalec s.n.* (POZG-V-0057833).

CONSERVATION STATUS – LC (Least Concern) (Bárrios & Copeland 2021).

USES – This species is used in traditional medicine, particularly in the treatment of skin ailments such as wounds, itching, piles, sores, jellyfish stings, fish stings, snake bites and allergic reactions (Xavier-Santos *et al*. 2022; Chan *et al*. 2016; Burkhill *et al*. 1985). For example. In Thailand the leaves are mashed to a paste with vinegar and in China the leaves are used topically for pain, boils and bed sores (Chan *et al*. 2016). The diuretic and laxative properties of this plant mean it can also treat rheumatism, dropsy, urethral discharge, and, in Papua New Guinea, stomachache. It has additional spiritual uses in India as the leaves are used in ritual baths to eliminate evil spirits (Chan *et al*. 2016). Other uses include using the stoloniferous stems in rope making, and ecologically, the species is a pioneering coloniser of sand dunes and acts as a sand binder (Burkhill *et al*. 1985).

### 15.7. Ipomoea involucrata P.Beauv

Type: Nigeria/Benin, *Palisot de Beauvois s.n.* (holotype: G-DC [G00023040!]).

Exceedingly variable annual or perennial *liana*; *stem* slender, twining, prostrate or climbing, terete, ± 1 mm in diameter, up to 8 m long, with few black dots on surface, villose. *Leaf*: petiole 1-8 cm long, slender, retrorsely villose; lamina entire, ovate, 3-7(−11) cm long, 2.5-6(−7) cm wide, cordate at the base, acute or acuminate at the apex, membranous, hairy to villous on both surfaces, more densely beneath; 5 basal veins, 5 pairs of secondary veins. *Inflorescence* axillary capituliform cymes, few-many-flowered, a dense head enclosed in a large foliaceous boat-shape involucre; peduncle 1-16 cm long, pubescent with whitish simple hairs; outer bracteoles connate in a hairy boat-shaped structure, 3-6 cm long, 1-1.5 cm wide, with 2 cusps, ± pubescent, inner bracteoles 1.5-2 cm long, 0.2-0.4 cm wide, obovate to linear-oblong, acute to aristate. *Flower*: pedicel 1.5-3 mm long, pubescent with whitish simple hairs; sepals subequal, narrowly elliptical, 6-15 mm long, 1.5-4 mm wide, margin setose, villose, persistent in fruit, outer sepals lanceolate, inner sepals shorter, more ovate; corolla infundibuliform, 2-5.5 cm long, 2-5 cm wide at mouth, ± 4-10 mm wide at the base, purple, rose, white or white with pink throat, limb 3-4 cm diameter, glabrous, midpetaline bands sparsely pilose; stamens included, filaments unequal, longest 10-15 mm long, shortest 5-7 mm long, broadened and hairy at the base, anthers obloid, base sagittate, 2-2.5 mm long, basifixed; pollen grain echinate; disc annular; ovary ovoid, 1 mm long, glabrous; 1 style 8-13 mm long, filiform, 2 stigma globular. *Fruit*: capsule globose, 6 mm in diameter, glabrous, 4-valved. *Seed* trigonal, 3.5-4 mm long, blackish, glabrous or shortly pubescent.

DISTRIBUTION – Tropical and southern Africa. In Guinea: across all four provinces.

HABITAT – In Guinea: forest edge, river edge, near settlement, submontane forest, tree savannah with clearing, found at 10-1470 m (elsewhere up to 2700 m).

CONSERVATION STATUS (PRELIMINARY ASSESSMENT) – LC (Least Concern) following the global IUCN (2012) guidelines, known from 1,415 occurrence points worldwide.

USES – The leaves are consumed as a vegetable (Meira et al. 2012). It has traditional uses as a talisman for fertility in Gabon (Mwanga Mwanga *et al*. 2022). The leaf sap has medicinal use for oedema, swelling, gout, pulmonary issues and menstrual problems (Srivastava & Rauniyar 2020).

SPECIMENS EXAMINED – **Guinea**: Guinée Forestière: Nzérékoré Region, Beyla Préfecutre, Simandou, 11 Nov. 2005, *Harvey 264* (HNG, K000460107!); 7 Oct. 1947, *Baldwin Jr. 9681* (K001591241!); Lola Préfecture, Nimba Mountains, SMFG iron ore mine concession, East side of Pierre Richaud, 13 Oct. 2011, *Phillipson, Bidault & Bilivogui 6310* (MO-2721496, P, SERG, WAG); Mts. Nimba, plot JRSL05, 9 Dec. 2007, *Nimba Botanical Team 1777* (WAG1209858!); Mts. Nimba, Camp Gouan, 8 Dec. 1990, *Bah & Bah 345* (HNG0000330!); Guinée Maritime: Boké Region: Boké Préfecture, Tarensa village, 16 Nov. 2007, *Camara 26* (HNG, K001061441!); Boké Region: Boké Préfecture, Near to Kolaboui, Wourikaria, 15 Dec. 2011, *Keita, Doré & Sow 456* (HNG0000440!); Near to Conakry, Jun. 1913, *Morvan s.n.* (L2731218!); Kindia Region: Télimélé Préfecture, Kerekere, next to River Kogon, 16 Feb. 2011, *Molmou 05* (HNG, K001591240!); Kindia Préfecture, Friguiagbé, 12 Nov. 1938, *Chillou 879* (BR0000016085336!); Haute Guinée: Kouankan, 8 Dec. 1962, *Lisowski 90679* (POZG-V-0057689, POZG-V-0057688, WAG1209857!); 6 km NE. of Kankan, 10 Jan. 1967, *Lisowski 90680* (BR0000016085329!, POZG-V-0057674); Moyenne Guinée: Labé Region: Mali Préfecture, Téliré, 30 km NE of Labé, 31 Jan. 1990, *Malaisse 2529* (BR0000016085350!).

ADDITIONAL SPECIMENS – **Guinea**: Macenta Préfecture, Seredou, 20 Jan. 1993, *Lisowski s.n.* (POZG-V-0057662); Macenta, 10 Nov. 1962, *Lisowski s.n.* (POZG-V-0057693, POZG-V-0057700); Haute Guinée: Kankan Region: Kankan Préfecture, Kankan, 15 Dec. 1966, *Lisowski s.n* (POZG-V-0057694); Faranah Region: Kissidougou Préfecture, Dabadou, 12 Mar. 1967, *Lisowski s.n.* (POZG-V-0057701).

### 15.8. Ipomoea argentaurata Hallier f

Type: Togo, between Misahohoe and Bismarck Burg, Dec. 1891, *Buttner 746* (B, *n.v.*).

Annual *climber*; main *stem* erect, branched from the base, terete, 2-4mm in diameter, with dense adpressed short white hairs and longer spreading yellow golden hairs. *Leaf*: *petiole* 2-10mm long, hairy like the stem; lamina entire, oblong, 3.5-6cm long, 2-3cm wide, subcordate at the base, rounded and mucronate at the apex, upper surface densely strigose and deep green, lower surface silvery-silky; 3-5 basal veins, 7-9 pairs of secondary veins. *Inflorescence* axillary large cymes; *peduncle* 1-5cm long, densely strigose with long, golden-yellow hairs; bracteoles ±2.5cm long, hairy like the peduncles. *Flower*: *pedicel* 1-5 mm long, hairy like the peduncles; *sepals* equal, linear-lanceolate or almost linear, apex acuminate, 1.5-3cm long, silky white hairs on back with yellow strigose margin, persistent in fruit; corolla infundibuliform, 5cm long, nearly 4.5cm wide, whitish, hirsute, lobes entire, mid-petaline bands strigose on the outside; pollen spinulose pantoporate; *disc* annular; style 1, filiform, stigmas 2, globose. *Fruit*: capsule glabrous. *Seed* dense dark brown, pubescent.

DISTRIBUTION – West Tropical Africa to Chad. In Guinea: Moyenne Guinée and Haute Guinée.

HABITAT – In Guinea: roadside, shrub; found at 375-620m.

CONSERVATION STATUS (PRELIMINARY ASSESSMENT) – LC (Least Concern), following the global IUCN (2012) guidelines, known from 526 occurrence points worldwide.

USES – *I. argentaurata* has medicinal uses to treat depression and act as a genital stimulant. It also has social and religious uses, in superstitions, sayings and aphorisms (Burkill *et al*. 1985).

SPECIMENS EXAMINED – **Guinea**: Guinée Maritime: Near Conakry, Jun. 1913, *Morvan 302490* (L2741494!); Haute Guinée: Kankan Region: Kankan Préfecture, Route Kerouané-Kankan, au sud de Bissandougou, 16 Nov. 2022, *Bidault 6008* (BRLU, HNG, K, MO, P, SERG).

ADDITIONAL SPECIMENS – **Guinea**: Haute Guinée: Kankan, 2 Jan. 1967, *Lisowski s.n.* (POZG-V-0057338).

### 15.9. Ipomoea heterotricha *Didr*

Type: Guinea, *Mortensen s.n.* [lectotype: C (C10004084); isolectotype: C (C10004083 p.p)].

Annual *herb*, climber; *stem* prostrate or twining stems, cylindrical, 2-4mm in diameter, up to 1.5 mlong, pilose with long, spreading, yellowish simple hairs. *Leaf*: *petiole* 1.2-7.5cm long, densely pilose like the young shoots and peduncles; lamina entire, subtriangular to oval, 5.5-11.5cm long, 3.3-7.5cm wide, ± cordate at the base, acute at the apex, margin pilose, upper surface green with long adpressed hairs, lower surface silvery-velvety with dense adpressed hairs; venation pinnate, 10-13 pairs of secondary veins. *Inflorescence* axillary dense cymose,heads about 3cm wide, surrounded by persistent leaf-like bracteoles; *peduncle* 1-13cm long, densely pilose like the stem; *bracteoles* ovate-elliptical to narrowly oblong, outer bracteoles foliaceous, up to 3cm long, 1cm wide, inner bracteoles narrow, 1.4-1.7 cm long, 0.2-0.4 cm wide, margin similar hairs to the leaves. *Flower*: *pedicel* 1-10mm long, densely pilose with long yellowish hairs; *sepals* unequal, narrowly elliptical, apex acute, ± 9mm long, margins pubescent on both sides, hirsute externally, with long stiffish yellowish hairs towards the basal part, accrescent in fruit, shorter than the bracteoles, outer sepals linear-lanceolate, 6mm long; *corolla* infundibuliform, 1.3-2.4cm long, purple or white, with dark purple center, covered with long stiff, yellowish hairs on the upper part of the limb, slightly lobed and notched at the apex; *stamens* 5, included, *filaments* unequal, 5-6mm long, linear, broadened and hairy at the base, *anthers* ovoid, base sagittate, 1.8mm long, basifixed; *pollen* spinulose, pantoporate; *disc* annular; *ovary* slightly conical to ovoid, 5-6mm long, glabrous; *style* 1, 9-13 mm long, filiform, *stigmas* 2, globose. *Fruit*: capsule ovoid, 5-6mm long, glabrous, yellow green, often enclosed by calyx, 2-celled, 4-valved. *Seeds* 4, ovoid, 3-3.4 mm long, black, puberulous with yellowish-white hairs.

DISTRIBUTION – Tropical Africa. In Guinea: Guinée Maritime and Guinée Forestière.

HABITAT – In Guinea: fallow land, brushland at edge of bowal, grassland, roadside, forest edge; found at 244-992 m (elsewhere up to 3,000 m).

CONSERVATION STATUS (PRELIMINARY ASSESSMENT) – LC (Least Concern) following the global IUCN (2012) guidelines, known from 785 occurrence points worldwide.

SPECIMENS EXAMINED – **Guinea**: Guinée Forestière: Boké Préfecture, Mignan mountain 17 Nov. 2007, *Tchiengue 3028* (HNG, K000615476!, WAG); Nzérékoré, N. of Oueleba Camp, 16 Nov. 2007, *Cheek 13691* (K000436478!, WAG1209681); Nzérékoré Region: Lola Préfecture, Monts Nimba, en remontant vers l’amont de la Rivière Mion, 12 Nov. 2012, *Bilivogui, Diabaté & Mas 241* (MO-2451199, SERG, WAG); Macenta Préfecture, Kouankan, 8 Dec. 1962, *Lisowski 27192* (BR0000016083363!, WAG1209709!, WAG1209710!, POZG-V-0057618); Near to Zoubouroumai, brousse secondaire [8°32’19’’N 9°28’25’’W], 27 Nov. 1962, *Lisowski 27193* (BR0000016083370!, POZG-V-0057613); Guinée Maritime: Beyla Préfecture, Sangaredi, Para Gogo, Dalagala bowal, 19 Nov. 2013, *Fofana 207* (HNG, K000749897!).

ADDITIONAL SPECIMENS – **Guinea**: Guinée Maritime: Kindia Region: Dubréka, lisière d’une fôret dégradée [9°47’23“N 13°30’54”W], 27 Dec. 1978, *Lisowski 51025* (POZG-V-0057615).

### 15.10. *Ipomoea chrysochaetia* Hallier f. var. velutipes

*Ipomoea velutipes* Welw. ex Rendle

Type: Gabon, Luangr, Chinchene, Quelle von Maknaga, 1 Jun. 1874, *Soyaux 83* [holotype: M (M0109946!), isotype: K (K000097027!)].

Perennial *herb*; *stem* climbing, woody in the lower parts, terete, 2-3mm in diameter, 15m long, striated, hirsute to tomentose or pubescent to glabrous with short yellow or greyish simple hairs. *Leaf*: *petiole* 2-8(−13)cm long, hairy like stems; lamina entire, ovate, 4-13(−18)cm long, 2-12cm wide, cordate at the base, acute to acuminate at the apex, membranous to coriaceous, upper surface glabrous to tomentose or pubescent, lower surface pubescent to ± tomentose with long greyish simple hairs, especially on the veins; 5-7 basal veins, 3-5 pairs of secondary veins. *Inflorescence* axillary, subcapitulated or rarely loose, many-flowered; peduncle 2-32 cm long, robust, hirsute to pubescent with greyish simple hairs; *bracteoles* oval to narrowly oval-elliptical or sublinear, apex acute to acuminate, 1.7-2.5cm long, outer bracteoles larger, glandular and shortly hairy on the dorsal side. *Flower*: pedicel 2-7mm long, densely pubescent with simple greyish hairs; *sepals* unequal, narrowly oval-elliptical, apex 10-13mm long, margin sparsely pubescent, more densely so towards the base, becoming broadly ovate and acuminate in fruit, outer sepals 10-14mm long, 2-4mm wide, inner sepals 7-9mm long, 2-3mm wide; corolla infundibuliform-campanulate, 3-5.5 cm long, mauve to red-violet, darker inside, glabrous with sparsely pubescent greyish hairs on the midpetaline bands; *stamens* 5, unequal, *filaments* longest 7-14mm, shortest 4-6(−7)mm, broadened and slightly hairy at the base, *anthers* obloid, base sagittate, 2.8-3mm long; pollen spinulose, pantoporate; *disc* annular; ovary ovoid, 0.8-1.5mm high, velvety with erect hairs ± 4 mm long; *style* 1, 9-22 mm long, filiform, stigmas 2, globose. *Fruit*: capsule 5-8 mm long hairy, thin walled, crowned by the persistent base of the style, 4-valved. *Seeds* 4, ovoid to globose, 2-4 mm long, blackish-brown, shortly hispid, brown hairs.

DISTRIBUTION – West, east and central Africa. In Guinea: across all four provinces. HABITAT – In Guinea: forest, gallery forest, forest clearing, river edge; found at 126-1400 m.

CONSERVATION STATUS (PRELIMINARY ASSESMENT) – LC (Least Concern), following the global IUCN (2012) guidelines, known from 125 occurrence points worldwide.

SPECIMENS EXAMINED – **Guinea**: Guinée Maritime, Télimélé Préfecture, 5k from Lamba, 4k W of Daramagnaki, 20 Feb. 2011, *Molmou 70* (HNG, K001587952!); Guinée Forestière: Nzérékoré, Gazette, Simandou Range, 25 Jan. 2005, *Laws 49* (K000460114!); *s. loc*., Jan. 1995, *Ditian 16457* (K001587951!); Haute Guinée: Kankan Region: Kankan Préfecture, near to Dabadou, fôret galerie [9° 31’ 31“N 9° 33’ 38”W], 6 Jan. 1967, *Lisowski 90698* (WAG1209261!, POZG-V-0057452).

ADDITIONAL SPECIMENS – **Guinea**: Haute Guinée: 6 km à l’Est de Kankan [10° 23’ 6“N 9° 18’ 20”W], 9 Dec. 1966, *Lisowski 90654* (POZG-V-0057448, POZG-V-0057449, POZG-V-0057450); Moyenne Guinée: Labé Region: Koundara Préfecture, Koundara, 28 Oct. 1958, *Jacques-Georges 13903* (WAG1209263!); Guinée Forestière: Nzérékoré Region: Macenta Préfecture, Sérédou, 25 Jan. 1993, *Lisowski B-7366* (POZG-V-0057446); environs de Sérédou, Ziama [8° 22’ 30“N 9° 17’ 1”W], 17 Jan. 1993, *Lisowski B-7622* (POZG-V-0057447); Sérédou, bord de route [8° 22’ 30“N 9° 17’ 1”W], 20 Jan. 1993, *Lisowski B-7581* (POZG-V-0057451).

### 15.11. *Ipomoea coscinosperma* Hochst. ex Choisy

Type: Sudan Republic, Kordofan, Abu Gerad, *Kotschy 17* (lectotype: G-DC [G00023045!], isotypes: BR [BR0000008885128!], GOET [GOET002509!], HBG [HBG505586!], K [K000097015!, K000097016!], LD [LD1035518!], M [M0109940!, M0109941!], P [P00434153!, P00434154!], TUB [TUB005407!], WAG [WAG0000757!]).

Annual trailing *herb*, glabrescent or hairy; *stems* several, at first suberect but soon prostrate, ± stout, often angular, thinly hairy, glabrescent. *Leaf*: *petiole* 1.8(−3.7)cm long; lamina entire, narrowly elliptic-lanceolate or oblong, 4-11cm long, 1.5-1.9(−6.3)cm wide, rounded cuneate or faintly cordate at the base, blunt but apiculate at the apex, glabrescent, pilose or hairy on both surfaces. *Inflorescence* small, axillary, 1-3-flowered, hispid; *peduncle* <5mm long; *bracteoles* linear-subulate, ±4mm long, margin pilose. *Flower*: sessile; *sepals* subequal, lanceolate or ovate, apex acuminate, 6mm long, 1.8mm wide, margin hyaline, hispid, persistent in fruit; corolla narrowly infundibuliform, slightly longer than the calyx, 5-8 mm long, red or white; pollen spinulose, pantoporate; *disc* annular; *style* 1, filiform, *stigmas* 2, globose. *Fruit*: capsule globose, apiculate, 5-7.5mm in diameter, glabrous, crowned by the persistent style base. *Seeds* 4, brown, very shortly pubescent.

DISTRIBUTION – Tropical and southern Africa. In Guinea: Haute Guinée, in Kankan. HABITAT – In Guinea: found at 379m (elsewhere up to 1560m).

CONSERVATION STATUS (PRELIMINARY ASSESSMENT) – LC (Least Concern), following the global IUCN (2012) guidelines, known from 544 occurrence points worldwide.

SPECIMENS – **Guinea**: Haute Guinée, Kankan Region, Kankan Préfecture, Mont Kankan, savane degrade [10° 23’ 6“N 9° 18’ 20”W], 27 Nov. 1966, *Lisowksi 90672* (POZG-V-0057465).

### 15.12. Ipomoea blepharophylla Hallier f

Type: Sudan, Ghasal-Quellengebiet, grosse Seriba Ghattas, 2[6] May 1869, *Schweinfurth 1818*

(lectotype, designated here: P [P00434136!], isolectotype: K [K000097006!]).

Perennial *herb*, with a small ligneous stump; *stems* several from a woody rootstock, prostrate, up to 1m pubescent with short, dense, spreading, yellowish hairs. *Leaf*: petiole (0.1-)2-9(−15)mm long, pubescent; lamina entire, lanceolate or narrowly oblong, 3.7-8.2(−9)cm long, 0.7-3.2cm wide, rounded to subcordate or rarely cordate at the base, obtuse or acute and mucronate at the apex, glabrous or with odd hairs on the midrib and margins on both surfaces; venation pinnate, ciliate below, 3-4 pairs of secondary veins. *Inflorescence* axillary, 1(or 2)-flowered; peduncle 5-20(−25)mm long, bearing at the top 2 narrowly ovate bracteoles; *bracteoles* unequal length, lanceolate, 3-6mm long. *Flower*: *pedicel* 3-7(−10)mm long, elongating in fruit; *sepals* lanceolate, apex acute, 15-18mm long, 3-4mm wide, persistent in fruit, outer sepals pubescent and ciliate, inner sepals longer, narrowly ovate; corolla infundibuliform, distinctly narrowed below, tube 1.9cm long, 1.5mm wide, up to 5.5cm long, mauve with darker centre, pubescent outside on the mid-petaline bands, with long white hairs; *stamens* 5, included, filaments slightly unequal, broadened and pubescent at the base, *anthers* ovoid, base sagittate, 2mm long; pollen spinulose, pantoporate; *disc* annular; *ovary* ovoid, 9-10mm long, 6-9mm wide, glabrous, surmounted by the base of the style; *style* 1, filiform, stigmas 2, globose. *Fruit*: capsule globose, 9mm in diameter, glabrous, crowned by persistent style-base, 4-valved. *Seeds* 4, 4-4.5 mm long, brown, with greyish hairs, erect.

DISTRIBUTION – Tropical Africa. In Guinea: Haute Guinée. HABITAT – Found at altitudes of 760-1500m, globally.

CONSERVATION STATUS (PRELIMINARY ASSESSMENT) – LC (Least Concern) following the global IUCN (2012) guidelines, known from 162 occurrence points worldwide.

USES – *I. blepharophylla* has medicinal uses in Democratic Republic of Congo as the powered dried root bark is mixed with food to treat malaria. The crushed tuber is given as an enema for children for stomach aches and for amoebiasis in adults. It is a treatment for bloat in cattle (Mwanga Mwanga et al. 2022).

SPECIMENS EXAMINED – **Guinea**: Haute Guinée: Kankan Region: Kankan Préfecture, Kankan, *Jacques Félix 1518,* Jan. 1936 (P03866743!).

### 15.13. Ipomoea eriocarpa R.Br

Type: Australia, New Holland, *Banks & Solander s.n.* (holotype: BM [BM001040629!]).

Very variable annual *herb*; *stem* twining or prostrate, terete, up to 1.5mm in diameter, pilose with yellowish hairs. *Leaf*: *petiole* 1-4.5cm long, villose with yellow-greyish hairs; lamina entire, ovate to linear-oblong, 2.5-8cm long, 0.6-4cm wide, sub-hastate with rounded lobes at the base, long-attenuate to acuminate at the apex, margin pilose-strigose or glabrescent, upper surface glabrescent, more so on the veins, lower surface sparsely pubescent with yellowish hairs especially on the veins; venation pinnate, 5-7 basal veins, 7-11 pairs of secondary veins. *Inflorescence* axillary compact cymes, most commonly 3-many-flowered, very rarely only 1-flowered; sub-sessile or pedunculate, <1 cm long, villose with greyish hairs; bracteoles linear or lanceolate, pilose. *Flower*: *pedicel* ±5 mm long, pilose with greyish hairs; *sepals* subequal, ovate, apex acuminate, basal part 5mm long, 3-4mm wide, apical part 4mm long, 0.5mm wide, villose with greyish hairs, spreading in young fruit.; *corolla* tubular to infundibuliform, 6-9mm long, 13mm wide, mauve, white, pink or white with a mauve centre, limb 1.5cm diameter, pilose on the midpetaline bands; pollen spinulose, pantoporate; *disc* annular; *ovary* long villous; style 1, filiform, stigmas 2, globose. *Fruit*: capsule ovoid-globose to globose, 5-6mm in diameter, pubescent, crowned by the style base. *Seeds* 4, 2.5mm long, black, glabrous, finely punctate.

DISTRIBUTION – Tropical Africa, Asia and Australia. In Guinea: across all four provinces.

HABITAT – In Guinea: Bowal humid zone, agricultural field, brushwood at the edge of agricultural land, roadsides; found at altitudes of 15-460m (elsewhere up to 1,550m).

CONSERVATION STATUS (PRELIMINARY ASSESSMENT) – LC (Least Concern), following the global IUCN (2012) guidelines, known from 3,801 occurrence points worldwide.

USES - The leaves of the plant area eaten as a vegetable, sometimes in soups, in Africa and India. They are dried out in the sun, cooked and served with a staple food (Srivastava & Rauniyar 2020). A root decoction of this plant aids in the fermentation of a local drink ‘kwete’ in Uganda. The seeds are consumed in India. Medicinal uses of this plant include treating menstrual pain, headaches, rheumatism, leprosy, epilepsy, ulcers and fever. *I. eriocarpa* has environmental uses to bind soil, smother weeds and is used as animal food (Mwanga Mwanga et al. 2022).

SPECIMENS EXAMINED – **Guinea**: Haute Guinée: Kankan Region: Kankan Préfecture, à environ 15 km à l’ouest de Kouroussa, sur la route de Dabola, 16 Nov. 2022, *Bidault 6009* (BR, BRLU, G, HNG, K, MO, P, SERG, WAG); Kouroussa Préfecture, 23 Dec. 1901, *Pobéguin 564* (BR0000016788145!, K001594257!); Near to Kankan, next to Milo River, *Lisowski 90673,* 16 Dec. 1966 (BR0000016788121, POZG-V-0057508); Kankan Préfecture, 5km NW of city Kankan, next to Milo River[10° 22’ 51“N 9° 18’ 33”W], 15 Dec. 1966, *Lisowski 90674* (BR0000016788138!, POZG-V-0057510); Moyenne Guinée, Tougé Préfecture, Niavelli, below Niavelli, towards Afia Village, 12 Nov. 2018, *Molmou 1933* (K000875238!); Boké Préfecture, Dougoula Village, SE of Dougoula Village, 15 Dec. 2014, *van der Burgt 1043* (HNG, K001061443!); Bindêlya, Sep. 1901, *Paroisse 33* (K001594256!).

ADDITIONAL SPECIMENS – **Guinea**: Haute Guinée: près de la ville de Kankan, au bord de la Milo [10° 23’ 6“N 9° 18’ 20”W], 12 Dec. 1966, *Lisowski 90668* (POZG-V-0057507, POZG-V-0057508); Guinée Maritime: Conakry, route Conakry – Durbreka, dale latéritique humide [9° 37’ 57“N 13° 35’ 15”W], 27 Dec. 1978, *Lisowski 90309* (POZG-V-0057509); Boke Region: Boké Préfecture, près de Kamsar, rizière humide [10° 39’ 44“N 14° 34’ 45”W], 20 Jan. 1979, *Lisowski 51325* (POZG-V-0057548); Moyenne Guinée: Boké Region: Gaoual Préfecture, Sériba, 29 Oct. 1956, *Adam 13938* (FHI0084914).

### 15.14. *Ipomoea biflora* (L.) Pers

*Convolvulus biflorus* L. in Sp. Pl., ed. 2.: 1668 (1763).

Type: Chine, Hong Kong, haies des jardins à Kennedy-Town, 20 Sept. 1893, *Bodinier 386* [neotype: E00558893; isotype: P (P00622221!)].

*Ipomoea verticillata* Forssk.

Generally annual *herb*; *stem* climbing or prostrate, sometimes with a woody taproot, terete, 2 mm in diameter, often reaching 1-2.5m in height, pubescent to villous with white spreading hairs. *Leaf*: *petiole* 1-9cm long, pilose; lamina entire, oval to oblong-ovate, 2-10cm long, 2-5cm wide, cordate to subhastate at the base, obtuse to acute and mucronate at the apex, margin membranous, lobes pubescent to glabrous on both surfaces; 3-5 basal veins, 4-6 pairs of secondary veins. *Inflorescence* axillary cyme, 1-3-flowered; peduncle 0.5-4.5cm long, thin, hairy to sparsely hirsute; *bracteoles* narrowly oval-elliptical, apex 2-3mm long. *Flower*: *pedicel* 2-4mm long, reflexed in fruit, pilose; *sepals* unequal, 7-8mm long, 5-6mm wide, margin pubescent to glabrous, ciliated, with long and stiff hairs, accrescent in fruit, 3 outer sepals triangular, base cordate to subhastate, apex acute to largely acuminate, 5-8mm long, 2--3mm wide, 2inner sepals smaller, narrowly oval-elliptical; corolla infundibuliform, 1-1.3cm long, 1.5-2cm wide, pale mauve or white, with a purple or violet centre, pubescent on the outside of the midpetaline bands; *stamens* 5, included, *filaments* 2-3mm long, linear, hairy at the base, *anthers* obloid, 0.8-1mm long; pollen spinulose pantoporate; *disc* annular; *ovary* glabrous, 2-celled, 4-ovuled; *style* 1, 4 mm long, filiform, glabrous, stigmas 2, globose. *Fruit:* capsule globose, 5-9 mm in diameter, glabrous, crowned by the accrescent sepals, with apex formed by the persistent style base. *Seeds* 4, 4 mm long, greyish, tomentose.

DISTRIBUTION – Throughout Africa, Arabia, southeast Asia and Australia. In Guinea: majority of the records in Guinée Forestière, one in Haute Guinée.

HABITAT – In Guinea: ruderal area, shrubby savannah; found at 443-580 m.]

CONSERVATION STATUS (PRELIMINARY ASSESSMENT) – LC (Least Concern) following the global IUCN (2012) guidelines, known from 1,875 occurrence points worldwide.

SPECIMENS EXAMINED – **Guinea**: Guinée Forestière: Nzérékoré, Macenta Préfecture, Koenkan, endroit rudéral, 8 Dec. 1962, *Lisowski 90671* (WAG1744966!, POZG-V-0059045).

ADDITIONAL SPECIMENS – **Guinea:** Guinée Forestière: Nzérékoré Region: Macenta Préfecture, Macenta, 11 Nov. 1962, *Lisowski s.n.* (POZG-V-0059048); Haute Guinée: Faranah Region: Faranah Préfecture, Nazwa: Bordou [9° 43’ 34“N 10° 35’ 35”W], 28 Oct. 1966, *Lisowski 90670* (POZG-V-0059046); Guéckédou Préfecture, Guéckédou, bord d’une route [8° 33’ 43“N 10° 7’ 48”W], 17 Dec. 1962, *Lisowski 90666* (POZG-V-0059044); Macenta Préfecture, environs de Sérédou, Ziama [8° 22’ 30“N 9° 17’ 1”W], 23 Jan. 1993, *Lisowski B-7345* (POZG-V-0059043); *loc. cit*., 17 Jan. 1993, *Lisowski B-7613* (POZG-V-0059042, POZG-V-0059050, POZG-V-0059051); près de Macenta, brousse secondaire [8° 32’ 35“N 9° 28’ 7”W], 11 Nov. 1962, *Lisowski 87083* (POZG-V-0059049).

### 15.15. Ipomoea setifera Poir

Type: Guyana, *Brocheton s.n.* [(holotype: P-LAM [P-LAM00357506]; isotype: P-JUSS [P-JUSS6811]).

Trailing or twining herb; stem terete, 1-2 mm in diameter, hirsute with stiff yellowish hairs. *Leaf*: petiole 1-8 cm long, thinner than stem, glabrous but often with scattered tubercles; lamina entire, ovate-deltoid or subreniform, 4-14 cm long, 3-11 cm wide, broadly cordate with wide-spreading obtuse or rounded auricles at the base, obtuse, emarginate and mucronate, less commonly acute or acuminate at the apex, glabrous on both surfaces, lower surface paler; venation reticulate, 3-5 basal veins, 3-5 pairs of secondary veins. *Inflorescence* axillary cymes, 1-3(−5)-flowered; peduncle (0.3-)3-5(−12) cm long, sometimes tuberculate; bracteoles ovate, long mucronate, 1.2-2 cm long, 0.6-1.5 cm wide, persistent, concealing the pedicel bases. *Flower*: pedicel 8-28 mm long; sepals unequal, glabrous, persistent in fruit, outer sepals finely aristate, abaxially 5-winged, wings smooth or often softly tubercled, elliptic, apex acute, 15-22 cm long, 10-15 cm wide, inner sepals ovate, ± 15 mm long, 6 mm wide, pale, unwinged; corolla infundibuliform, 5.5-8 cm long, pink, limb ± 4 cm in diameter, glabrous, unlobed; pollen grain echinate; disc annular; 1 style filiform, 2 stigma globular. *Fruit*: capsule ovoid, 10-12 mm long and wide, often enclosed in slightly accrescent sepals. *Seed* 7-8 mm, minutely pubescent.

DISTRIBUTION – Native from Mexico to tropical America, introduced to west and central tropical Africa. In Guinea: Guinée Maritime, near the coast.

HABITAT – In Guinea: edge of fountain, humid rice field at edge of, mangroves; found at altitudes of 5-100 m (elsewhere up to 550 m).

CONSERVATION STATUS (PRELIMINARY ASSESSMENT) – LC (Least Concern) following the global IUCN (2012) guidelines, known from 637 occurrence points worldwide.

SPECIMENS EXAMINED – **Guinea**: Guinée Maritime: Boké Region: Boffa Préfecture, on road from Conakry to Boffa, 4 Mar. 1949, *Portères s.n.* (P01183742!, P01183743!); Kindia Region, Coyah Préfecture, près de Manéah, rizière humide au bord des mangroves [9° 42’ 41“N 13° 24’ 15”W], 26 Dec. 1978, *Lisowski 51099* (BR0000017391474!, POZG-V-0057946).

ADDITIONAL SPECIMENS – **Guinea**: Guinée Maritime: Boké Region: Boffa Préfecture: Tougnifili, Jugaya, rizière à hydromorphie temporaire [10° 23’ 46“N 14° 25’ 24”W, approximate], 6 Feb. 1979, *Lisowski 51414* (POZG-V-0057947).

### 15.16. Ipomoea quamoclit *L*

Type: Inde, Herb. Clifford: 66, Ipomoea 1 (lectotype: BM [BM000558077]).

Annual *herb*, glabrous; stem twining or prostrate, angular, 1-2 mm in diameter, up to 8m in length, smooth. *Leaf*: petiole 1-3.2 cm long, with 2 subsessile pseudostipules at the base resembling small leaves; lamina lobed, deeply pinnatisect to the main vein, ovate to oblong in outline, 3.5-10 cm long, 3-6 cm wide, lobes opposite to subopposite segments, 10-18 pairs, very narrow to linear or filiform, the lower two reflexed and bifid, glabrous on both surfaces; venation following each lobe. *Inflorescence* axillary cyme, 1-3 flowered; peduncle up to 10 cm long, spindly, with two triangular bracteoles at the top, ± 1 mm long, acute. *Flower*: pedicel 1-20 mm long, slightly clavate; sepals unequal, apex mucronate, persistent in fruit, outer sepals oblong, 4-5 mm long, ± 2 mm wide, inner sepals ovate-elliptic, ± 6 mm long, ± 2 mm wide; corolla salver-shaped, cylindrical tube 2-2.5 cm long, narrowing slightly and gradually towards the base, 2.9-3.5 cm long, bright red, glabrous, with spreading lobes, largely triangular, sharp and mucronate; stamens exserted, filaments equal, threadlike, broadened and hairy at the base, inserted 4-5 mm above the base of the tube, anthers obloid, base sagittate, 1-2 mm long; pollen grain echinate; disc annular; ovary ovoid, 1.5 mm long, 4-celled, 1-ovuled per cell; 1 style 19-30 mm long, filiform, broadened and jointed at the base, 2 stigma globular. *Fruit*: capsule ovoid, 8 mm long, crowned by the base of the style, 4-valved. *Seed* fusiform, blackish, sparsely pubescent.

DISTRIBUTION – Native from Mexico to central America, introduced and invasive in the tropics worldwide. In Guinea: Guinée Maritime, Haute Guinée and Guinée Forestière.

HABITAT – In Guinea: secondary vegetation, near settlement, cultivated; found at 10-950 m (elsewhere up to 1500 m).

CONSERVATION STATUS (PRELIMINARY ASSESSMENT) – LC (Least Concern) following the global IUCN (2012) guidelines, known from 933 occurrence points worldwide.

USES – It is cultivated for ornamental use and is known for attracting hummingbirds (Wood *et al*., 2020) (Mwanga Mwanga *et al*. 2022). It has medicinal uses for haemorrhoids, carbuncles, piles, diabetes, purgative and bleeding from cuts and wounds. The leaf juice is used for body weaknesses including ulcers and chest pain (Srivastava & Rauniyar 2020).

SPECIMENS EXAMINED – **Guinea**: Guinée Maritime: Boké Préfecture, within Dougoula village, 18 Nov. 2007, *Fofana 38* (HNG, K000745537!); Guinée Forestière: Nzérékoré Region, Macenta + Beyla Préfecture, Simandou Range, Canga East or Oueleba, 10 May 2009, *Haba 638* (HNG, K000683679!).

ADDITIONAL SPECIMENS – **Guinea**: Haute Guinée: Kankan Region: Kankan, endroit ruderal, 2 Dec. 1966, *Lisowski 90583* (POZG-V-0057907); Kankan, 15 Dec. 1966, *Lisowski 90590* (POZG-V-0057908); Guinée Forestière: Nzérékoré Region: Macenta [8° 32’ 19“N 9° 28’ 25”W], 6 Dec. 1962, *Lisowski 10631* (POZG-V-0057906); Nzérékoré, 28 Nov. 1962, *Kaden 8* (MW0589685).

### 15.17. Ipomoea alba *L*

Type: Icon in Rheede, Hort. Ind. Malabar 11: t. 50, 1962 (designated by Verdcourt, 1963).

Perennial *herb*, glabrous; *stem* prostrate or twining, terete, striate, 3-4 mm in diameter, up to 2.4m long, mostly smooth, some older stems muriculate, glabrous, ligneous at the base. *Leaf*: *petiole* 5-20cm long, slender, glabrous; lamina entire or shallowly 3-lobed, ovate or orbicular in outline, 6-20cm long, 5-16cm wide, cordate at the base, acute, acuminate or obtuse and mucronulate at the apex, glabrous on both surfaces, acumen covered in small blackish glands on both sides; 7-9 basal veins, 2-3 pairs of arcuate secondary veins. *Inflorescence* axillary, in a united cyme, 1-many-flowered, glabrous; *peduncle* 1-24cm long, cylindrical, glabrous; *bracteoles* small, glabrous, caducous. *Flower*: *pedicel* 0.7-2cm long, lengthening to 2.5-3cm and becoming very thick and clavate in fruit, glabrous; *sepals* unequal, subcoriaceous, glabrous, persistent, often reflexed in fruit, 2-3 outer sepals elliptic, 5-12 mm long, 6-7 mm wide, apex with a long, often curved, awn-like appendage 4-10 mm long, inner sepals longer, suborbicular, apex shortly mucronulate, 8-15 mm long, 9 mm wide; coroll*a* hypocrateriform, tube very slender, 5mm wide and cylindrical to slightly angular, 7-12cm long, 3–8.5(–9)cm wide, white or greenish-cream below, glabrous, opening at night, scented; stamens slightly exserted, glabrous, filaments subequal, 1-3cm long, not broadened at the base, inserted into the upper part of the tube, anthers ovoid to obloid, 4-5mm long; pollen spinulose, pantoporate; *disc* annular; *ovary* obpyriform, glabrous; style 1, 10.5–12.2cm long, filiform, glabrous, stigmas 2, globose. *Fruit*: capsule ovoid, mucronulate, 2.5-3cm long, 1.5-2.3cm wide, glabrous, surrounded by ascending sepals, 2-celled, 4-valved. *Seeds* 4, ovoid, 10-12mm long, 7-9mm wide, white to black, glabrous, smooth.

DISTRIBUTION – Native to tropical America, introduced to Africa, tropical Asia and Pacific. In Guinea: Guinée Forestière.

HABITAT – In Guinea: disturbed habitat, cultivated; found at the altitude of 580 m (elsewhere up to 2800 m).

CONSERVATION STATUS – LC (Least Concern) (Canteiro 2021).

USES – *I. alba* is cultivated for ornamental use and nutritional value (Heine 1963; Meira *et al*. 2012). It has medicinal uses for treating paralysis, swelling, snake bites, constipation, wounds and for its antibacterial, antifungal and cytotoxic properties. The seeds contain small amounts of substances similar to LSD so are used for their hallucinogenic properties, particularly in Afro-Brazilian religious rituals. It is also known to be used in rubber production to improve elasticity (Wood *et al*. 2020; Canteiro 2021).

NOTE – This species is possibly cultivated in Guinea. One specimen [Guinée Maritime: Conakry, *Morvan s.n.*, Jun. 1913 (L2740748)] is of mixed content: *I. alba* (inflorescence and fruit) and *I. mauritiana* (leaves). There is no other record of *I. alba* in Guinée Maritime, suggesting that the occurrence may refer to *I. mauritiana*, to which elements of *I. alba* were mistakenly added.

SPECIMENS EXAMINED – **Guinea**: Koudiafaza, Bindelya: 4 Dec. 1901, *Paroisse 4* (K001591035!).

ADDITIONAL SPECIMENS – **Guinea:** Guinée Forestière: Nzérékoré Region: Macenta Préfecture, Macenta, brousse secondaire [8° 32’ 35“N 9° 28’ 7”W], 18 Dec. 1962, *Lisowski 90665* (POZG-V-0057309); près de Macenta, brousse secondaire [8° 32’ 35“N 9° 28’ 7”W], 30 May 1963, *Lisowski 90657* (POZG-V-0057308).

### 15.18. *Ipomoea muricata* (L.) Jacq

*Convolvulus muricatus* L. in Mant. Pl. 1: 44 (1767).

Type: Inde, Suratt, Braad s.n. in Herb. Linn. No. 218.18 (lectotype: LINN [LINN-HL218-18]).

*Ipomoea turbinata* Lag.

Annual *herb*, glabrous; stem twining, cylindrical to angular, muricate. *Leaf*: petiole 4-12 cm long, smooth or muricate; lamina entire, ovate or orbicular, 7-18 cm long, 6.5-15 cm wide, cordate at the base, acuminate and mucronate at the apex; venation pinnate, 5-9 basal veins, 5-7 pairs of secondary veins. *Inflorescence* axillary cyme, 1-few-flowered; peduncle 3-6 cm long, muricate; bracteoles oblong, apex acute, ± 8 mm long, margin scarious. *Flower*: pedicel 1-2 cm long, becoming thick and reaching 5-7 cm in fruit, clavate, smooth; sepals subequal, ovate to oblong, spreading in fruit, 2 outer sepals 6-7 mm long, with a long awn 4-6 mm long, 3 inner sepals 7-8 mm long with a shorter awn; corolla narrowly infundibuliform to hypocrateriform, 5-7.5 cm long, white to pale blue-purple, glabrous, opening at night; stamens not or slightly exserted, filaments 1.5–1.8(–2) cm, broadened and hairy at the base, inserted in upper part of tube, anthers obloid, base sagittate, 2.8–3.2 long mm; pollen grain echinate; disc annular; ovary glabrous, 4-ovuled; 1 style ± 4.5 cm long, filiform, 2 stigma globular. *Fruit*: capsule ovoid, 1.8-2 cm long, glabrous, surmounted by the persistent base of the style, 2-celled. *Seed* ovoid, flattened, 9-10 mm long, 5 mm wide, brown, pubescent with white hairs.

DISTRIBUTION – Native from Mexico to tropical America, introduced to tropical Africa and Asia. In Guinea: Guinée Forestière, in Macenta Préfecture.

HABITAT – In Guinea: forest edge; also in garbage dumps; found at 560 m (elsewhere up to 1670 m).

CONSERVATION STATUS (PRELIMINARY ASSESSMENT) – LC (Least Concern) following the global IUCN (2012) guidelines, known from 242 occurrence points worldwide.

USES - It is cultivated for ornamental use (Wood et al. 2020; Mwanga Mwanga *et al*. 2022). It is known medicinally for treating skin ailments in the Philippines including chronic and gangrenous wounds, burns and cuts. The crude drug of *I. muricata* is used for the treatment of pharyngitis and otitis externa (Meira et al. 2012).

SPECIMENS – **Guinea**: Guinée Forestière: Nzérékoré Region: Macenta Préfecture, Macenta, Nazwa: Zoubouroumaï, au N de Zoubrouma [8° 32’ 19“N 9° 28’ 25”W], 18 Jan. 1993, *Lisowksi B-7417* (POZG-V-0059007).

### 15.19. Ipomoea verbascoidea Choisy

Type: Angola, 1804, *J.J. da Silva s.n.* (holotype: P [P00150787!]; isotypes: LISC, P [P00150788!]).

*Shrub* or *liana*; stem erect or climbing, subcylindrical, 2-4 mm in diameter, up to 1.5 m tall, longitudinally striated, white or yellowish tomentose when young, becoming pubescent to glabrous with age. *Leaf*: petiole 2-10 cm long, slender, canaliculate, tomentose, with 2 glands at the level of the insertion of the lamina; lamina entire or slightly sinuate, broadly ovate to ovate-orbicular, (3-)7.5-15 cm long, 5-12.5(−14) cm wide, cordate or truncate at the base, obtuse or subacute, shortly apiculate, rarely acuminate at the apex, upper surface green to brownish green, finely pubescent with whitish hairs, lower surface whitish to greyish, with dense whitish hairs; venation pinnate, depressed above, projecting below, 8-10 pairs of secondary veins, the lower 4 subpalmated. *Inflorescence* axillary lax cymes, solitary or in ± 3-flowered cymes; peduncle 1-2.5 cm long, tomentose; 2 bracteoles unequal length, linear-oblong to linear-oblanceolate, 1.3-2 cm long, 5-9 mm wide, tomentose, situated at the base of the pedicel, keeled. *Flower*: pedicel ±1.2 cm long, tomentose like the stem; sepals subequal, elliptic, obtuse, 1-2 cm long, 0.6-1.2 cm wide, coriaceous, margin scarious, tomentose, persistent in fruit, outer sepals 11-16 mm long, 6-8 mm wide, inner sepals 13-17 mm long, (5-)7 mm wide; corolla infundibuliform, tube broadly cylindrical, 6.5-11 cm long, tube 7 cm long, 2.3 cm wide, white, pink or rose-purple with deeper mauve throat, glabrous; stamens ± equal, filaments 6 cm long, with a triangular and pilose base, anthers narrowly obloid, base sagittate, 6-12 mm long; pollen grain echinate; disc annular; ovary ovoid, 2.5-3 mm long, glabrous, 2-celled, 4-ovuled; 1 style 4.6-8(−8.5) cm long, filiform, 2 stigma globular. Fruit: capsule oblong, ovoid or globose, 1.3 cm in diameter, 2-2.5 cm long, glabrous, surrounded ± completely by persistent sepals, 4-valved. *Seed* ovoid, subtrigonal, 6-7 cm long, brown, densely covered with long white or sometimes fulvous cottony hairs, giving the dehisced fruit the appearance of an open cotton ball.

DISTRIBUTION – Tropical and southern Africa. In Guinea: Moyenne Guinée.

HABITAT – In Guinea: found at ca. 700 m (elsewhere up to 1320 m).

CONSERVATION STATUS (PRELIMINARY ASSESSMENT) – LC (Least Concern) following the global IUCN (2012) guidelines, known from 203 occurrence points worldwide.

USES - The tuber of this plant is occasionally consumed as a source of moisture. However, doing so causes a slightly narcotic or paralysing effect leaving the consumer unable to move for a while so it is not considered safe for ingestion (Leffers 2003).

SPECIMENS EXAMINED – **Guinea**: Moyenne Guinée: Mamou Préfecture, Timbo, 16 Jun. 1902, *Pobéguin 1064* (BR0000017395410!).

### 15.20. *Ipomoea carnea* Jacq. *subsp. fistulosa* (Mart. ex Choisy) D.F.Austin

*Ipomoea fistulosa* Mart. ex Choisy in A.P.de Candolle, Prodr. 9: 349 (1845).

Type: Brazil, *Martius 2398* (lectotype: M [M0184890!], isolectotypes: M [M0184891], [M0184892], [M0184894], [M0184889]).

*Undershrub*; *stem* erect or ascending, cylindrical or angular, stout, hollow, 1-2m longitudinally striated, canescent when young, becoming glabrous, the oldest lenticel, glabrous, the youngest subsucculent and puberulous. *Leaf*: *petiole* 2-10(−12)cm long, with 2 glands at the top (extrafloral nectaries); lamina entire, elongate ovate, 10-22cm long, 6-9 cm wide, truncate and slightly cordate at the base, acuminate, obtuse or acute, and mucronate at the apex, puberulous with grey hairs, becoming glabrous on both surfaces; 3-5 basal veins, 8-9(−10) pairs of secondary veins. *Inflorescence* long-pedunculate axillary, somewhat compact cymes, few-many-flowers; *peduncle* 1-12cm long, thick, flattened, puberulous, dichotomously branched; *bracteoles* oval, ± 5mm long, puberulous underneath, deciduous. *Flower*: *pedicel* (0.6-)1.5-2.5cm long, puberulous; *sepals* subequal, suborbicular to largely oval, apex obtuse and marginate, 5-6cm long, (5-)6-7mm wide, margin scarious, puberulous, persistent in fruit; corolla infundibuliform, 5-9 cm long, 5(−10)cm wide, pink to light mauve with a purple throat, limb ± 10cm in diameter, pubescent on the outside; *stamens* 5, unequal, *filaments* longest 2-2.6cm long, shortest 1-1.7cm long, linear, broadened and hairy at the base, anthers obloid, base sagittate, 8-9mm long; *pollen* spinulose, pantoporate; *disc* annular; *ovary* conical, 3-4mm long, pubescent above, 2-4-ovuled; *style* 1 filiform, stigmas 2, globose. *Fruit*: capsule ovoid, mucronate, 15-20mm long, brownish, 4-valved. *Seeds* 4, trigonal, 8-9mm long, black, with long hairs of ± 1 cm, yellowish brown, silky.

DISTRIBUTION – Native to tropical America, introduced to tropical Africa, Asia and Australia.

CONSERVATION – Preliminary assessment of Least Concern following the global IUCN (2012) guidelines, known from 1,034 occurrence points worldwide.

USE – Ornamental (Mwanga Mwanga *et al*. 2022).

OBSERVATIONS – Between the rivers Diani and Nzérekoré (Lisowski, 2009).

NOTES – No specimens were found to document the presence of this species in Guinea; this record is based only on the observations by S. Lisowski (2009).

### 15.21. Ipomoea barteri Baker

Type: Nigeria, Jeba, on the Kworra, 1858, *Barter s.n.* [holotype: K (K000097032!)].

Perennial *herb*; *stem* climbing or prostrate, terete, less than 1mm in diameter, up to 1m, with enlarged storage roots, fusiform or globose, 2-3mm long, 0.8-1.8cm wide, glabrous to scattered fine hairs. *Leaf*: *petiole* (0.1-)0.4-3cm long, usually thicker than the stem, pubescent; lamina entire, narrowly oval to elliptical or oblong-linear, (0.5-)3-14cm long, (0.1-)0.2-2.6(−3)cm wide, cuneate to rounded at the base, obtuse and mucronate at the apex, margin membranous, green or sometimes mottled purple and glabrous to pubescent or hispid on both surfaces; 3 basal veins, 5-7 pairs of secondary veins. *Inflorescence* axillary, 1-2-flowered; peduncle 1-5mm long; *bracteoles* subulate, 1-3 mm long. *Flower*: *pedicel* 5-20mm long; *sepals* equal, oval to orbicular, apex obtuse, 4-5mm long, externally wrinkled, rarely nearly smooth, margin glabrous, persistent in fruit; corolla infundibuliform, 4.5-5.5(−6)cm long, mauve, violet, dark red or white with a coloured centre, glabrous; *stamens* 5, included, *filaments* (3-)4-15mm long, broadened at the base, hairy or glabrous, *anthers* obloid, ± 2 mm long; *pollen* spinulose, pantoporate; *disc* annular; *ovary* ovoid, glabrous, 4-5-celled; *style* 1, filiform, enlarged towards the base, glabrous, *stigmas* 2, globular. *Fruit*: capsule globose, 5-10-seeded, deciduous style. *Seed* ovoid, 3-4mm long, brown, pubescent.

DISTRIBUTION – Widespread in tropical Africa. In Guinea: Moyenne Guinée.

HABITAT – In Guinea: found at 650m (elsewhere up to 1800m).

CONSERVATION STATUS (PRELIMINARY ASSESSMENT) – LC (Least Concern) following the global IUCN (2012) guidelines, known from 149 occurrence points worldwide.

SPECIMENS EXAMINED – **Guinea**: Moyenne Guinée: Fouta Djalon, entre Soumbalako et Boulivel, Sep. 1907, *Chevalier 18640* (P00434132!); Fouta Djalon: between Irebeleya and Timbo, Sep. 1907, *Chevalier 18294* (P00434133!).

### 15.22. Ipomoea pyrophila A.Chev

Type: Mali, Koundian, 17 Feb. 1899, *Chevalier 421* (holotype: P [P00434164]; isotype: P [P00434165]).

Twining *geophyte*; stem terete, 1-2 mm in diameter, woody rhizome, emitting many stems, prostrate or ascending, tomentose with yellow hairs. *Leaf*: petiole 3-25 mm long, curved, less so on older leaves, villose with simple yellowish hairs; lamina entire, oblong-ovate to triangular-lanceolate, base retuse to shallowly cordate, apex obtuse, mucronate, 1-2.5(−6.5) cm long, 1-1.5(−2.5) cm wide, broadly cordate at the base, acute to shortly acuminate, mucronate at the apex, upper surface light green and sparsely pubescent, more dense along the basal veins, with yellowish simple hairs, lower surface green yellowish and sparsely pubescent all over, with yellowish simple hairs; 5 basal veins, 3-4 pairs of secondary veins. *Inflorescence* axillary cymes, 1-3-flowered; peduncle 1-5.2 cm long, tomentose, with simple yellow-whitish hairs; 3 bracteoles narrowly elliptic to oblong, 3-6 mm long, margin long ciliate, villose with simple yellowish hairs, situated at the end of the peduncle, 1-veined. *Flower*: pedicel 2-6(−13) mm long, same hairiness as peduncle; sepals unequal, persistent in fruit, 2 outer sepals slightly narrower at the base, tapering lanceolate, 6-10(−11) mm long, 4-8 mm wide, densely villose on the outside with whitish hairs, 3 inner sepals elliptic lanceolate, apex acute, 6 mm long, 1.5-2 mm wide, glabrous with a stripe of long whitish simple hairs along the back, forming into a long apex; corolla infundibuliform, 2-3.5 cm long, mauve, entirely villous with whitish simple hairs; stamens included, filaments unequal, 2 longest 9 mm long, 3 shortest 7 mm long, broadened and weakly hairy at the base; pollen grain echinate; disc annular; ovary pyriform, contracted at the base, glabrous; 1 style 8-14 mm long, filiform, glabrous, included, 2 stigma globular. Fruit: capsule globose, 5-6 mm in diameter, glabrous. *Seed* obloid, ± 3 mmm long, pubescent.

DISTRIBUTION – West and central Africa: Côte d’Ivoire, Togo, Bénin, Nigeria, Democratic Republic of Congo and Central African Republic. In Guinea: Haute Guinée and Guinée Forestière.

HABITAT – In Guinea: forest, submontane forest, savannah, savannah-woodland, rocky area, river edge.

CONSERVATION STATUS (PRELIMINARY ASSESSMENT) – LC (Least Concern) following the global IUCN (2012) guidelines, known from 60 occurrence points worldwide.

SPECIMENS EXAMINED – **Guinea**: Guinée Forestière: Nzérékoré Region, Macenta + Beyla Préfecture, Simandou Range, adjacent stream cascade, 13 Nov. 2005, *Harvey 296* (HNG, K000460112!); Simandou Range, N. of Pic de Fon, Pic de Dabatini, 1 Dec. 2008, *van der Burgt 1332* (HNG, K000615038!, WAG, SERG); Simandou Range, along road from Pic de Fon, 18 Oct. 2008, *Darbyshire 476* (HNG, K000615039!, P03948550!, MO, WAG); Haute Guinée: Kankan Region, Kérouané Préfecture, Simandou North, Mountain range near Damaro, 11 Feb. 2012, *Simons 845* (K001587971!, WAG1529294!); Mandiana Préfecture, Koundian, 17 Feb. 1899, *Chevalier 421* (K001587970!, P00434165!, P00434164!).

### 15.23. Ipomoea tenuirostris Choisy

Type: Ethiopia: in districtu Memsach prope Genniam, 18 Nov. 1838, *Schimper 1064* (holotype P [P00434221]; isotypes: BR [BR0000008251169], EA [EA000001214], G [G00023038, G00023039], GOET [GOET005705], K [K000097020, K000097021, K000097022], LG [LG0000090028243], M [M0109971], MPU [MPU007056, MPU001521], P [P00434220, P00434219, P00434221], S [S11-40565], STU [STU000322, STU000323], TUB [TUB005460], US [US00664212]).

Perennial *herb*; stem slender, twining or prostrate, sometimes flowering when stem very short, branched, terete, 1-3 mm in diameter, up to 3 m, pilose with yellowish spreading hairs. *Leaf*: petiole 2.5-8 cm long, pilose with greyish hairs; lamina entire, ovate to oblong, 2.2-12 cm long, 0.7-7.5 cm wide, cordate at the base, acute and mucronate at the apex, dry and papery, upper surface sparsely pubescent to glabrous, lower surface puberulous with short greyish hairs; 5-7 basal veins, 4-6 pairs of secondary veins. *Inflorescence* axillary, often lax, sometimes dense cymes, (1 or)2-7-flowered; peduncle 1.5-9 cm long, hirsute with yellowish hairs, secondary peduncle pubescent; bracteoles narrowly oval-elliptical, 2-3 mm long. *Flower*: pedicel 1-3.5 cm long, same hairiness as peduncle; sepals equal, ovate to lanceolate, apex acute, 7-12 mm long, 1.5-3 mm wide, hirsute with yellowish simple hairs, less so on the reverse and margin, persistent in fruit, 2 inner sepals narrower, with hyaline margin; corolla infundibuliform, 2.5-3.8 cm long, 1.5-2.2 cm wide, white to mauve with a violet purple centre, pubescent with greyish hairs on the distinctly purple midpetaline bands; stamens included, filaments unequal, 2-9 mm, broadened and hairy at the base, anthers ovoid-oblong; pollen grain echinate; disc annular; ovary glabrous, 2-celled, 2-ovuled per cell; 1 style 8-10 mm long, filiform, increasingly large at the base, glabrous, 2 stigma globular. Fruit: capsule globose, 6-7 mm in diameter, glabrous, crowned by persistent style base. *Seed* trigonal, 6-8 mm in diameter, brown, long white hairs on the angles, minutely velvety.

DISTRIBUTION – Tropical Africa. In Guinea: Moyenne Guinée and Guinée Forestière.

HABITAT – In Guinea: forest, swamp edge, stream bank forest; found at 1200-1300 m (elsewhere up to 2450 m).

CONSERVATION STATUS (PRELIMINARY ASSESSMENT) – LC (Least Concern) following the global IUCN (2012) guidelines, known from 157 occurrence points worldwide.

USES - This plant has medicinal uses as a natural colic remedy in Democratic Republic of Congo (Mwanga Mwanga *et al*. 2022).

SPECIMENS EXAMINED – **Guinea**: Guinée Forestière: Kankan Region: Kerouané: Simandou nord, crête, bloc C, 7 Nov. 2022, *Bidault 5933* (SERG!, MO!, BRLU!, P!, K!, HNG!); Nzérékoré Préfecture: Mts. Nimba, Gouan Forest, 21 Dec. 2008, *Haba 67* (BR0000005091560!, K001594114!, MO, WAG1744452,); Oueleba main swamp, 16 Nov. 2007, *Cheek 13685* (K000436548!).

ADDITIONAL SPECIMENS – **Guinea**: Moyenne Guinée: Farannah Region: Sarafinian, 1 Feb. 1909, *Chevalier 20638* (P00434216!, P00434217!, P00434218!).

### 15.24. Ipomoea violacea *L*

Type: Icon in Plumier, Codez Boerhaavianus, t. sub n. 851.

Perennial *herb*, glabrous; stem twining or prostrate, angular, 2-3 mm long in diameter, ochraceous, often longitudinally wrinkled but otherwise smooth. *Leaf*: petiole 3.5-9 cm long; lamina entire, circular to ovate, 5-16 cm long, 5-14 cm wide, deeply cordate at the base, acuminate or cuspidate and mucronulate at the apex, glabrous on both surfaces; 5 basal veins, 5-7 pairs of secondary veins. *Inflorescence* axillary, usually solitary, rarely in few-flowered cymes; peduncle 0.75-8 cm long; bracteoles 1-2 mm long, caducous, scale like. *Flower*: pedicel 1.5-3 cm long, becoming much thickened in fruit; sepals subequal, elliptic, apex obtuse, 1.6-2 cm long, margin hyaline, glabrous, reaching 3 cm in fruit; corolla salver-shaped or very narrowly infundibuliform, tube 7-8 cm long, white and/or pale greenish-yellow, limb 2-4 cm long, 5-6 cm wide, glabrous, opening about midnight; stamens included or shortly exserted; pollen grain echinate; disc annular; ovary; 1 style, 2 stigma globular. *Fruit*: capsule globose, 2-3 cm in diameter. *Seed* subtrigonal, ± 1 cm long, black, densely short tomentose and with a narrow ridge of longer hairs ± 3-6 mm long.

DISTRIBUTION – Tropical and subtropical coasts worldwide. In Guinea: Guinée Maritime, near Boké.

HABITAT – In Guinea: thicket bordered by mangrove, near to beach; found at up to 50 m globally.

CONSERVATION STATUS (PRELIMINARY ASSESSMENT) – LC (Least Concern) following the global IUCN (2012) guidelines, known from 1,896 occurrence points worldwide.

USES – This plant has medicinal uses as a diuretic, laxative, expectorant and for cough. The leaves are consumed for headaches and indigestion. The seeds have spiritual uses for communicating with God and have formally been used in sacred Aztec rituals (Srivastava & Rauniyar 2020; Mwanga Mwanga *et al*. 2022).

SPECIMENS EXAMINED – **Guinea:** Guinée Maritime: Boké Préfecture: Kamsar, beach on Taide Island, 26 Nov. 2013, *Guilavogui 682* (HNG, K000749896!).

### 15.25. Ipomoea triloba *L*

Type: Icon in Sloane, Voy. Jamaica 1: t. 97, f. 1 (1707).

Annual *herb*; stem twining, angular, 1-2 mm in diameter, caniculate, glabrous. *Leaf*: petiole 1.2-6 cm long, glabrous; lamina 3-lobed, palmatifid, ovate, 1.5-8 cm long, 1.5-6 cm wide, cordate at the base, acute to acuminate at the apex, glabrous on both surfaces; 7 basal veins, 3-5 pairs of secondary veins. *Inflorescence* axillary cymes; peduncle 3-5 cm long, glabrous or thinly pilose, secondary peduncle 0.2-0.5 cm long; bracteoles filiform, 2-3 mm long, 0.25 mm wide. *Flower*: pedicel 3-7 mm long, glabrous and muricate; sepals subequal, oblong, apex mucronate or caudate, 5-6(−10) mm long, long yellowish hairs on midrib and margins, persistent in fruit; corolla campanulate, 1.5-2(−2.5) cm long, pink, limb 1.3-1.6 cm in diameter, glabrous; pollen grain echinate; disc annular; 1 style filiform, 2 stigma globular. *Fruit*: capsule subglobose, 5-6 mm in diameter, bristly pilose, rarely glabrous. *Seed* 2.8-3 mm long, 2 mm wide, brown, glabrous.

DISTRIBUTION – Native to tropical America, introduced to tropics worldwide. In Guinea: Guinée Forestière and Haute Guinée.

HABITAT – In Guinea: weedy road side, gravel ground; found at 470-500 m. CONSERVATION STATUS – LC (Least Concern) (Contreras & Wood 2019). NOTE – Possibly cultivated in Guinea.

USES - This plant has medicinal properties including antioxidant, antimicrobial, antiviral, antibacterial, antifungal, hypotensive, analgesic, laxative, antimalarial and wound healing (Srivastava & Rauniyar 2020). It is also used in honey production in Cuba and Central America. It has importance due to it being a tertiary genetic relative of sweet potato *I. batatas* and potential gene donor. It can provide drought tolerance characteristics to sweet potato and potential heat tolerance (Contreras & Wood 2019).

SPECIMENS EXAMINED – **Guinea**: Guinée Forestière: Nzérékoré: Near Hotel Nimba, 13 May 2011*, Jongkind 10788* (BR0000017394819!, COI00092364!, K001594200!, WAG1755110!); Haute Guinée: Kankan Region: Kerouané, Simandou North, 12 Nov. 2022, *Bidault 5990* (SERG!, MO!, BRLU!, P!, K!, HNG!); Kankan Region: Kouroussa, 14 Jun. 1902, *Pobéguin 1070* (BR0000017394826!).

### 15.26. *Ipomoea batatas* (L.) Lam

*Convolvulus batatas* L. in Sp. Pl.: 154 (1753).

Type: India, Herb. Linn. (lectotype: LINN 218.12, designated by Verdcourt (1963: 114); iso-lectotypes: LINN 218.13, 218.14).

Perennial *herb* with underground, fusiform to ellipsoid, yellow or reddish edible storage root, colour depending on cultivar; *stem* prostrate, ascending or rarely twining, containing a milky sap, angular in young, 1-2mm in diameter, often rooting at the lower nodes, striate, glabrous or glabrescent. *Leaf*: *petiole* 4-20cm long, glabrous or hairy; lamina entire or 3-5-lobed, palmatifid to palmatisect, triangular to broadly ovate in outline, 4-14cm long, 4-16cm wide, truncate or cordate at the base, acute to acuminate and mucronate at the apex, lobes triangular to lanceolate, glabrous to slightly pubescent on both surfaces, upper surface dark green, lower surface light green; venation pinnate, green or reddish, 3-7 basal veins, 3-7 pairs of secondary veins. *Inflorescence* axillary cymes, 1-several-flowered; *peduncle* 3-18cm long, stout, angular, glabrous or pubescent; *bracteoles* narrow, oblong, acute, 2-3mm long, early deciduous. *Flower*: *pedicel* 3-12 mm long; *sepals* subequal, oblong to elliptical-oblong, apex acute and distinctly mucronate, 7-12mm long, 3-5mm wide, subcoriaceous, glabrous or pilose on the back and fimbriate, persistent in fruit, outer sepals oblong to elliptic-oblong, 7-12mm long, 2-3mm wide, inner sepals elliptic-oblong or ovate oblong, 9-12mm long, 4-5 mm wide; corolla infundibuliform, 3-5cm long, violet or lilac to white with a purple throat, glabrous; *stamens* 5, included, *filaments* unequal, 2 longest 8–14mm long, 3 shortest 6–9mm long, broadened and hairy at the base, *anthers* obloid, base sagittate, 2-3mm long; *pollen* spinulose pantoporate; *disc* annular; *ovary* ovoid, 1.5-2mm long, sparsely pubescent, with long erect hairs, 4-celled; style 1, 15-20mm long, filiform, glabrous, stigmas 2, globose. *Fruit*: capsule ovoid, 8-12mm long, 4-valved. *Seeds* 4, ovoid to irregularly trigonal, 4-7mm long, black, glabrous.

DISTRIBUTION – Native to tropical America, cultivated in all tropical and subtropical regions. In Guinea: Guinée Maritime and Guinée Forestière.

HABITAT – In Guinea: forest, forest clearence, river edge, gallery forest, roadside, cultivated; found at 400-550m.

CONSERVATION STATUS – Data Deficient (DD) (Rowe *et al*. 2019). VERNACULAR NAMES – “sweet potato”.

USES – The tuberous roots of sweet potato are consumed and are among the top ten most important food crops worldwide so have large economical value and are cultivated in almost all tropical and subtropical countries (Wood *et al*. 2020). The roots are eaten boiled, fried or braised. When dried, they can be used to produce flour which is mixed with maize, sorghum and soya flour to produce food for children. The leaves and shoots are also eaten as a vegetable (Mwanga Mwanga *et al*. 2022). This plant has medicinal uses to treat mouth and throat tumours, asthma, bug bites, burns, catarrh, ciguatera, convalescence, diarrhoea, dysclactea, fever, nausea, renosis, splenosis, stomach issues, and whitlows. The leaves can be used as an alterative, aphrodisiac, astringent, bactericide, demulcent, fungicide, laxative and tonic. A variety of white sweet potato is eaten raw to treat anaemia, hypertension and diabetes in Kagawa, Japan (Meira *et al*. 2012).

SPECIMENS EXAMINED – **Guinea**: Guineée Maritime: Kindia Region: Télimélé Préfecture, 14 Nov. 1938, *Chillou 896* (WAG1209077!, WAG1209078!); Guinée Forestière: Nzérékoré Region: Guégédou, 17 Dec. 1962, *Lisowski 91010* (BR0000016066182!, POZG-V-0057385).

### 15.27. Ipomoea sagittifolia Burm.f

Type: Indonesia, Java, *Garcin s.n.* (holotype: G-Burman).

*Ipomoea marginata* (Desr.) Manitz

*Ipomoea sepiaria* J.König ex Roxb.

Perennial *herb*; stem trailing or twining, slender in diameter, yellow brown, with woody tuberous root emitting several stems, furrowed, glabrous or hirsute with long spreading hairs.

*Leaf*: petiole 1-4 cm long; lamina entire, variable, ovate to triangular or narrowly ovate-elliptical, 4-16.5 cm long, 1-15 cm wide, cordate, sagittate, hastate or truncate at the base, acuminate to acute and apiculate at the apex, upper surface glabrous or slightly puberulous, especially towards the edge; 5 basal veins, 3-4 pairs of secondary veins. *Inflorescence* axillary subumbelliform cyme, few-many-flowered; peduncle 2-20 cm long; numerous bracteoles oval to narrowly oval-elliptical, apex obtuse to rounded, 2.5-5(−6) mm long. *Flower*: pedicel (1-)3-6(−10) mm long, glabrous; sepals equal or slightly unequal, widely oval-elliptical to elliptical-oblong, apex obtuse to acute and often mucronate, 4-5(−8) mm long, persistent in fruit, outer sepals shorter, smooth or slightly and sparsely warty, puberulous, inner sepals smooth, glabrous; corolla infundibuliform, 2.7-8 cm long, tube 1.5-4 cm long, 0.3-0.8 cm wide, white to mauve, sometimes purple or violet inside, glabrous; stamens included, filaments unequal, longest 8-10 mm long, shortest 5-6 mm long, broadened and sparsely hairy at the base with glandular hairs, anthers ovoid, base sagittate, apex acute, 2-3 mm long; pollen grain echinate; disc annular; ovary ovoid, 1-2.2 mm long, glabrous, 2-celled; 1 style 1.9-2 cm long, filiform, included or slightly exserted, 2 stigma globular. *Fruit*: capsule globose, apiculate, 5-7 mm in diameter, glabrous, brown. *Seed* subtrigonal, ± 3 mm tomentose.

DISTRIBUTION – Tropical Africa, Asia and Australia. In Guinea: Haute Guinée. HABITAT – Found at 10-1220 m, globally.

CONSERVATION STATUS (PRELIMINARY ASSESSMENT) – LC (Least Concern) following the global IUCN (2012) guidelines, known from 216 occurrence points worldwide.

SPECIMENS EXAMINED – **Guinea**: Haute Guinée: Kankan Region: Kouroussa Préfecture, Kouroussa, 14 Jun. 1904, *Pobéguin 1068* (P03524090!, P03524087!); Dieudion, 24 Mar. 1908, *Pobéguin 1845* (P03524088!).

### 15.28. *Ipomoea nil* (L.) Roth

*Convolvulus nil* L. in Sp. Pl., ed. 2.: 219 (1762).

Type: Icon. in Dillenius, Hort. Eltham 1: 96, t. 80, f. 91 (1732).

Annual herb; stem twining, terete, 1 mm in diameter, pubescent with bristly simple whitish hairs. *Leaf*: petiole 1.5-4 cm long, densely villose with bristly simple whitish hairs; lamina entire or 3-lobed, palmatifid, ovate to circular, 4-14 cm long, 3-13.5 cm wide, cordate at the base, acuminate at the apex, lobes ± acuminate at the apex, middle lobe ovate to oblong, acuminate, lateral ones obliquely ovate to broadly falcate, acuminate, pubescent with adpressed whitish simple hairs on both surfaces, more dense below; 5 basal veins, 4-6 pairs of secondary veins. *Inflorescence* axillary dense cyme, solitary or in lax few-flowered cymes; peduncle 2-10 cm long, densely hirsute with whitish hairs; bracteoles linear to filiform, 5-8 mm long. *Flower*: pedicel 5-10 mm long, same hairiness as peduncle; sepals subequal, linear-lanceolate, apex long-attenuated, 15-28 mm long, 3.5 mm wide at the base, 1.5-2 mm wide above, margin at the base densely pilose with dorsally erect hairs, persistent in fruit; corolla infundibuliform, 5-7.5 cm long, magenta to mauve with paler tube, often white inside, glabrous; stamens included, filaments unequal, longest 20-22 mm long, shortest 12-15 mm long, broadened and pubescent with long hairs at the base, anthers obloid, base sagittate, 3 mm long; pollen grain echinate; disc annular; ovary ovoid, glabrous, 3-celled; 1 style filiform, 2 stigma globular. *Fruit*: capsule ovoid to globose, 0.8-1.2 cm long, glabrous, surmounted by the persistent base of the style, surrounded by the calyx. *Seed* obovoid-trigonal, 4.5-6 mm long, black, puberulous with fine greyish hairs.

DISTRIBUTION – Native to tropical and subtropical America, introduced to Africa, tropical Asia, Australia and the Pacific. In Guinea: Guinée Forestière and Moyenne Guinée.

HABITAT – In Guinea: forest, secondary vegetation, bush edge, submontane forest-grassland transition; found at altitudes of 1140 m (elsewhere up to 2000 m).

CONSERVATION STATUS (PRELIMINARY ASSESSMENT) – LC (Least Concern) following the global IUCN (2012) guidelines, known from 3,247 occurrence points worldwide. NOTE – This species is possibly cultivated in Guinea.

USES - Ornamental use (Mwanga Mwanga *et al*. 2022).

SPECIMENS EXAMINED – **Guinea**: Guinée Forestière: Nzérékoré Region, Macenta + Beyla Préfecture, Simandou Range, Beyla District, 13 Nov. 2005, *Tchiengue 2398* (HNG, K000460105!); Nzérékoré Region: Macenta Préfecture, Near Macenta city, 25 Aug. 1990, *Cordonnier 447* (BR0000016096103!); Moyenne Guinée: Fouta Djalon, 1 Jun. 1963, *Lisowski 91020* (BR0000016096110!).

### 15.29. *Ipomoea indica* (Burm.) Merr

*Convolvulus indicus* Burm. in Auctuarium: 2 verso (1755). Type: Icon. in Besler, Hort. Eyst. Aest. Ord. 8: t. 2 (1613)

Perennial herbaceous *climber*; *stem* twining or prostrate 1-1.5mm in diameter, up to several meters long, rooting at the nodes, pubescent with retrorse hairs, subligneous at the base. *Leaf*: *petiole* 2-10cm long, pubescent; lamina entire or 3-lobed, palmatifid, ovate in outline, 5-12cm long, 3-15cm wide, cordate at the base, acuminate at the apex, lobes acuminate, pilose to glabrescent on both surfaces, more pubescent below; 7-9 basal veins, 2-4 pairs of secondary veins. *Inflorescence* axillary dense cymes, few-many-flowers; *peduncle* 0.5-15cm long; *bracteoles* linear to lanceolate or ovate-lanceolate. *Flower*: *pedicel* 2-10cm long, pubescent like the stem; *sepals* subequal, lanceolate, apex caudate, acuminate, 1.5-2.3cm long, herbaceous, glabrescent, persistent in fruit, outer sepals ovate, apex long acuminate, 9–12mm long, ± pubescent, inner sepals narrower; *corolla* infundibuliform, 5-8cm long, blue or mauve-purple, often red-tinged, tube whitish at the base, limb flaring, glabrous, lobes broadley rounded, notched at apex; *stamens* 5, included, *filaments* unequal, 17–30mm, broadened and hairy at the base, *anthers* obloid, 3.5–5(–5.3)mm long; pollen spinulose, pantoporate; *disc* annular, lobed; *ovary* ovoid, 1-1.5mm long, glabrous, 3-celled; *style* 1, 30-33mm long, filiform, glabrous, *stigmas* 2, globose. *Fruit*: capsule ovoid-globose, 8-10mm in diameter, glabrous, 3-valved. *Seeds* 6, ovoid to ellipsoidal, 4-6mm long, brown-black, covered with an appressed pubescence.

DISTRIBUTION – Native to tropical and subtropical America, introduced to Africa, tropical Asia, Australia and the Pacific. In Guinea: Guinée Maritime and Guinée Forestiére.

HABITAT – In Guinea: cultivated, found at 580 m (elsewhere up to 1500 m).

CONSERVATION STATUS (PRELIMINARY ASSESSMENT) – LC (Least Concern) following the global IUCN (2012) guidelines, known from 5,077 occurrence points worldwide.

USES - This plant is cultivated for ornamental use, and is used medicinally in Hawaii as a purgative for healing broken bones (Wood *et al*. 2020; Mwanga Mwanga *et al*. 2022; Meira *et al*. 2012).

SPECIMENS EXAMINED – **Guinea**: Guineé Maritime: Kindia Préfecture, Fulaya, *Portères s.n.,* 29 Nov. 1961 (P00801285!); Titerekore, Teyèouau, 9 Dec. 1949, *Adam 7327* (BR0000016085169!, WAG1208759!).

ADDITIONAL SPECIMENS – **Guinea:** Guinée Forestière: Nzérékoré Region: Macenta Préfecture, près de Macenta, lisière d’une brousse secondaire [8° 32’ 35“N 9° 28’ 7”W], 1 Jun. 1963, *Lisowski 91020* (POZG-V-0057626); *loc. cit*., 18 Nov. 1962, *Lisowski 65876* (POZG-V-0057625); *loc. cit*., 18 Nov. 1962, *Lisowski 91031* (POZG-V-0057627).

### 15.30. *Ipomoea obscura* (L.) Ker Gawl

*Convolvulus obscurus* L. in Sp. Pl., ed. 2.: 220 (1762).

Type: Icon in Dillenius, Hort. Eltham. 1: 99, t. 83, f. 95 (1832).

*Ipomoea ochracea* G. Don

Perennial *herb*; stem several to many, prostrate to twining, filiform, cylindrical, subligneous at base, 1-2 mm in diameter, up to 3 m long, with a taproot, pilose or glabrescent. *Leaf*: petiole 1-11 cm long, slender, pubescent or glabrescent; lamina entire or slightly undulate, ovate, rarely linear-oblong, 2.5-8.5 cm long, 0.4-7.5 cm wide, cordate with rounded auricles at the base, acuminate or apiculate and mucronate at the apex, margin often ciliate, membranous, pubescent or glabrescent on both surfaces, lower surface paler; venation pinnate, 5-7 basal veins, 2-4 pairs of secondary veins. *Inflorescence* shortly pedunculate axillary cymes, 1-several-flowered, pubescent, 1-4(−5.5) cm long; peduncle 1-8 cm long, slender, glabrous, pubescent or thinly pilose; bracteoles triangular, acute, 1-2 mm long. *Flower*: pedicel 1-2 cm long, at first erect but in fruit relaxed and thickened towards the apex, sometimes minutely verrucose, glabrous or less commonly, pubescent or pilose; sepals subequal, ovate, ovate-orbicular, ovate-lanceolate or lanceolate, apex acute or apiculate, 4-8 mm long, 1.7-4 mm long, often wrinkled or muricate, margin scarious, glabrous or pilose with long white trichomes, in fruit all somewhat accrescent, ultimately often spreading or reflexed, 2 outer sepals shorter, ovate, apex acute to shortly acuminate or mucronate, inner sepals ovate-elliptic, apex obtuse, occasionally mucronate; corolla infundibuliform, 1.5-2.5 cm long, yellow, orange, cream or white, concolorous or with purple centre, often weakly lobed, 3-4 cm diameter, glabrous or the midpetaline areas thinly hairy towards the apices; stamens included, filaments unequal, 2 longer, 3 shorter, broadened and hairy at the base, anthers ovoid, base sagittate, 2-3 mm long; pollen grain echinate; disc annular, 0.5 mm high glabrous; ovary elongated to rounded and distinctly prolonged at apex, 2.8-5 mm long, glabrous, 2-celled, 4-ovuled; 1 style 7-8 mm long, filiform, glabrous, 2 stigma globular, decurrent. *Fruit*: capsule globose-ovoid, 7-12 mm long, 5-10 mm wide, glabrous, crowned by persistent style base. *Seed* ovoid, 4-5.5 mm long, black, appressed pubescent.

DISTRIBUTION – Tropical to southern Africa, tropical America, Asia, Australia and Pacific islands. In Guinea: Moyenne Guinée, Haute Guinée and Guinée Forestière.

HABITAT – In Guinea: gallery forest, montane forest, forest edge, rocky area; found at 529-1570 m (elsewhere up to 1650 m).

CONSERVATION STATUS (PRELIMINARY ASSESSMENT) – LC (Least Concern) following the global IUCN (2012) guidelines, known from 3,420 occurrence points worldwide.

NOTES - Two specimens collected in Guinea of the name *Ipomoea geophiloides* were found in the K herbarium, yet no reference to this name was discovered in the literature. The name *Merremia geophiloides* A. Chev., a synonym of Ipomoea obscura was found, and so it was assumed to be associated to this species. When reassessing the morphological characteristics of the specimens, it was agreed they belonged to the species Ipomea obscura. The examination of one K specimen identified as *Ipomoea tenuirostris* Choisy revealed it did not match the characteristics in the morphological data matrix or the other material of the same species, primarily due to a differing corolla colour. It also turned out to be a specimen of *Ipomoea obscura*, which was correctly re-identified.

SPECIMENS EXAMINED – **Guinea**: Haute Guinée: Kankan Region: Kérouané Préfecture, Simandou nord, forêt galerie à l’est de point zéro, 13 Nov. 2022, *Bidault 6001* (BRLU, HNG, K, MO!, P!, SERG); Mandiana Préfecture, Koundian, 17 Feb. 1899, *Chevalier 416* (P00434195!, P00434196!); Guinée Forestière: Nzérékoré Region: Macenta Préfecture, Koenkan, edge of coffee plantation, 8 Dec. 1962, *Lisowski 90659* (BR0000017551205!, POZG-V-0057822); *loc. cit.* 8 Dec. 1962, *Lisowski 90661* (POZG-V-0057821); Guinée Forestière: Nzérékoré Region: Simandou Range, Pic de Fon, 24 Jan. 2005, *Cheek 12063* (HNG, K000460113!, P); Guinée Forestière: Nzérékoré Region: Peak of Oueleba, 2 Dec. 2008, *Diallo 12* (K000615037!, WAG1755913!); Moyenne Guinée: Faranah Region: Sarafinian, 1 Feb. 1909, *Chevalier 20637* (P00434137!, P00434138!, P00434139!); Mamou Préfecture: Kaba, 10 Jan. 1909, *Chevalier 20388* (P00434140!, P00434141!).

ADDITIONAL SPECIMENS – **Guinea**: Guinée Forestière: Nzérékoré Region: Macenta, 18 Jan. 1993, *Lisowski B-7431* (POZG-V-0057824); Guinée Forestière: Nzérékoré Region: Macenta Préfecture, Seredou, 17 Dec. 1962, *Lisowski 90664* (POZG-V-0057820).

### 15.31. Ipomoea rubens Choisy

Type: India, Sillet, 1829, *Wallich 1421* (lectotype: G [G00227258!]; iso-lectotypes: G00134909!, K001113073!).

Perennial twining *herb*; stem rather woody, finely striate when dry, terete, 1-3 mm in diameter, up to 4 m long, densely short-pilose with soft greyish hairs. *Leaf*: petiole 3-7 cm long, slender, pilose like the stems; lamina entire or shallowly 3-lobed, broadly ovate to circular, 5-15 cm long, 4-12 cm wide, cordate with rounded auricles at the base, acuminate and mucronate at the apex, upper surface pubescent to densely villose, lower surface less densely pubescent to glabrous below with greyish hairs; 7-9 pairs of secondary veins. *Inflorescence* compact, axillary cymes with short branches and consequently flowers subumbellate, 1-few-flowered; peduncle 2-15 cm long, cylindrical, pilose like the stems; bracteoles oval to linear, 3-7 mm long, pubescent, caducous. *Flower*: pedicel 5-17 mm long, pilose; sepals subequal, ovate, apex obtuse or minutely mucronate, 6-8(−11) mm long, 3-6(−10) mm wide, hairy on top but not ciliate, pilose, persistent in fruit, outer sepals accrescent to 16 mm in fruit, ovate-deltoid, apex acute, inner sepals apex obtuse, margin distinctly hyaline; corolla infundibuliform, 4-5 cm long, purple or mauve with darker centre, limb 4-5 cm in diameter, sparsely pilose, midpetaline bands with silky hairs on the outside; stamens included, filaments unequal, ± 3 or 8-9 mm long, filiform, broadened at the base, hairy, anthers obloid, base sagittate, ± 2 mm; pollen grain echinate; disc annular, lobed at the top; ovary glabrous, 2-celled, 4-ovuled; 1 style 15-17 mm long, filiform, included, 2 stigma globular. *Fruit*: capsule globose, 12-13 mm in diameter, glabrous. *Seed* ovoid, 6-9 mm long, 4.3-4.5(−5) mm wide, densely hairy, white or yellowish hairs 2.5 mm long.

DISTRIBUTION – Tropical Africa, Asia and Australia, introduced in tropical and subtropical America. In Guinea: Moyenne Guinée and Haute Guinée.

HABITAT – In Guinea: river edge; found at 1046 m (elsewhere up to 1300 m). CONSERVATION STATUS – LC (Least Concern) (Ghogue 2020).

USES - This plant is known for use as animal food (POWO 2024).

SPECIMENS EXAMINED – **Guinea**: Haute Guinée: Kankan Region: Mandiana Préfecture, Koundion, 17 Feb. 1899, *Chevalier 416* (K001587949!); Kouroussa Préfecture, Morikéniéba, 6 Mar. 1899, *Chevalier 446* (K001587948!); Moyenne Guinée: Labé Region: Route from Labé to Pita, next to river, 15 Feb. 1979, *Lisowski 51497* (BR0000017390200!, POZG-V-0057929).

## DISCUSSION

In Guinea, 51 species of Convolvulaceae are found, of which 38 are native and 13 introduced (Table I). These 51 species represent 16 genera, yet they are very unevenly distributed across these genera, with 63% belonging to *Ipomoea* alone (32 species), and the remaining 19 species are distributed across 15 genera, with most genera represented by a single species, or up to three species. Thus, the majority of these genera only contain native species in Guinea; *Ipomoea* and *Distimake* Raf. contain both native and introduced species, and *Aniseia* Choisy and *Argyreia* Lour. are present only as introduced genera. The most surprising result is the total absence of *Convolvulus*, one the largest genera and most widespread in the family, although with a primarily temperate to sub-tropical distribution, and therefore not expected to occur in this region due to its ecological niche.

**Table 1.**
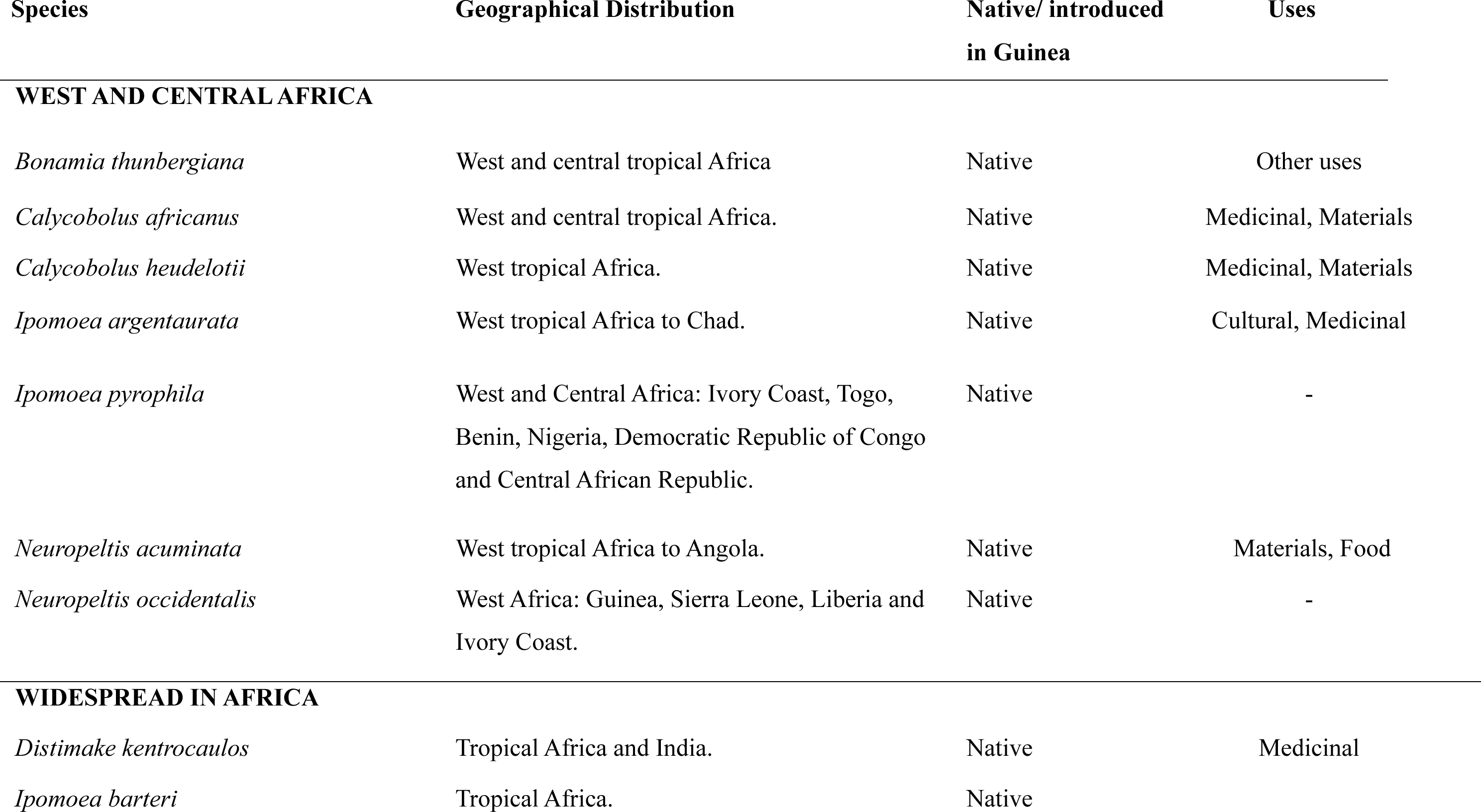
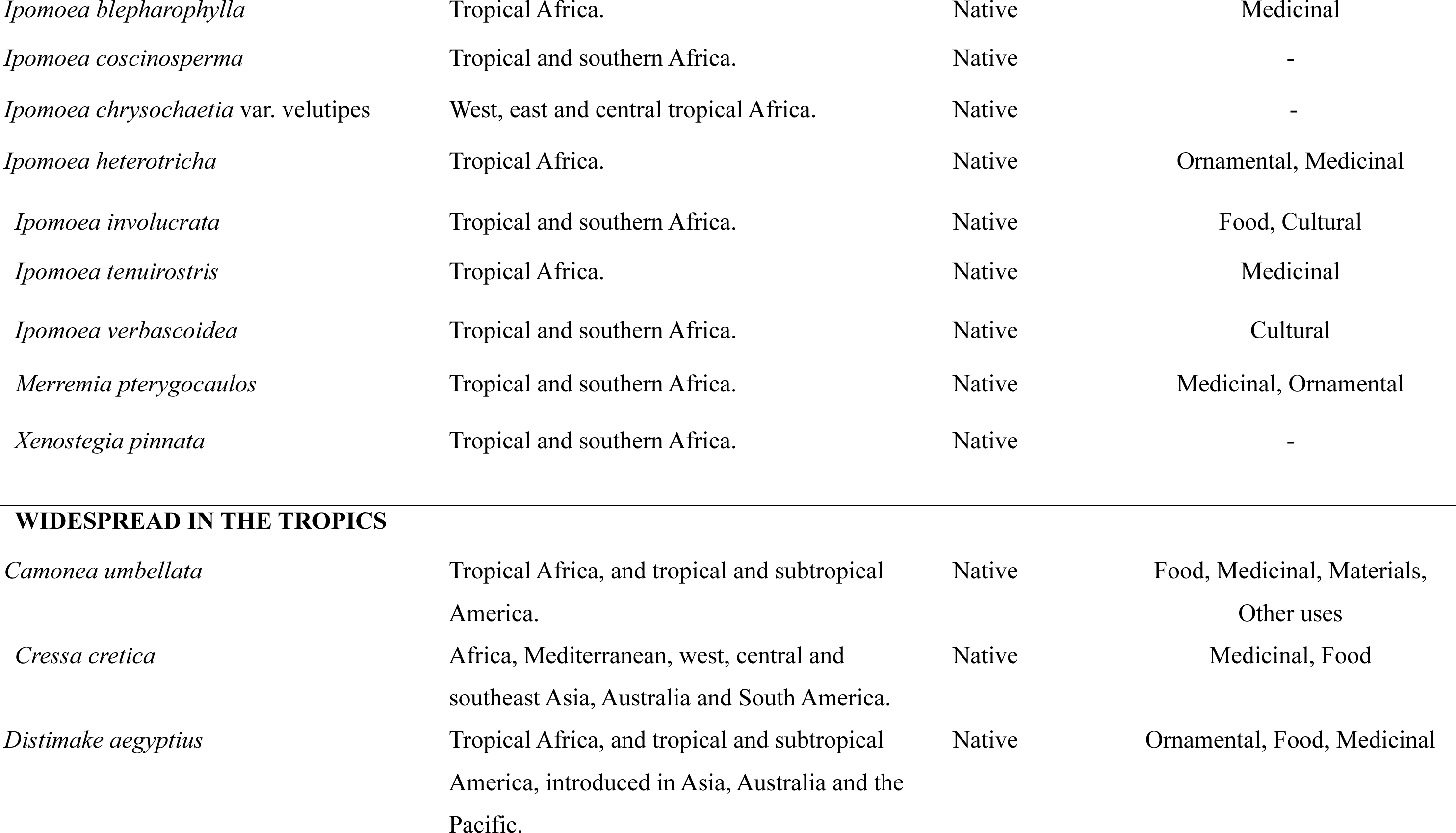
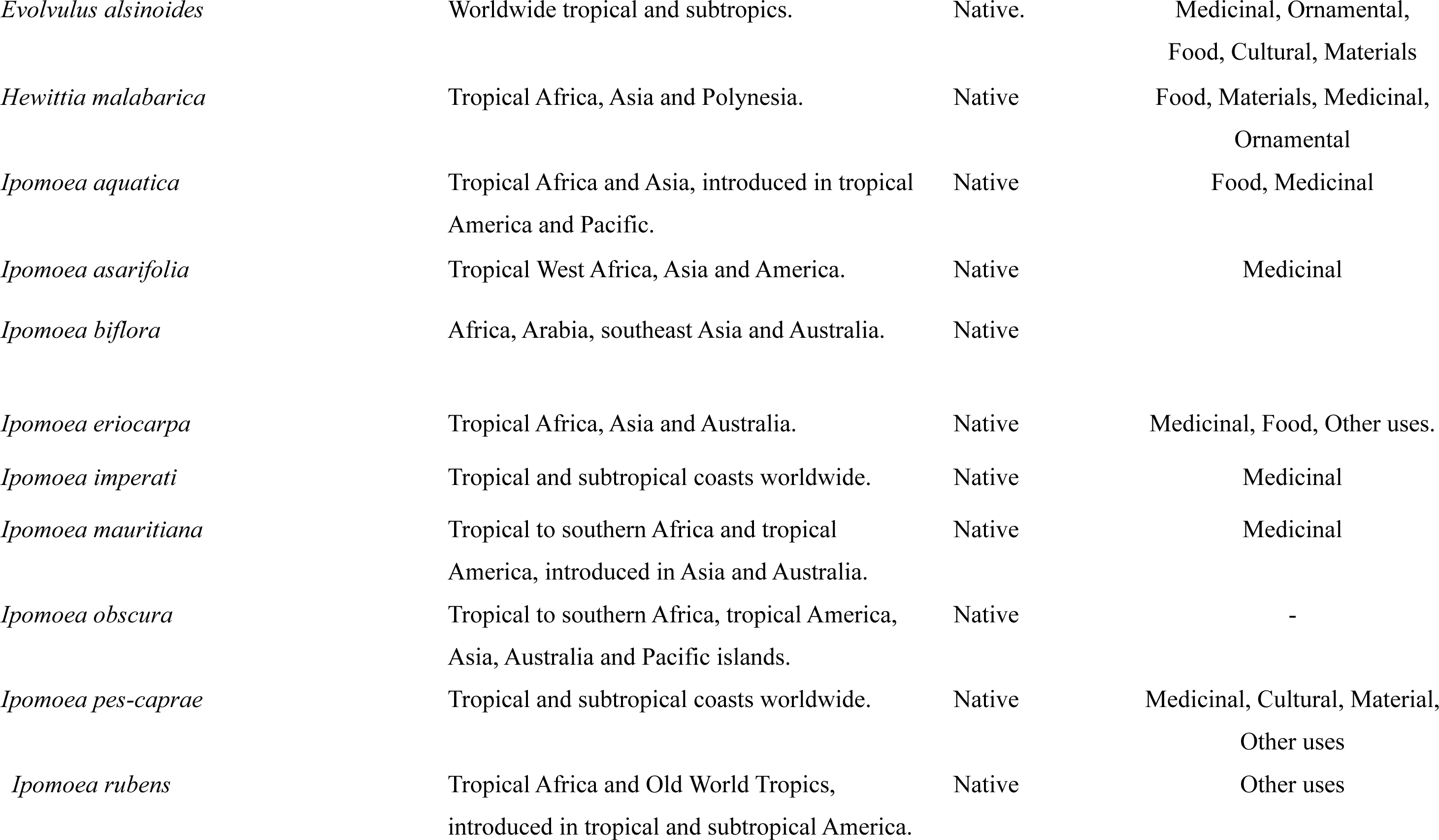
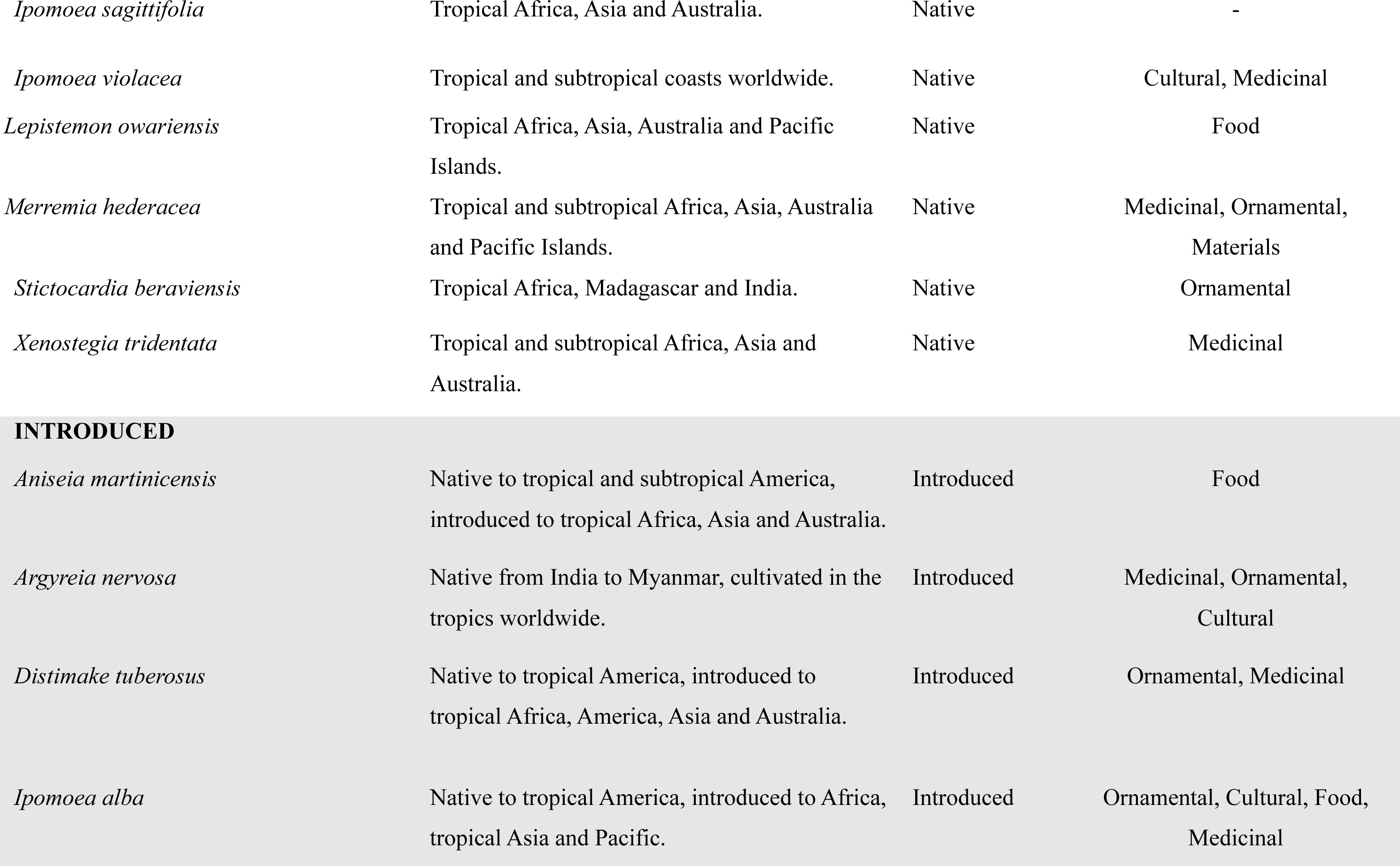
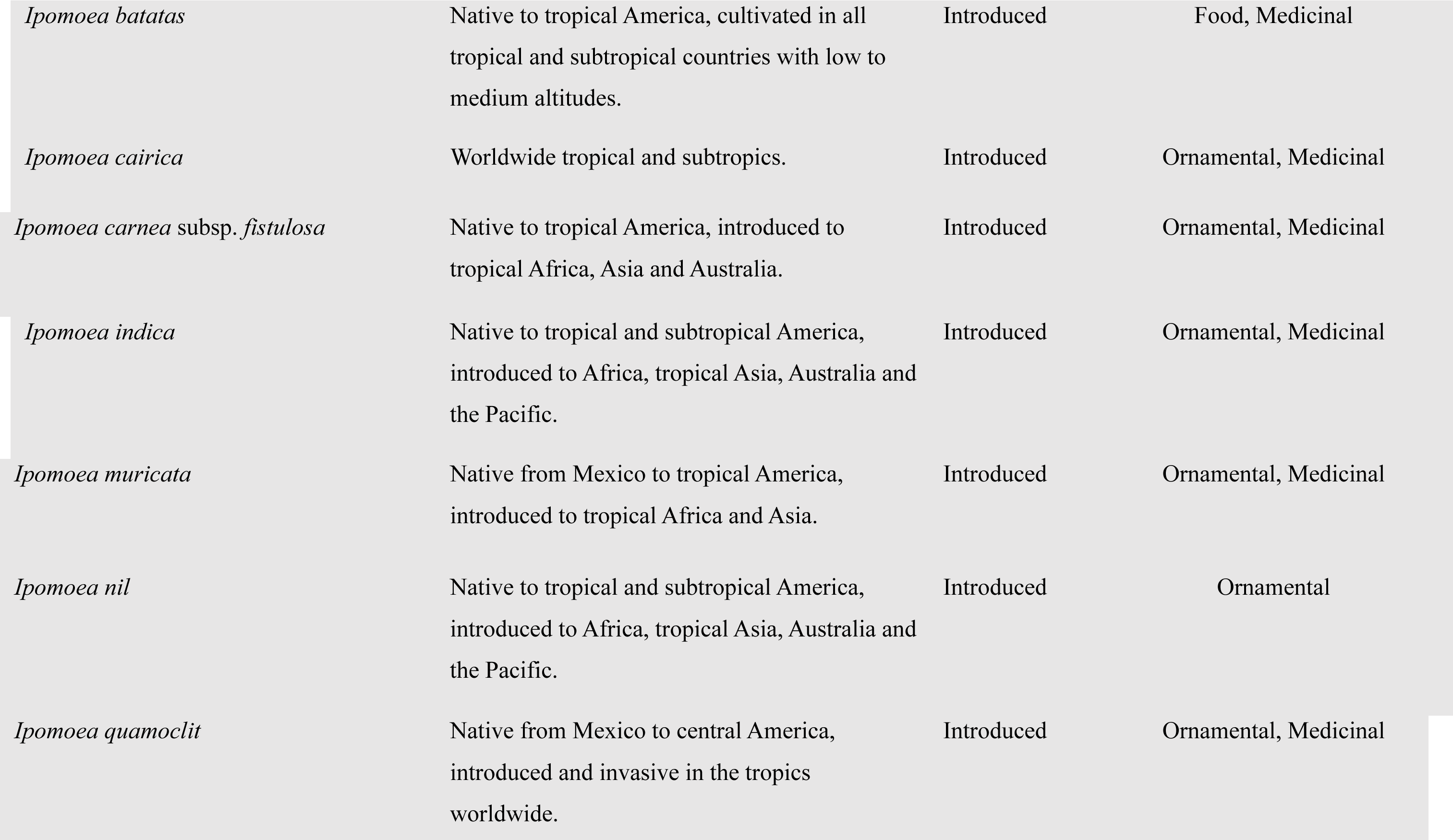
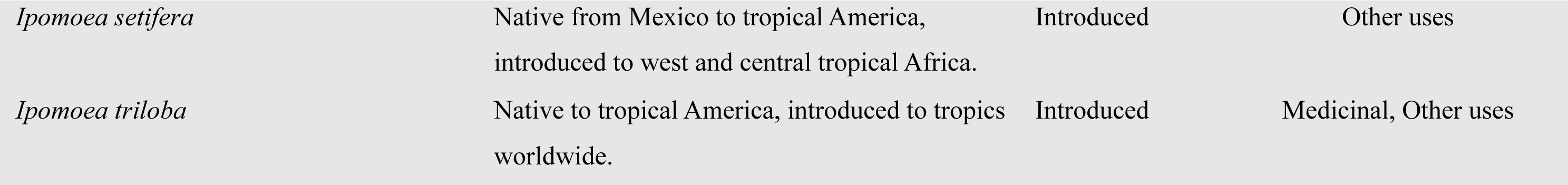
List of species of Convolvulaceae in Guinea, with notes on distribution, origin and traditional uses.

| Species | Geographical Distribution | Native/ introduced<br>in Guinea | Uses |
| --- | --- | --- | --- |
| **WEST AND CENTRAL AFRICA** |  |  |  |
| *Bonamia thunbergiana* | West and central tropical Africa | Native | Other uses |
| *Calycobolus africanus* | West and central tropical Africa. | Native | Medicinal, Materials |
| *Calycobolus heudelotii* | West tropical Africa. | Native | Medicinal, Materials |
| *Ipomoea argenteaurata* | West tropical Africa to Chad. | Native | Cultural, Medicinal |
| *Ipomoea pyrophila* | West and Central Africa: Ivory Coast, Togo, Benin, Nigeria, Democratic Republic of Congo and Central African Republic. | Native | - |
| *Neuropeltis acuminata* | West tropical Africa to Angola. | Native | Materials, Food |
| *Neuropeltis occidentalis* | West Africa: Guinea, Sierra Leone, Liberia and Ivory Coast. | Native | - |
| **WIDESPREAD IN AFRICA** |  |  |  |
| *Distimake kentrocaulos* | Tropical Africa and India. | Native | Medicinal |
| *Ipomoea barteri* | Tropical Africa. | Native |  |
| *Ipomoea blepharophylla* | Tropical Africa. | Native | Medicinal |
| *Ipomoea coscinosperma* | Tropical and southern Africa. | Native | - |
| *Ipomoea chrysochaetia* var. *velutipes* | West, east and central tropical Africa. | Native | - |
| *Ipomoea heterotricha* | Tropical Africa. | Native | Ornamental, Medicinal |
| *Ipomoea involucrata* | Tropical and southern Africa. | Native | Food, Cultural |
| *Ipomoea tenuirostris* | Tropical Africa. | Native | Medicinal |
| *Ipomoea verbascoidea* | Tropical and southern Africa. | Native | Cultural |
| *Merremia pterygocaulos* | Tropical and southern Africa. | Native | Medicinal, Ornamental |
| *Xenostegia pinnata* | Tropical and southern Africa. | Native | - |
| **WIDESPREAD IN THE TROPICS** |  |  |  |
| *Camonea umbellata* | Tropical Africa, and tropical and subtropical America. | Native | Food, Medicinal, Materials,<br>Other uses |
| *Cressa cretica* | Africa, Mediterranean, west, central and southeast Asia, Australia and South America. | Native | Medicinal, Food |
| *Distimake aegyptius* | Tropical Africa, and tropical and subtropical America, introduced in Asia, Australia and the Pacific. | Native | Ornamental, Food, Medicinal |
| *Evolvulus alsinoides* | Worldwide tropical and subtropics. | Native. | Medicinal, Ornamental,<br>Food, Cultural, Materials |
| *Hewittia malabarica* | Tropical Africa, Asia and Polynesia. | Native | Food, Materials, Medicinal,<br>Ornamental |
| *Ipomoea aquatica* | Tropical Africa and Asia, introduced in tropical<br>America and Pacific. | Native | Food, Medicinal |
| *Ipomoea asarifolia* | Tropical West Africa, Asia and America. | Native | Medicinal |
| *Ipomoea biflora* | Africa, Arabia, southeast Asia and Australia. | Native |  |
| *Ipomoea eriocarpa* | Tropical Africa, Asia and Australia. | Native | Medicinal, Food, Other uses. |
| *Ipomoea imperati* | Tropical and subtropical coasts worldwide. | Native | Medicinal |
| *Ipomoea mauritiana* | Tropical to southern Africa and tropical<br>America, introduced in Asia and Australia. | Native | Medicinal |
| *Ipomoea obscura* | Tropical to southern Africa, tropical America,<br>Asia, Australia and Pacific islands. | Native | - |
| *Ipomoea pes-caprae* | Tropical and subtropical coasts worldwide. | Native | Medicinal, Cultural, Material,<br>Other uses |
| *Ipomoea rubens* | Tropical Africa and Old World Tropics,<br>introduced in tropical and subtropical America. | Native | Other uses |
| *Ipomoea sagittifolia* | Tropical Africa, Asia and Australia. | Native | - |
| *Ipomoea violacea* | Tropical and subtropical coasts worldwide. | Native | Cultural, Medicinal |
| *Lepistemon owariensis* | Tropical Africa, Asia, Australia and Pacific Islands. | Native | Food |
| *Merremia hederacea* | Tropical and subtropical Africa, Asia, Australia and Pacific Islands. | Native | Medicinal, Ornamental, Materials |
| *Stictocardia beraviensis* | Tropical Africa, Madagascar and India. | Native | Ornamental |
| *Xenostegia tridentata* | Tropical and subtropical Africa, Asia and Australia. | Native | Medicinal |
| **INTRODUCED** |  |  |  |
| *Aniseia martinicensis* | Native to tropical and subtropical America, introduced to tropical Africa, Asia and Australia. | Introduced | Food |
| *Argyreia nervosa* | Native from India to Myanmar, cultivated in the tropics worldwide. | Introduced | Medicinal, Ornamental, Cultural |
| *Distimake tuberosus* | Native to tropical America, introduced to tropical Africa, America, Asia and Australia. | Introduced | Ornamental, Medicinal |
| *Ipomoea alba* | Native to tropical America, introduced to Africa, tropical Asia and Pacific. | Introduced | Ornamental, Cultural, Food, Medicinal |
| *Ipomoea batatas* | Native to tropical America, cultivated in all tropical and subtropical countries with low to medium altitudes. | Introduced | Food, Medicinal |
| *Ipomoea cairica* | Worldwide tropical and subtropics. | Introduced | Ornamental, Medicinal |
| *Ipomoea carnea* subsp. *fistulosa* | Native to tropical America, introduced to tropical Africa, Asia and Australia. | Introduced | Ornamental, Medicinal |
| *Ipomoea indica* | Native to tropical and subtropical America, introduced to Africa, tropical Asia, Australia and the Pacific. | Introduced | Ornamental, Medicinal |
| *Ipomoea muricata* | Native from Mexico to tropical America, introduced to tropical Africa and Asia. | Introduced | Ornamental, Medicinal |
| *Ipomoea nil* | Native to tropical and subtropical America, introduced to Africa, tropical Asia, Australia and the Pacific. | Introduced | Ornamental |
| *Ipomoea quamoclit* | Native from Mexico to central America, introduced and invasive in the tropics worldwide. | Introduced | Ornamental, Medicinal |
| *Ipomoea setifera* | Native from Mexico to tropical America,<br>introduced to west and central tropical Africa. | Introduced | Other uses |
| *Ipomoea triloba* | Native to tropical America, introduced to tropics<br>worldwide. | Introduced | Medicinal, Other uses |

Previously, the Checklist of Vascular Plants of the Republic of Guinea (Gosline *et al*. 2023) reported 58 taxa of Convolvulaceae for Guinea, including the genus *Cuscuta* which is here not treated, with 3 species present. From the 55 taxa reported in CVPRG, excluding *Cuscuta*, our work has refined this list to 51 taxa, which is due to the following taxonomic changes: 1) *Xenostegia tridentata* subsp*. angustifolia* (Jacq.) Lejoly & Lisowski is here treated as a synonym of *Xenostegia tridentata* (L.) D.F.Austin & Staples; 2) *Ipomoea ochracea* (Lindl.) Sweet is here treated as a synonym of *I. obscura* (L.) Ker Gawl. (Mwanga-Mwanga *et al*. 2022); 3) *Ipomoea chrysochaetia* var. *velutipes* is the only representative of this species in Guinea, and we have established that the typical variety does not occur in the country; 4) *Ipomoea carnea* subsp. *fistulosa* is the only representative of this species in Guinea, and we have established that the typical variety does not occur in the country.

Within Guinea, native species of Convolvulaceae seem to be distributed broadly across the country, with higher frequencies in and around cities along the coast, and in Guinée Forestière near Mount Nimba. This may reflect sampling bias as the coastal cities are easier to reach for collectors and there has been focus on surveying the Mount Nimba region due its high diversity of flora and fauna. Furthermore, continuous conflict in Guinea’s neighbouring countries has meant the border regions are inaccessible for collectors so are often avoided (Arieff 2009). The distribution of introduced species is scattered across the country, showing no particular hotspots. Only 35% of all native Convolvulaceae occurrences fall within protected areas, and many of the areas are still under threat. However, eight of the species have been already assessed as Least Concern on the IUCN Red List, and all 45 species assessed as part of this study are given preliminary Least Concern status.

In addition to the popular use of Convolvulaceae as ornamental plants for their large showy flowers of a diversity of colours (purple, blue, orange) - e.g. morning glories and bindweeds – and the widely appreciated crop sweet potato (*Ipomoea batatas* (L.) Lam.), there is rich traditional knowledge of a much wider range of uses in Guinea, where 83% of the species of Convolvulaceae are traditionally used, as medicine (60%), ornamental (33%), and food (21%). Less represented applications are the use of the root or stem fibres in construction; consumption of the seeds in cultural or religious rituals, for their hallucinogenic properties; and a range of other specific uses. These “other uses” are, however, not neglectable as they are overall present in 27% of the species.

